# Functional validation of allele-specific LMNB1 silencing in patient-derived astrocytes as a therapeutic option for Autosomal Dominant Leukodystrophy

**DOI:** 10.64898/2026.09.09.747273

**Authors:** Martina Lorenzati, Arianna Contato, Fernando Josa-Prado, Marta Ribodino, Maryam K. Ardakani, Ersilia Nicorvo, Elena Signorino, Giacomo Turrini, Valentina Cerrato, Roberta Parolisi, Filippo Lucchini, Gabriele Piacenti, Giacomo Donati, Valentina Proserpio, Juan Antonio Garcìa-Leòn, Marzia Rossato, Luciano Conti, Alfredo Brusco, Giulia Ramazzotti, Stefano Ratti, Pietro Cortelli, Elisa Giorgio, Annalisa Buffo

**Affiliations:** Department of Neuroscience, University of Turin, Italy; Neuroscience Institute Cavalieri Ottolenghi, University of Turin, Italy; Department of Biotechnology, University of Verona, Italy; Department of Life Sciences and Systems Biology, University of Turin, Italy; Departamento Biologia Celular, Genetica y Fisiologia, Instituto de Investigacion Biomedica de Malaga-IBIMA-Plataforma BIONAND, Facultad de Ciencias, Universidad de Malaga, Malaga, Spain; Centro de Investigacion Biomedica en Red sobre Enfermedades Neurodegenerativas (CIBERNED), Madrid, Spain; Department of Cellular, Computational and Integrated Biology, Interdepartmental Center of Medical Sciences, University of Trento, Italy; Cellular Signalling Laboratory, Anatomy Center, Department of Biomedical and Neuromotor Sciences (DIBINEM), University of Bologna, Bologna, Italy; Department of Molecular Medicine, University of Pavia, Italy

**Keywords:** Human astrocytes, leukodystrophy, hiPSC, LMNB1, shRNA, organotypic cultures, RNA therapeutics

## Abstract

Adult-onset Autosomal Dominant Leukodystrophy (ADLD) is a rare fatal leukodystrophy caused by increased LMNB1 gene dosage, most commonly resulting from duplication of the LMNB1 locus. Because ADLD is a gene dosage disorder, selective reduction of pathological LMNB1 expression represents a rational therapeutic strategy. Although allele-specific RNA interference has previously been shown to lower LMNB1 levels in patient-derived fibroblasts and directly reprogrammed neurons, its therapeutic effects have not been evaluated in disease-relevant human glial cells or using functional efficacy endpoints. Here, we established human induced pluripotent stem cell-derived astrocytes from ADLD patients as a human glial model in which to validate allele-specific LMNB1 silencing across molecular, cellular, and functional readouts. ADLD astrocytes recapitulated increased LMNB1 expression and characteristic nuclear abnormalities and displayed transcriptional alterations affecting extracellular matrix organization, calcium homeostasis, metabolism and RNA processing. Functionally, these cells also exhibited functional phenotypes suitable for therapeutic evaluation: astrocyte-conditioned medium impaired the viability of both murine and human oligodendroglial cultures, while conditioned-medium and direct astrocyte-seeding paradigms revealed impaired post-lesion myelin recovery in lysolecithin-treated cerebellar organotypic slices. Allele-specific LMNB1 silencing restored physiological LMNB1 levels, corrected nuclear abnormalities, attenuated astrocyte-mediated oligodendroglial toxicity, improved post-lesion myelin recovery, and was associated with selective transcriptional programs associated with extracellular support and cholesterol metabolism. Together, these findings provide molecular, cellular, and functional validation of allele-specific LMNB1 dosage correction in patient-derived human astrocytes and offer key support for LMNB1-lowering strategies in disease-relevant human glial cells.

## Introduction

Adult Onset Autosomal Dominant Leukodystrophy (ADLD, OMIM #169500) is a progressive and fatal neurological disorder classically characterized by early autonomic symptoms, cognitive impairment, pyramidal tract signs, cerebellar dysfunctions, and symmetric loss of central nervous system (CNS) white matter (Padiath et al. 2006). ADLD is caused by an excessive lamin B1 (LMNB1) production most commonly due to gene duplication (Padiath et al. 2006; Giorgio et al. 2013), less frequently to alterations in the *LMNB1* regulatory landscape (Giorgio et al. 2015; Nmezi et al. 2019). Disease onset typically occurs in the fourth to fifth decade of life with autonomic manifestations preceding pyramidal and cerebellar abnormalities by several years (Coffeen et al. 2000). Neuropathological and magnetic resonance imaging studies have linked increased LMNB1 expression to diffuse myelin loss affecting the frontal and parietal white matter, corticospinal tracts and, depending on the underlying mutation, the cerebellum (Melberg et al. 2006; Quattrocolo et al. 1997; Bergui et al. 1997; Padiath and Fu 2010).

LMNB1 is a core component of the nuclear lamina where it participates in nuclear architecture and multiple other cellular processes (Gerace and Huber 2012; Wong et al 2022). Although the molecular mechanisms linking LMNB1 overexpression to ADLD remain incompletely understood, converging evidence indicates that excessive LMNB1 accumulation perturbs cellular homeostasis and ultimately promotes CNS demyelination (Lin et al. 2013; Heng et al. 2013; Bartoletti-Stella et al. 2015; Rolyan et al. 2015; Giacomini et al. 2016; Nmezi et al. 2025). Because ADLD results from increased LMNB1 gene dosage, reducing LMNB1 expression toward physiological levels represents a rational therapeutic strategy. In this context, an allele-specific RNA interference (ASP-RNAi) approach was previously developed to selectively reduce LMNB1 dosage while preserving physiological expression from the unaffected allele, providing proof of concept for gene-dosage correction as a potential treatment for ADLD (Giorgio et al., 2019). However, this strategy has so far been validated only in patient-derived fibroblasts and directly reprogrammed neurons (Giorgio et al. 2019). Its efficacy therefore remains to be confirmed in disease-relevant glial cell types, including human oligodendroglial cells and astrocytes. Moreover, previous validation studies did not establish whether lowering LMNB1 levels leads to measurable functional benefit.

Oligodendrocytes are considered major cellular targets of the disease. This view is supported by evidence from patients carrying *LMNB1* variants associated with prominent oligodendroglial pathology (Nmezi et al. 2025), as well as by mouse models in which oligodendrocyte-specific LMNB1 overexpression induces severe demyelinating phenotypes, consistent with a cell-autonomous contribution to disease pathogenesis (Heng et al. 2013; Rolyan et al. 2015). However, neuropathological analyses of ADLD brain tissue also reveal prominent astroglial abnormalities, including dysmorphic astrocytes with blunted processes (Dimartino et al. 2024; Alturkustani et al. 2013). Accordingly, forced LMNB1 overexpression in human glioblastoma–astrocytoma cells induces severe alterations in cellular signaling that are only partially rescuable, together with reactive-like features and increased cell death. In contrast, these effects are limited in LMNB1-overexpressing oligodendrocyte-like cells, underscoring the pivotal role of astrocytes in intercellular communication and oligodendrocyte survival. These findings suggest that astrocytes may represent the primary cellular target of LMNB1 overexpression, with astrocytic dysfunction subsequently impairing oligodendrocyte maturation and myelin production (Ratti et al. 2021a, 2021b). Over the years, several models of the disease have been developed to better understand the molecular and cellular mechanisms underlying its pathogenesis (Neri et al. 2023), also studying the role of Lamin B1 in other laminopathies and diseases (Koufi et al. 2023; Evangelisti et al. 2022). Together, these observations suggest that ADLD pathogenesis may also include a non-cell-autonomous component, whereby dysfunctional astrocytes contribute to oligodendroglial impairment and myelin loss, thereby representing a therapeutically relevant cellular target for LMNB1 dosage correction.

To address this hypothesis, we established a patient-derived hiPSC astrocyte model of canonical ADLD, hereafter referred to as ADLD astrocytes. We identified cellular abnormalities that could serve as primary pathological readouts and developed functional assays to uncover disease-associated astrocyte dysfunction. We then used these readouts to assess the therapeutic potential of the ASP-RNAi strategy previously developed to restore physiological LMNB1 dosage (Giorgio et al. 2019). Our findings show that selective silencing of one of the three *LMNB1* alleles through ASP-RNAi normalizes LMNB1 levels and rescues critical cellular and functional defects in ADLD astrocytes and disease-relevant models. These results provide key support for allele-specific *LMNB1* silencing as a promising therapeutic strategy for ADLD.

## Results

### 1. Development and characterization of ADLD disease-relevant cellular models: finding primary and secondary read-outs to be exploited for ASP-RNAi validation

Disease-relevant cellular models are essential for evaluating therapeutic strategies, as they should recapitulate both the primary pathological hallmarks of the disease and functionally relevant phenotypes that can serve as efficacy endpoints. We therefore investigated the cellular, molecular and functional properties of patient-derived ADLD astrocytes to confirm their suitability as a disease-relevant model and to identify robust LMNB1-dependent readouts amenable for validation of our ASP-RNAi strategy.

#### 1.1 Patient-derived ADLD astrocytes recapitulate LMNB1 dosage and nuclear phenotypes suitable for therapeutic correction

Human iPSC lines derived from three ADLD patients carrying the ‘C’ allele of the targeted SNP on the duplicated *LMNB1* copy (Giorgio et al, 2019; Giorgio et al, 2013) and three from healthy subjects (CTRL, Suppl. Figure 1A-C) were used to generate human astroglia based on previously published protocols (Douvaras et al. 2014; Barbar et al. 2020), with minor modifications. The procedure comprised an initial neurulation step followed by suspension culture of glial progenitor spheres, which, upon plating, generated migrating glial cells. CD49f-positive (^+^) astrocytes were isolated from the migratory population by cell sorting and used for downstream analyses (Figure 1A).

**Figure 1.**
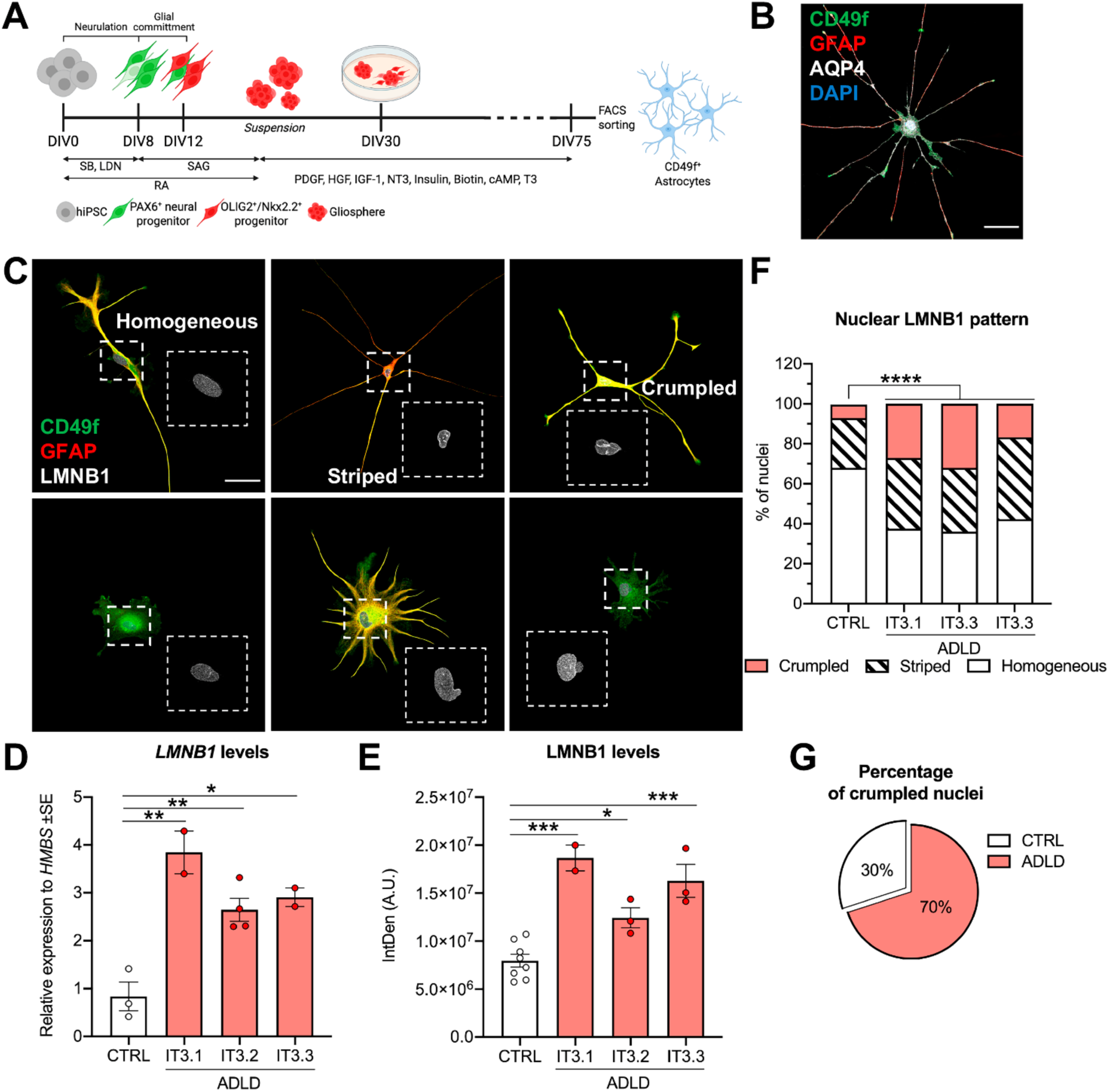
ADLD astrocytes display pathological hallmarks. (A) Schematic representation of the differentiation protocol used to generate human astrocytes from hiPSCs. Created with Biorender. (B) Immunofluorescence staining for the astroglial markers CD49f (green), GFAP (red), and AQP4 (white). (C) Representative astrocytes showing the morphological heterogeneity of these cultures. Distinct nuclear lamina patterns (white) are visualized by anti-LMNB1 staining and classified as homogeneous (left), striped (center), or crumpled (right). CD49f is expressed in all cells (green), with variable co-expression of GFAP (red), confirming astrocyte identity. (D, E) ADLD astrocytes display pathological hallmarks comprising increased levels of *LMNB1* mRNA (D) and protein (E). (F) Quantification of nuclear LMNB1 staining patterns showing nuclear abnormalities. In ADLD astrocytes, crumpled nuclei are more frequent compared to CTRL cells, while homogeneous patterns are reduced. (G) Distribution of all crumpled nuclei scored across CTRL and ADLD cultures. Across all samples analyzed, 70% of the total crumpled nuclei were detected in ADLD astrocytes. CD49f, Integrin alpha-6; GFAP, Glial Fibrillary Acid Protein; AQP4, Aquaporin 4; LMNB1, Lamin B1. Statistics: One-way ANOVA, Tukey’s multiple comparison (D-E; n=2 to 4 independent experiments/cell line; 3 ADLD vs 3 CTRL cell lines); Chi-square Test (F); Fisher’s Exact test (G); *, p<0.05; **, p<0.01; ***, p<0.001; ****, p<0.0001. Scale bar: 50 μm in (B, C).

All lines responded robustly and comparably to gliogenic induction (Suppl. Figure 1D-E) and generated similar proportions of astrocytes (35% CD49f+ cells/total for CTRL cells, 27% CD49f+ cells/total for ADLD cells; Suppl. Figure 1F - dot colors in the graph correspond to the different cell lines). The resulting astrocytes displayed heterogeneous morphologies and expression of typical astroglial markers (Figure 1B, C). Notably, ADLD astrocytes showed a marked increase in LMNB1 expression at both the mRNA and protein levels (about 3- and 2-fold increase, respectively; Figure 1D, E), thereby recapitulating the primary pathological hallmark of the disease *in vitro*.

Increased LMNB1 levels are associated with characteristic nuclear morphological alterations, detectable by anti-LMNB1 staining, which represent established cellular pathological readouts of ADLD (Giorgio et al. 2019). Consistently, ADLD astrocytes exhibited a higher frequency of nuclear abnormalities compared to CTRL cells (Figure 1F, G), with an overall expansion of the proportion of nuclei displaying a crumpled morphology (CTRL: 7%, ADLD: 24%, on average). In line with this enrichment, ADLD astrocytes accounted for 70% of all crumpled nuclei scored (42/60), whereas CTRL cells contributed the remaining 30% (18/60; total nuclei scored: 259 CTRL, 176 ADLD; Figure 1G).

Given prior reports of abnormal astrocyte morphology in post-mortem material (Coffeen et al. 2000; Melberg et al. 2006; Alturkustani et al. 2013; Dimartino et al. 2024) and reactive features in LMNB1-overexpressing glioblastoma-astrocytoma derived astroglial-like cells (Ratti et al. 2021b), we next investigated whether LMNB1 overexpression was associated with overt morphological or neurochemical alterations in ADLD astrocytes. However, although ADLD astrocytes showed a modest tendency toward increased nuclear size, overall cellular extension, soma size, and GFAP and AQP4 levels, none of these differences reached statistical significance indicating no overt morphological or reactive changes (Suppl. Figure 2).

Taken together, these data indicate that under basal culture conditions ADLD hiPSC-derived astrocytes robustly recapitulate LMNB1 overexpression and its associated nuclear alterations, supporting their suitability as a cellular model for validating LMNB1 silencing strategies. differentiation protocol used to generate human astrocytes from hiPSCs. Created with Biorender. (B) Immunofluorescence staining for the astroglial markers CD49f (green), GFAP (red), and AQP4 (white). (C)

Representative astrocytes showing the morphological heterogeneity of these cultures. Distinct nuclear lamina patterns (white) are visualized by anti-LMNB1 staining and classified as homogeneous (left), striped (center), or crumpled (right). CD49f is expressed in all cells (green), with variable co-expression of GFAP (red), confirming astrocyte identity. (D, E) ADLD astrocytes display pathological hallmarks comprising increased levels of *LMNB1* mRNA (D) and protein (E). (F) Quantification of nuclear LMNB1 staining patterns showing nuclear abnormalities. In ADLD astrocytes, crumpled nuclei are more frequent compared to CTRL cells, while homogeneous patterns are reduced. (G) Distribution of all crumpled nuclei scored across CTRL and ADLD cultures. Across all samples analyzed, 70% of the total crumpled nuclei were detected in ADLD astrocytes. CD49f, Integrin alpha-6; GFAP, Glial Fibrillary Acid Protein; AQP4, Aquaporin 4; LMNB1, Lamin B1. Statistics: One-way ANOVA, Tukey’s multiple comparison (D-E; n=2 to 4 independent experiments/cell line; 3 ADLD vs 3 CTRL cell lines); Chi-square Test (F); Fisher’s Exact test (G); *, p<0.05; **, p<0.01; ***, p<0.001; ****, p<0.0001. Scale bar: 50 µm in (B, C).

#### 1.2 Transcriptomic profiling identifies astrocyte pathways suitable for designing LMNB1-dependent functional rescue assays

To further investigate if ADLD astrocytes display molecular alterations that define functional rescue endpoints, we performed a comparative transcriptomic analysis against CTRL astrocytes. Bulk RNA-seq revealed coordinated transcriptional changes affecting gene programs associated with core astrocytic functions.

Differential expression analysis identified 125 differentially expressed genes (DEGs) in ADLD versus CTRL astrocytes, comprising 67 upregulated and 58 downregulated genes (Figure 2A, Suppl. Table 1). Gene Ontology (GO) enrichment analysis (Biological Process category; Suppl. Table 2A) on DEGs revealed extracellular matrix (ECM) organization as one of the most significantly overrepresented terms (Figure 2B). The enriched signal was driven by the upregulation of genes encoding ECM structural components, such as *COL13A1* and *COL4A2*, as well as genes involved in ECM-related regulation and cell-matrix signaling, including *CAV1*, *CAV2*, *ITGA3*, and *PTPRT* (Figure 2C). These data suggest the presence of enhanced ECM remodeling accompanied by altered matrix-dependent signaling, pointing to a reshaping of the extracellular environment through matrix deposition, turnover, and release of ECM-associated bioactive factors. In parallel, downregulated genes were enriched for processes related to altered Calcium Ion Homeostasis, Cytosolic Concentration and Transport (*ADORA2A, GRM5, GRIN2A, GPR35, TRDN, CAMK2B, CAPN3*; Figure 2B, D), consistent with impaired activity-dependent astrocyte communication and calcium signaling, both of which are key regulators of secretory activity. Gene Set Enrichment analysis (GSEA) corroborated these findings by highlighting enrichment of gene sets linked to cell adhesion (“*cadherin binding*” and “*α-catenin binding*”), in line with altered ECM interactions and including *PTPRT* among the core-enriched genes, as well as glutamate-dependent calcium signaling (“*glutamate receptor activity*”), with *GRM5* and *GRIN2A* in the core enriched-genes (Suppl. Table 3). Consistent with these results, altered expression of representative genes from each process, namely *PTPRT* and *GRM5*, was confirmed by qRT-PCR (Suppl. Figure 3A, B).

**Figure 2.**
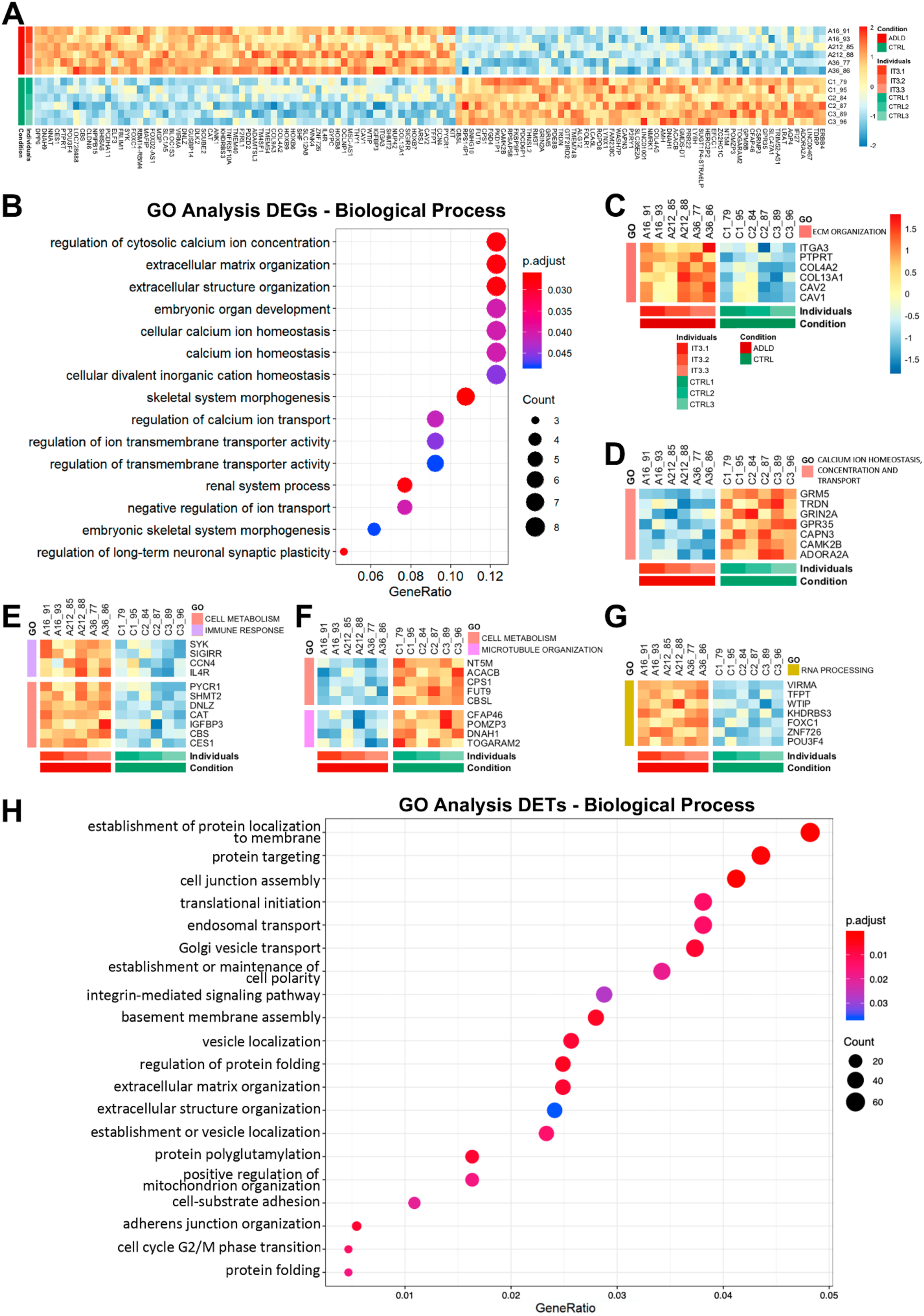
ADLD astrocytes display transcriptional alterations in extracellular matrix, calcium-related signaling, metabolism, microtubule organization and RNA processing. (A) Heatmap of the 125 Differentially Expressed genes (DEGs) in CTRL vs ADLD hiPSC-derived astrocytes (n=2 experimental replicates/cell line; 3 ADLD vs 3 CTRL cell lines). (B-D) GO analysis performed on DEGs (Biological Process) reveals extracellular matrix organization among genes enriched in ADLD astrocytes and regulation of cytosolic calcium ion concentration/homeostasis among genes reduced in ADLD astrocytes. Heatmaps of DEGs involved in these two biological processes are shown in (C) and (D), respectively. (E-G) Heatmaps of selected up and down-regulated genes involved in cellular metabolism and microtubule organization (E, F), and RNA processing (G). DEGs were defined based on p adjusted value ≤ 0.05 and a Log2 Fold Change |log2FC| ≥ 0.5. The labels and color scale shown in (C) apply to panels (D-G). (H) GO Biological Processes enriched among Differentially Expressed Transcripts, comprising extracellular matrix organization and cellular metabolism.

Because GO enrichment may not capture all biologically meaningful changes, we further examined individual DEGs according to their known or predicted functions. This gene-level analysis revealed additional dysregulated processes that complemented the GO-based results (Suppl. Table 2B). This analysis highlighted the upregulation of immune-related genes (*IL4R*, *CCN4*, *SIGIRR*, and *SYK*; Figure 2E), together with downregulation of genes linked to microtubule organization (*TOGARAM2*, *DNAH1*, *POMZP3*, *CFAP46*; Figure 2F). Genes assigned to cellular metabolism showed a mixed pattern, with both upregulated (*CES1*, *CBS*, *IGFBP3*, *CAT*, *DNLZ*, *SHMT2*, *PYCR1*) and downregulated (*CBSL*, *FUT9*, *CPS1*, *ACACB*, *NT5M*) transcripts, indicating metabolic remodeling rather than a unidirectional metabolic shift (Figure 2E, F). Several of these genes are linked to mitochondrial function, suggesting that this metabolic remodeling in ADLD astrocytes includes a mitochondrial component. Consistent with the RNA-seq findings, altered expression of SHMT2, a mitochondrial metabolic gene previously implicated in another hypomyelinating leukodystrophy (Escande-Beillard et al. 2020), was confirmed by qRT-PCR (Suppl. Figure 3C).

Given the reported association between LMNB1 and RNA regulation (Bartoletti-Stella et al. 2015), we next investigated whether ADLD astrocytes show alterations in RNA processing. While we did not detect all genes previously reported as dysregulated in ADLD tissues, our analysis nonetheless revealed upregulation of genes involved in RNA processing (*POU3F4, ZNF726, FOXC1, KHDRBS3, WTIP, TFPT, VIRMA*, Figure 2G). Consistently, transcript-level analysis identified 2757 differentially expressed transcripts (DETs, 1470 upregulated and 1287 downregulated; Suppl. Table 4A) and 4090 differentially alternative spliced genes (DASGs; cumulative across splicing categories) in ADLD astrocytes (Supplementary Figure 3D-F), indicating widespread perturbation of transcript usage and alternative splicing. These findings are in line with previous evidence implicating LMNB1 in RNA processing in other cellular systems (Bartoletti-Stella et al. 2015; Shin et al. 2025), and suggest that similar mechanisms contribute to astrocyte pathology in ADLD. GO biological process analysis of DET-associated genes (Figure 2H; Suppl. Table 4B) further strengthened DEG-based findings identifying ECM organization (*COL4A5, COL13A1, COL4A2, ITGA3, COL9A3*) and cellular metabolism (*SHMT2, CBS*) among the most significantly dysregulated categories in ADLD astrocytes.

Together, these convergent analyses reveal broad disruption of core astrocytic programs in ADLD, comprising extracellular matrix organization, cell adhesion, and glutamate-related signaling. These changes point to altered interactions with the extracellular environment and neighboring cells, providing a rationale for expanding beyond rescue of primary readouts toward assays that test whether defective astrocyte-oligodendrocyte interactions contribute to oligodendroglial dysfunction and are normalized by ASP-RNAi-mediated LMNB1 lowering.

### 2. ASP-RNAi strategy ameliorated ADLD-specific molecular, cellular and functional phenotypes in ADLD astrocytes

We next evaluated the therapeutic potential of the ASP-RNAi approach (Giorgio et al. 2019) across molecular, cellular, and functional disease phenotypes. We first assessed normalization of LMNB1 expression and correction of ADLD-associated nuclear abnormalities as primary readouts. We then tested whether these effects translated into restoration of disease-relevant astrocyte functions and examined the transcriptional programs associated with functional rescue.

ADLD hiPSC-derived mixed glial cultures were transduced with lentiviral particles encoding siRNA-T4 (i.e. targeting the ‘T’ non-duplicated allele; Giorgio et al. 2019), together with a GFP reporter (ADLD shRNA). Lentiviral particles expressing a scrambled shRNA and GFP served as controls. Following transduction, CD49f-positive cells were isolated to enrich for the astroglial population (Materials and Methods; Supplementary Fig. 4A,B). Three days after sorting, ASP-siRNA reduced LMNB1 protein levels by approximately 40%, restoring them to values comparable to those observed in control cells (Supplementary Fig. 4C,D). Importantly, LMNB1 normalization was accompanied by a marked correction of the characteristic ADLD-associated nuclear phenotype, with crumpled nuclei becoming only rarely detectable (Supplementary Fig. 4C).

Together, these findings demonstrate effective target engagement and correction of primary cellular phenotypes through ASP LMNB1 silencing.

#### 2.1 ASP-RNAi mediated LMNB1 lowering rescues astrocyte-mediated oligodendroglial toxicity in murine and human cultures

We next tested whether restoring physiological LMNB1 levels in ADLD astrocytes could also rescue disease-relevant functional alterations. Guided by the transcriptomic changes, we first assessed whether soluble factors released by ADLD astrocytes impaired oligodendroglial viability, thereby establishing a functional readout for astrocyte-targeted LMNB1 silencing. To this aim, we collected astrocyte-conditioned medium (ACM) from high-density ADLD or CTRL astrocyte cultures and applied it during maturation to murine oligodendroglial cultures (mOL), comprising both oligodendrocyte precursor cells (OPCs) and their differentiating progeny (Figure 3A). ADLD ACM significantly increased oligodendroglial cell death, as reflected by a higher number of Cytotox^+^ cells per field relative to CTRL ACM (Figure 3B, C). Cytotox^+^ profiles often displayed the morphology of apoptotic bodies (Figure 3B, white arrows), indicating that soluble factors released by ADLD astrocytes exert a toxic effect on murine oligodendroglial cells.

**Figure 3.**
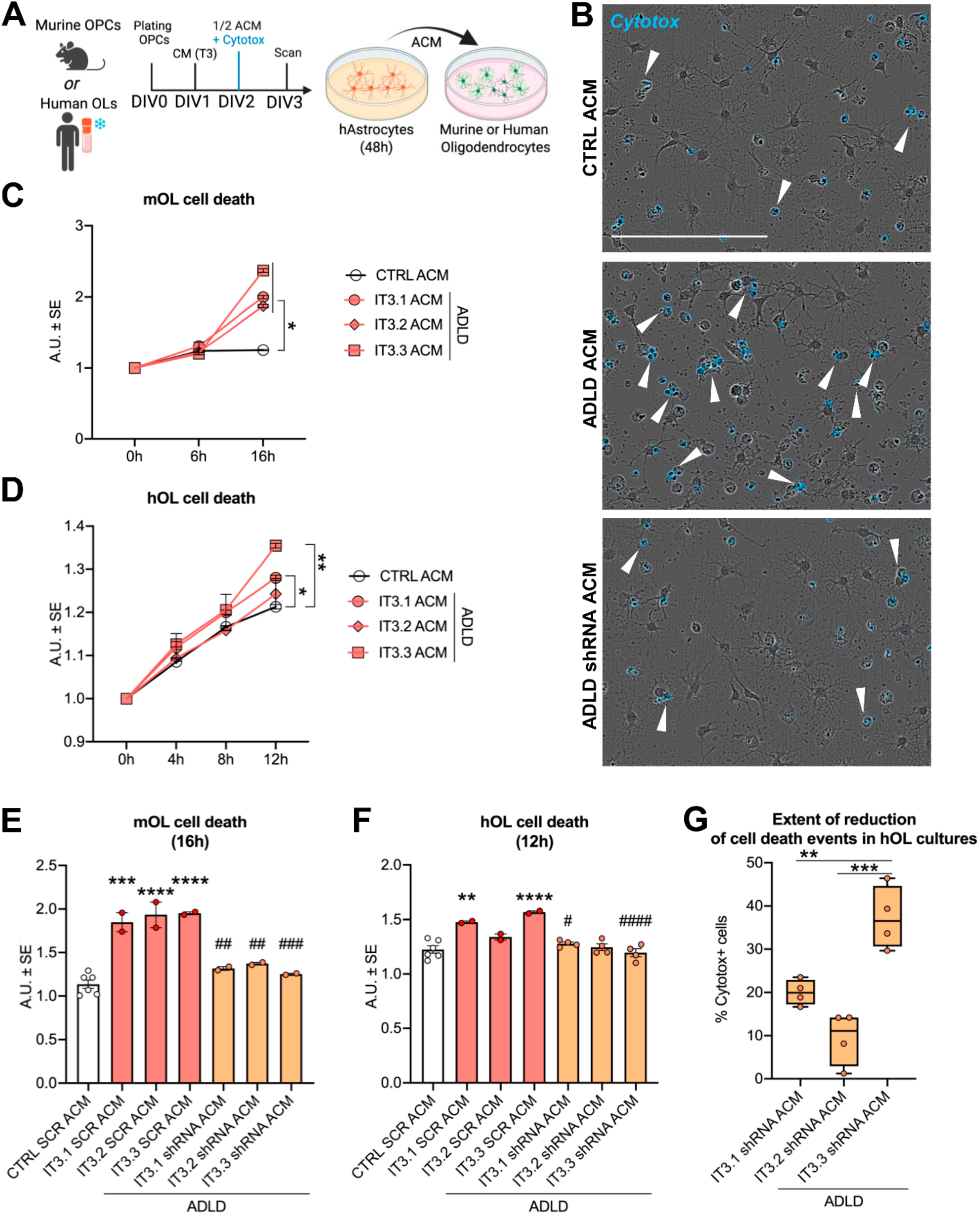
ADLD astrocyte-conditioned medium decreases the viability of murine and human oligodendroglial cells *in vitro,* and LMNB1 lowering attenuates this effect. (A) Schematic representation of the experimental protocol used to test if ADLD ACM compromises oligodendroglial viability through paracrine mechanisms. Created with Biorender. (B) Representative images of mOL cultures 16h after the start of the culture with CTRL ACM (left), ADLD ACM (center) or ADLD shRNA ACM (right). Cyan staining indicates dying Cytotox^+^ cells, white arrows indicate apoptotic-body-like profiles. (C, D) Quantifications of OL cell death over time, measured by Cytotox NIR Object counts (Incucyte), in mOL (C) and hOL (D) cultures treated with CTRL, ADLD, or ADLD shRNA ACM. Values are expressed relative to the “0h” time point for each condition. (E, F) Cell-death readout in murine (E) and human (F) oligodendroglia 16h and 12h, respectively, after the application of CTRL/ADLD SCR or IT3.1/2/3 shRNA ACM. (G) Percentage of reduction of cell death in hOL cultures treated with shRNA compared with the corresponding SCR ACM conditions. ACM, Astrocyte Conditioned Medium; mOL, murine oligodendrocytes; hOL, human oligodendrocytes; SCR, scramble Statistics: Two-way ANOVA, Sidak’s multiple comparison (C, D; n= 2-4 experiments/condition; 3 ADLD vs 2 CTRL cell lines); One-way ANOVA, Tukey’s multiple comparison (E, F, G; n=2-4 experiments/condition). IT3.1/2/3 SCR ACM vs CTRL SCR ACM: *, p<0.05; **, p<0.01; ***, p<0.001; ****, p<0.0001. IT3.1/2/3 shRNA ACM vs ADLD SCR ACM: #, p<0.05; ##, p<0.01; ### p<0.001; #### p<0.0001. CTRL SCR ACM vs IT3.1/2/3 shRNA ACM: ns. Scale bar: 200 µm in (B)

We then asked whether this cytotoxic activity extends to induced human oligodendroglia (hOL) (García-León et al. 2020) by treating these cells with CTRL or ADLD ACM (Figure 3D). In line with results in murine cells, ADLD ACM increased the proportion of Cytotox^+^ hOLs, although the effect was less pronounced overall. Notably, whereas murine oligodendroglial cultures exhibited relatively consistent responses across ACM from different ADLD donors, hOLs displayed greater donor-to-donor variability, with IT3.3 ACM eliciting the strongest cytotoxic response, IT3.1 showing a milder trend in the same direction, and IT3.2 failing to reach statistical significance.

Collectively, these data show that soluble factors released by ADLD astrocytes impair oligodendroglial viability through contact-independent mechanisms. Despite a more uniform response in murine cultures and greater heterogeneity in human oligodendroglia, this phenotype provided a disease-relevant readout for testing the functional outcome of LMNB1 normalization. We therefore collected the astrocyte-conditioned medium from high-density cultures transduced with either SCR or LMNB1-targeting shRNA and applied it to oligodendroglial cultures as described above.

The toxic activity of ADLD ACM was attenuated after LMNB1 normalization in both murine and human systems (Figure 3E-G). In murine oligodendroglia, cell death was significantly reduced compared with the SCR condition (ADLD shRNA ACM vs ADLD SCR ACM; Figure 3E), corresponding to an approximately 30% decrease in mOL cell death and approaching levels observed with control donor ACM (Figure 3E). In human hOLs (Figure 3F), the most pronounced toxicity attenuation was observed with IT3.3 shRNA ACM (approximately 36% reduction in Cytotox^+^ cells relative to IT3.3 SCR ACM), whereas smaller effects were observed with IT3.1 shRNA ACM (∼20% vs IT3.1 SCR ACM) and IT3.2 shRNA ACM (∼11% vs IT3.2 SCR ACM) (Figure 3F, G). Overall, these findings indicate that restoring astrocytic LMNB1 dosage toward physiological levels reduces the detrimental effects of ADLD ACM. (E, F) Cell-death readout in murine (E) and human (F) oligodendroglia 16h and 12h, respectively, after the application of CTRL/ADLD SCR or IT3.1/2/3 shRNA ACM. (G) Percentage of reduction of cell death in hOL cultures treated with shRNA compared with the corresponding SCR ACM conditions. ACM, Astrocyte Conditioned Medium; mOL, murine oligodendrocytes; hOL, human oligodendrocytes; SCR, scramble Statistics: Two-way ANOVA, Sidak’s multiple comparison (C, D; n= 2-4 experiments/condition; 3 ADLD vs 2 CTRL cell lines); One-way ANOVA, Tukey’s multiple comparison (E, F, G; n=2-4 experiments/condition). IT3.1/2/3 SCR ACM vs CTRL SCR ACM: *, p<0.05; **, p<0.01; ***, p<0.001; ****, p<0.0001. IT3.1/2/3 shRNA ACM vs ADLD SCR ACM: #, p<0.05; ##, p<0.01; ### p<0.001; #### p<0.0001. CTRL SCR ACM vs IT3.1/2/3 shRNA ACM: ns. Scale bar: 200 µm in (B).

#### 2.2 ASP-RNAi-mediated LMNB1 lowering restores astrocyte-dependent myelin recovery in an *ex vivo* demyelination model

Having established that ADLD astrocytes impair oligodendroglial viability through soluble factors and that normalization of astrocytic LMNB1 dosage mitigates this effect, we next investigated whether LMNB1 correction could rescue a more complex disease-relevant phenotype, namely impaired myelin repair. Because ADLD is characterized by progressive white matter loss and limited myelin restoration, we asked whether ADLD astrocytes contribute to defective post-lesion remyelination and whether this phenotype can be improved by restoring physiological LMNB1 levels.

IT3.3 was selected as the reference pathological cell line for these experiments because it showed the strongest *LMNB1* dosage-dependent effect on ACM-mediated human oligodendrocyte survival *in vitro*. To model myelin injury and recovery, cerebellar slices were exposed to LPC to obtain a demyelination-remyelination model. In this model, LPC exposure reduced the fraction of calbindin (CALB) positive (^+^) Purkinje axonal profiles associated with MBP⁺ structures (MBP^+^/CALB^+^ axons), consistent with myelin damage. This fraction subsequently increased during spontaneous recovery (SR; Supplementary Figure 5A), supporting its use as a readout of post-lesion myelin restoration (see Supplementary Figure 5B, C for a detailed example of MBP^+^/CALB^+^ axon). Although this measure did not specifically identify newly formed myelin segments, the temporal dynamics of this metric justified its use as a proxy for myelin loss and recovery. After the peak of demyelination, a subset of slices was exposed to ACM derived from high-density CTRL SCR, IT3.3 SCR, or IT3.3 shRNA astrocyte cultures and analyzed after 2 days of treatment (Figure 4A). Representative whole-slice images of the different experimental conditions are shown in Supplementary Figure 5D (panels I-IV). Slices exposed to IT3.3 SCR ACM showed a significantly reduced MBP⁺/CALB⁺ axon fraction compared with both CTRL SCR and SR conditions, which did not differ from each other (Figure 4B, C). By contrast, IT3.3 shRNA ACM significantly increased the MBP⁺/CALB⁺ axon fraction relative to IT3.3 SCR ACM, reaching values above control (Figure 4B, C).

**Figure 4.**
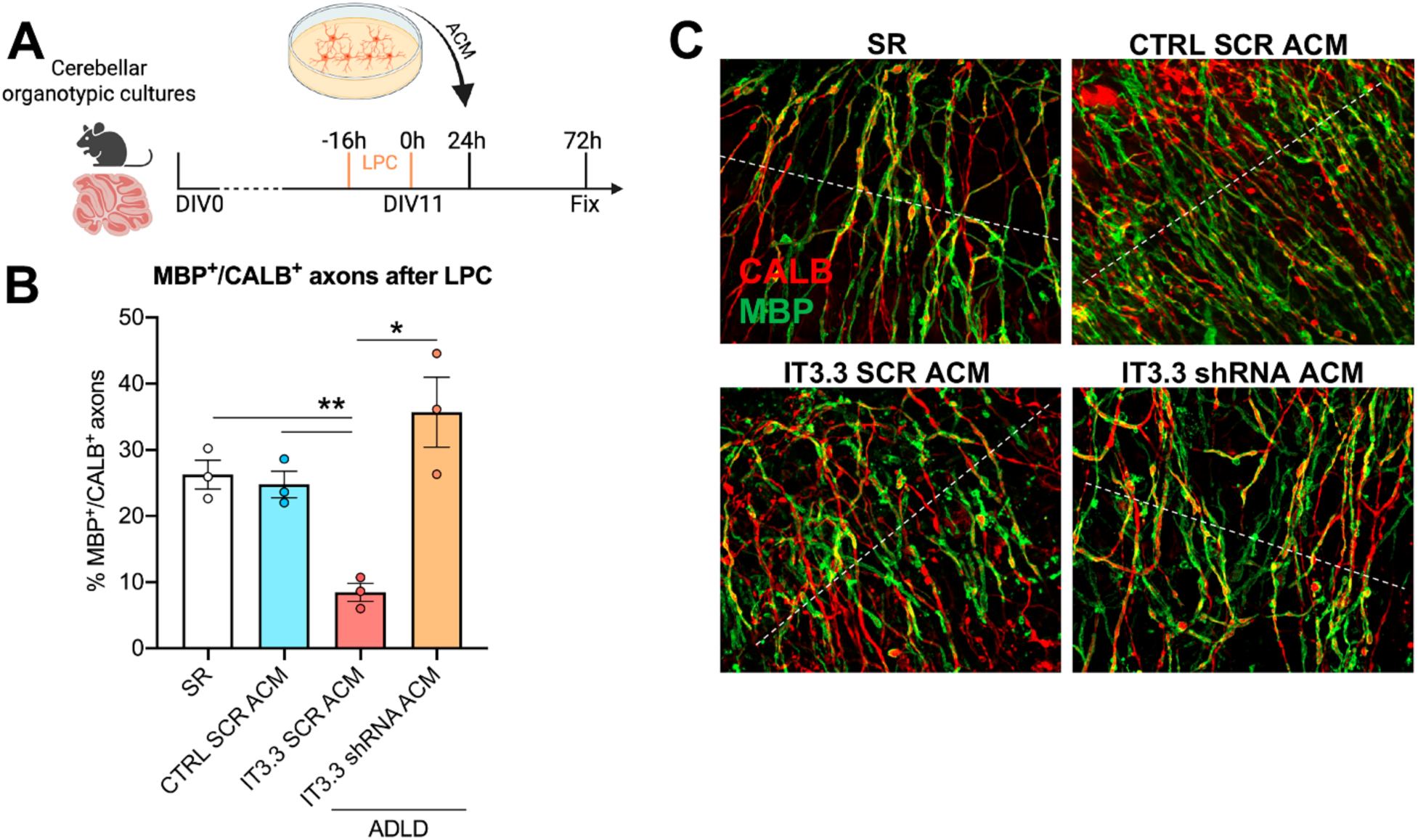
LMNB1 lowering restores the inhibitory effect of ADLD astrocyte-conditioned medium on post-lesion myelin recovery *ex vivo*. (A) Schematic representation of the experimental protocol used to test if ADLD ACM compromises the remyelination process on organotypic LPC-demyelinated murine cerebellar slices. Timing of the LPC demyelinating agent and ADLD ACM application are shown. Created with Biorender. (B) Fractions of MBP^+/^CALB^+^ axonal profiles, indicative of remyelinated axons, in CTRL SCR, IT3.3 SCR and shRNA conditions. Values are expressed relative to the condition of spontaneous recovery (SR) at 72h post LPC. (C) Representative images of cerebellar Purkinje neuron axons in organotypic cultures stained for CALB (red) and MBP (green) in CTRL SCR, IT3.3 SCR and shRNA conditions. A white dotted line perpendicular to the predominant orientation of CALB^+^ axons is shown as an example of the used approach (see also **Suppl. Figure 5B, C**). LPC, Lysolecithin; ACM, Astrocyte Conditioned Medium; CALB, Calbindin; MBP, Myelin Basic Protein. Statistics: RM One-way ANOVA, Tukey’s multiple comparison (B; n=3 independent experiments). *, p<0.05, **, p<0.01. Scale bar: 50 µm in (C).

To determine whether LMNB1 normalization could also rescue astrocyte-mediated myelin impairment through direct cell-cell interactions, we next examined a contact-dependent model by seeding pathological IT3.3 SCR or IT3.3 shRNA-treated astrocytes onto cerebellar organotypic slice cultures (Figure 5A). Representative whole-slice images of the different experimental conditions are shown in Supplementary Figure 5D (panels V, VI). In the presence of IT3.3 shRNA-treated astrocytes, a significant increase of MBP+/CALB+ axon tracts was observed compared with the IT3.3 SCR condition, reaching an approximate 50% increase (Figure 5C). Notably, the extent of MBP⁺/CALB⁺ axon tracts in the IT3.3 shRNA condition was comparable to that observed in the SR condition, suggesting myelin recovery (Figure 5B, C). These findings demonstrate that ADLD astrocytes impair post-lesion myelin restoration through both soluble and contact-dependent mechanisms, and that correcting astrocytic LMNB1 dosage rescues this disease-associated phenotype in an ex vivo demyelination/remyelination model.

**Figure 5.**
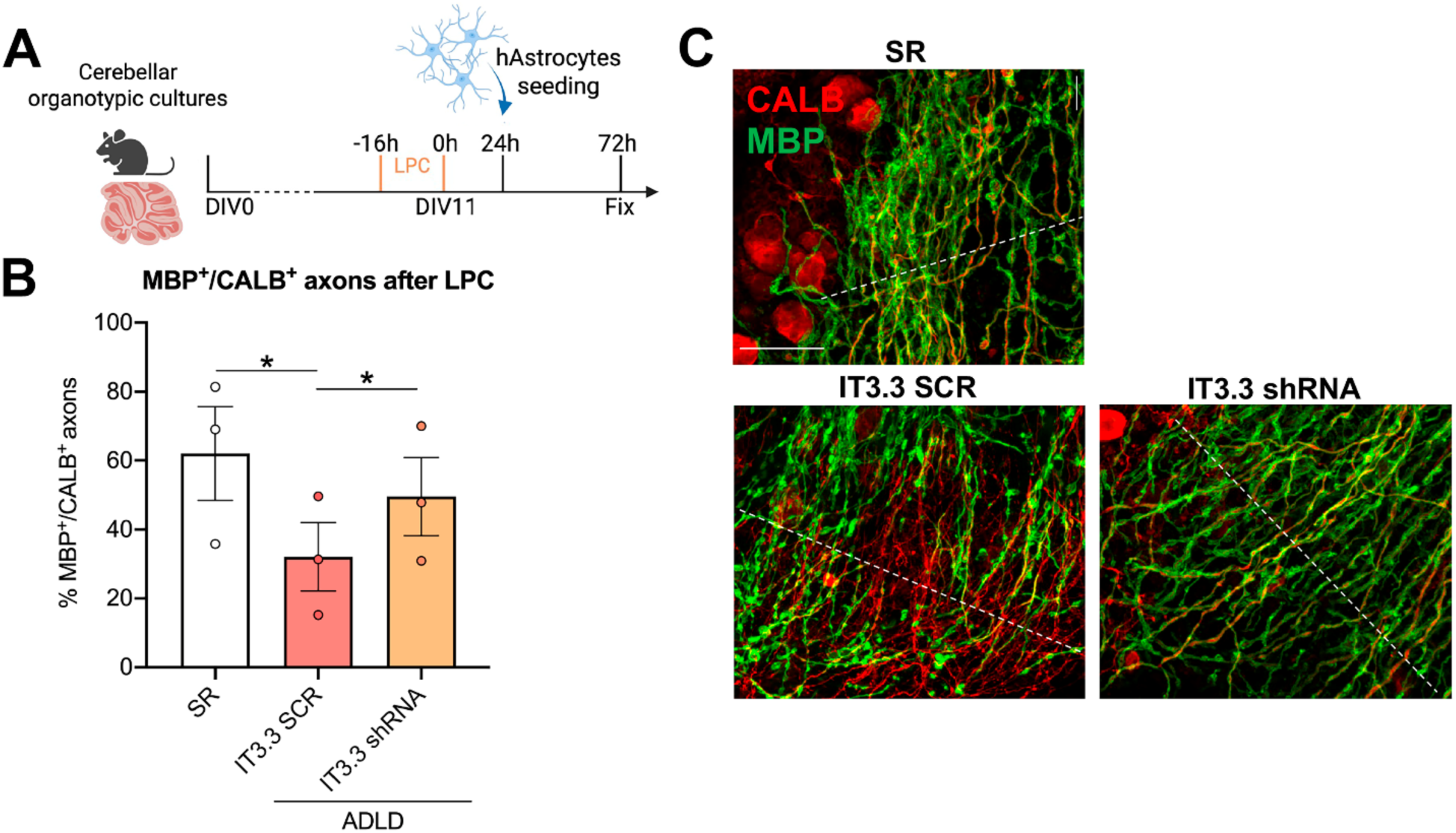
ADLD shRNA astrocytes seeding onto organotypic cerebellar slices improves post-lesion myelin recovery. (A) Schematic representation of the experimental protocol. Timing of LPC application and ADLD astrocytes seeding are shown. Created with Biorender. (B) Fractions of MBP^+^/CALB^+^ axonal profiles, used as proxy of myelin recovery, in SR, IT3.3 SCR and shRNA conditions. Values are expressed relative to the condition of spontaneous recovery (SR) at 72h post LPC. (C) Representative images of cerebellar Purkinje axons in organotypic cultures stained for CALB (red) and MBP (green) in SR, IT3.3 SCR and shRNA conditions. A white line perpendicular to the predominant orientation of CALB^+^ axons is shown as an example of the used approach. LPC, Lysolecithin; AU, Arbitrary Units; CALB, Calbindin; MBP, Myelin Basic Protein. Statistics: RM One-way ANOVA, Tukey’s multiple comparison (B; n=3 independent experiments). *, p<0.05, **, p<0.01. Scale bar: 50 µm in (C).

Altogether, results of ASP-RNAi experiments support LMNB1 overexpression as a key driver of astrocyte-mediated oligodendroglial dysfunction and provide evidence supporting astrocytic LMNB1 normalization as a potential therapeutic strategy to restore glial function and promote myelin repair in ADLD.

#### 2.3 LMNB1 ASP-silencing induces selective metabolic and extracellular-support programs linked to restored astrocyte function

To investigate the transcriptional programs associated with the functional rescue induced by LMNB1 normalization, we further performed single cell (sc)RNA-seq on ADLD astrocytes transduced with SCR or *LMNB1*-targeting shRNA. Again, we focused on IT3.3 as the most informative background in which to probe rescue-associated mechanisms. In contrast to the bulk RNA-seq dataset, which compared ADLD and CTRL astrocytes across independent donor lines, this experiment was designed as a perturbational analysis to identify transcriptional programs responsive to *LMNB1* lowering in diseased cells.

Differential expression analysis identified 220 genes enriched in the *LMNB1*-targeting shRNA condition and 25 genes enriched in scramble-treated cells (Suppl. Table 8A). Consistent with effective target engagement, *LMNB1* itself was retained among the transcripts enriched in the SCR condition, in line with the reduction in *LMNB1* expression achieved by shRNA treatment (Figure 6A). GO analysis of the shRNA-enriched genes (Figure 6B; Suppl. Table 8B) highlighted pathways related to steroid biosynthesis and, more specifically, cholesterol metabolism, including *SQLE*, *NSDHL*, *FDPS*, *DHCR24*. Additional upregulated genes encoded secreted or matricellular factors with potential trophic or protective functions, such as *CRYAB*, *ANXA2*, *CCN1*, *A2M*, and included extracellular matrix-related genes such as *LAMC1*, *COL11A1*, *P4HA2*, *DAG1* (Figure 6C). Beside these genes, several transcriptional alterations previously identified by RNA-seq were not rescued by LMNB1 silencing (e.g. genes related to calcium homeostasis). Together, these data suggest that partial LMNB1 normalization in ADLD astrocytes is associated with selective reprogramming of metabolic and extracellular-support pathways relevant to oligodendroglial function and myelin maintenance, rather than with a global reversion of the ADLD transcriptional state. However, these changes provide potential mechanistic insights into how LMNB1 dosage correction restores astrocyte capacity to support oligodendroglial survival and myelin repair.

**Figure 6.**
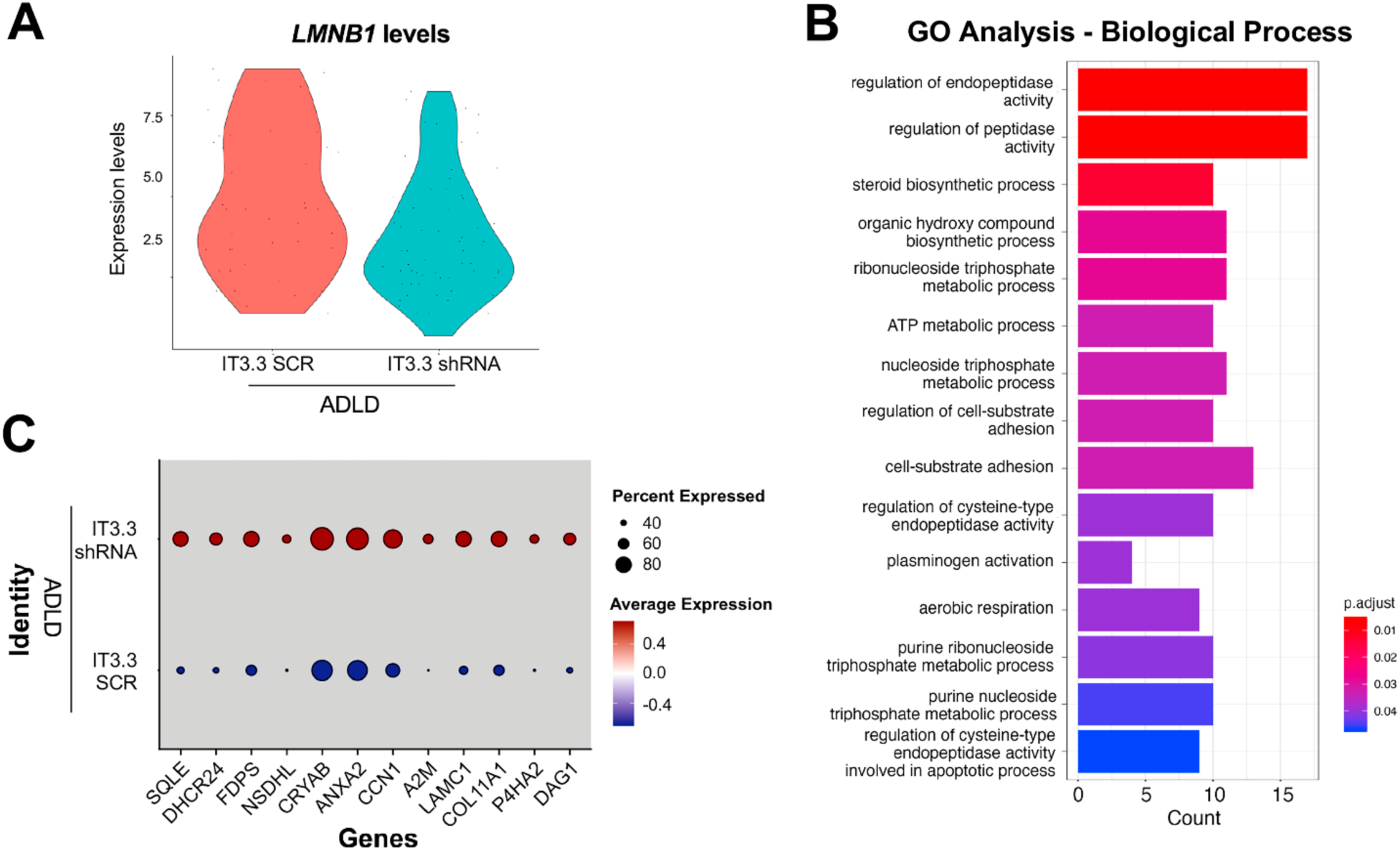
Single-cell RNA-seq identifies LMNB1-responsive metabolic and extracellular-support programs in LMNB1 lowered IT3.3 astrocytes. (A) *LMNB1* is among the genes enriched in IT3.3 SCR astrocytes, consistent with effective *LMNB1* lowering in the shRNA condition. (B) GO analysis of Biological Processes performed on genes enriched in IT3.3 shRNA astrocytes highlights steroid/cholesterol biosynthetic processes and pathways related to extracellular support functions. (C) Representative genes contributing to the enriched categories include cholesterol biosynthesis genes (*SQLE, DHCR24, FDPS, NSDHL*), secreted/trophic factors (*CRYAB, ANXA2, CCN1, A2M*), and extracellular matrix-related genes (*LAMC1, COL11A1, P4HA2, DAG1*).

## Discussion

The present study provides functional validation of an ASP LMNB1-lowering strategy in patient-derived human glial cells and supports ASP-RNAi as a promising therapeutic option for ADLD. Previous work showed that ASP-RNAi can selectively reduce the expression from one of the three LMNB1 alleles in ADLD fibroblasts and directly reprogrammed human neurons (Giorgio et al. 2019). Here, we extend this approach to hiPSC-derived astrocytes 16 and demonstrate that restoration of LMNB1 dosage toward physiological levels produces measurable molecular, cellular and functional benefits. ASP LMNB1 silencing corrected nuclear abnormalities, reduced the detrimental effects of astrocyte-conditioned medium on oligodendroglial viability, and improved post-lesion myelin recovery in organotypic slice cultures. These findings move the therapeutic proof-of-concept beyond molecular target engagement and demonstrate that LMNB1 dosage correction can rescue disease-relevant phenotypes.

The rationale for ASP silencing is closely linked to the genetic basis of ADLD (OMIM #169500), an ultra-rare neurodegenerative disease caused by LMNB1 overexpression in at least the brain of patients. In most cases, increased LMNB1 expression results from intra-TAD duplications encompassing the *LMNB1* gene, whereas less common deletions altering the LMNB1 regulatory landscape have also been reported (Padiath et al. 2006; Giorgio et al. 2013; Giorgio et al. 2015; Nmezi et al. 2025; Dimartino et al. 2024). Because LMNB1 is an essential component of the nuclear lamina, therapeutic interventions should reduce pathological overexpression while preserving physiological gene function. The ASP-RNAi strategy addresses this requirement by preferentially targeting the non-duplicated allele, thereby reducing LMNB1 dosage without complete gene suppression.

Patient-derived astrocytes provided a relevant cellular platform in which to evaluate the efficacy of this strategy within the broader glial environment affected in ADLD. The disease has traditionally been viewed as oligodendrocyte-centered, supported by evidence that oligodendrocyte-targeted Lamin B1 overexpression is sufficient to induce age-dependent demyelination (Rolyan et al., 2015), and by the identification of an oligodendrocyte-specific regulatory element involved in pathological LMNB1 expression (Nmezi et al. 2025). On the other hand, recent neuropathological and experimental studies have also raised the possibility that astrocytes may contribute to disease biology, suggesting ADLD as an astrocytopathology (Ratti et al. 2021a, 2021b; Ratti et al. 2022; Neri et al. 2023; Dimartino et al. 2024). These two scenarios are not mutually exclusive, but both cell types might synergistically act as disease players and, consequently, they could both represent cellular contexts in which LMNB1 dosage correction can provide benefit.

A central advance of this study is the demonstration that LMNB1 normalization in ADLD astrocytes improves higher-order functional readouts involving oligodendroglia and myelin. Conditioned medium from ADLD astrocytes impaired oligodendroglial viability in both murine and human cultures, whereas conditioned medium collected after LMNB1 ASP-silencing showed reduced toxicity. Rescue was more consistent in murine cultures and more variable in human oligodendroglia, but its direction was conserved across systems. This variability may reflect species-specific responses and donor-dependent susceptibility, while the overall convergence supports a relationship between LMNB1 dosage and the functional output of patient-derived astrocytes.

The organotypic slice experiments further strengthen the therapeutic relevance of these findings. In both conditioned-medium and astrocyte-seeding paradigms, LMNB1 lowering increased the recovery of MBP-associated CALB^+^ axonal profiles after LPC-induced demyelination. Although this readout cannot distinguish newly formed myelin from preserved or reorganized myelin structures, it indicates improved post-lesion myelin recovery within a complex multicellular environment. Together, these results are consistent with the broader literature showing that astrocytes actively regulate oligodendrocyte lineage survival, differentiation, and myelin maintenance through metabolic support, secreted factors, extracellular vesicles, and control of the extracellular environment (Camargo et al. 2017; Willis et al. 2020; Molina-Gonzalez et al. 2023; Hu et al. 2023). The attenuation of ADLD astrocytes’ effects following ASP-RNAi LMNB1 lowering links the observed astroglial dysfunction to abnormal gene dosage and establishes astrocytes as a responsive cellular substrate for therapeutic intervention.

The transcriptomic analyses provide initial insight into the cellular programs associated with the functional rescue. Bulk RNA sequencing indicates that ADLD astrocytes display alterations in extracellular-matrix organization, cell adhesion, calcium-related signaling, metabolism, and RNA processing. The single-cell dataset, rather than replicating the bulk CTRL-versus-ADLD comparison, identifies LMNB1-responsive programs within diseased astrocytes, including changes in cholesterol/steroid metabolism, trophic or matricellular genes, and extracellular-support pathways. The limited overlap at the level of individual genes is therefore not unexpected, given the different biological contrast, the single-cell versus bulk resolution, and the partial nature of the ASP-RNAi-driven dosage normalization. Our study does not identify the specific factors responsible for the altered activity of ADLD astrocyte-conditioned medium. The phenotype may result from changes in proteins, lipids, metabolites, extracellular vesicles, or a combination of reduced supportive and increased detrimental signals. The transcriptional data nominate candidate pathways, but direct biochemical and functional studies will be required to define the mediators of rescue. From a therapeutic perspective, however, the key finding is that correction of the upstream LMNB1 dosage defect is sufficient to improve the overall astrocyte output without requiring prior identification of each downstream effector.

Several limitations should be considered. hiPSC-derived astrocytes remain an imperfect proxy for a late-onset and slowly progressive disease as they do not reproduce aging, long-term glial adaptation, or the complete multicellular CNS environment. The bulk transcriptomic comparison involved non-isogenic lines, and the human oligodendroglial assays showed donor-dependent variability. In addition, the *ex vivo* and single-cell analyses focused on IT3.3, which displayed the strongest LMNB1-dependent phenotype. The MBP/CALB-based slice readout is also an indirect measure of myelin recovery and does not demonstrate *de novo* remyelination, and should be interpreted as evidence of improved post-lesion myelin recovery/rescue. Patient coverage represents an additional translational consideration. The ASP molecule used here requires the heterozygosity of a SNP located in the 3’UTR of the *LMNB1* gene (rs #1051644). Its applicability as allele-specific strategy will therefore depend on patient genotype. Clinical development may require a panel of ASP molecules targeting common informative variants or complementary approaches capable of reducing LMNB1 dosage across genetically diverse patients while preserving sufficient physiological expression. Efficient durable delivery throughout the CNS also remains a major challenge for clinical translation. Although the present study establishes biological proof-of-concept, widespread target engagement will be required to address the diffuse white matter pathology of ADLD. Encouragingly, oligonucleotide-based therapies have now demonstrated clinical feasibility for several neurological disorders through intrathecal administration, and a personalized antisense oligonucleotide targeting LMNB1 has recently entered clinical evaluation in ADLD (ClinicalTrials.gov ID: NCT06816498). These advances reinforce the therapeutic rationale of lowering LMNB1 in patients while emphasizing that optimization of CNS delivery, cellular targeting, and long-term efficacy will be critical for the successful translation of both ASO- and RNAi-based approaches.

Despite these limitations, the overall picture is coherent. Our findings support a model in which increased *LMNB1* dosage drives a structurally and transcriptionally altered astrocyte state that is sufficient to impair oligodendroglial support and post-lesion myelin recovery, through both diffusible and likely short-range mechanisms. By demonstrating that normalization of astrocytic LMNB1 dosage through ASP-RNAi rescues these disease-relevant phenotypes, our study expands the pathogenic understanding of ADLD beyond a purely oligodendrocyte-autonomous process. These findings provide a rationale for further development of LMNB1-targeting approaches aimed at restoring glial homeostasis and improving white matter resilience in ADLD.

## Materials and methods

### Experimental Animal

All experimental procedures (i.e. organotypic cultures and OPC cultures) were performed on C57BL/6 mice. The day of birth was considered as postnatal day zero (P0). Groups of 4- 5 mice were housed in transparent polycarbonate cages (Tecnoplast, Buggirate, Italy) provided with sawdust bedding enrichment and striped paper as nesting material. Food and water were provided ad libitum. Mice were maintained on a 12h light/dark cycle, at a room temperature of 21 °C ± 1 °C and a room humidity of 55% ± 5%. All the analyses were in accordance with the guidelines of the National Institutes of Health, the European Communities Council (2010/63/EU) and the Italian Law for Care and Use of Experimental Animals (DL26/2014). It was also authorised by the Italian Ministry of Health and the Bioethical Committee of the University of Turin.

### Human induced pluripotent stem cell cultures

Primary dermal fibroblasts from three canonical ADLD patients carrying a 325Kb intra-TAD duplication encompassing the *LMNB1* gene (IT3.1, IT3.2 and IT3.3; as described in Giorgio et al. 2013, Giorgio et al. 2019 and Dimartino et al. 2024) were reprogrammed using the Sendai virus system (Cytotune, Life Technologies), according to the manufacturer’s instructions. ADLD hiPSC lines and, as control, the commercial hiPSC lines ATCC- DYS0100, GIBCO TMOi001-A and WTSIi-004A (Suppl. Figure 1A) were maintained in vitronectin-coated six-well plates in Essential8 medium (ThermoFisher; A1517001) in a 37°C incubator at 5% CO2. The medium was changed every day. Cells were split when reaching 60-70% confluency, dissociated with Versene solution (ThermoFisher; 15040066) and re-plated in vitronectin coated six-well plates. The study adhered to the Declaration of Helsinki standards and was approved by the Ethics Committee of the IRCCS Mondino Foundation and Alma Mater Studiorum - Università di Bologna [Comitato Etico di Area Vasta

Emilia Centro N° 160-2023-OSS-AUSLBO; EM -Em1-OSS-AUSLBO-23003- Em.1-367- 2023-23003].

### Generation of human Astrocytes

CTRL and ADLD hiPSCs were differentiated into authentic human astrocytes following previously published protocols (Douvaras et al. 2014; Douvaras and Fossati 2015) with minor modifications (Figure 1A). Briefly, cells were dissociated with StemPro Accutase (ThermoFisher, Carlsbad, CA, USA) and seeded as single cells at low density in vitronectin-coated six-well plates in Essential8 medium. At day *in vitro* 0 (DIV0) and for the first seven days, cells were cultured in Neural Induction Medium, and cells were fed daily until DIV7. From DIV8 to DIV12 cells were maintained in culture in N2 Medium with daily medium change. At DIV12, adherent cells were detached via mechanical dissociation using the StemPro EZPassage Disposable Stem Cell Passaging Tool (ThermoFisher, Carlsbad, CA, USA) and cultured in suspension in low-attachment dishes to favour sphere aggregation in N2B27 Medium until DIV20. From DIV20 on, cells were cultured in PDGF medium and medium replacement was performed every other day (see Suppl. Table 5 for detailed media composition). At DIV30, spheres were picked and plated into poly-L-ornithine (0.1 mg/mL, PO; Sigma-Aldrich, Saint Louis, MO, USA) and Laminin (10mg/mL, Lam; Sigma-Aldrich, Saint Louis, MO, USA)-coated six-well plates. At DIV70-80, cells migrated out from the spheres were dissociated with StemPro Accutase for 30 minutes, passed through a 70 μm cell strainer and FACS-sorted for CD49f positivity (Barbar et al. 2020).

Once sorted for CD49f positivity, astrocytes were plated into PO/Lam- coated plates, cultured with different media and at different densities depending on the type of analysis. For ACM collection, astrocytes were plated at high density (50000 cells/cm^2^) in 24-well plates and cultured in minimum medium (DMEM-F12, N2, B27, MEM-NEEA, Insulin). Medium replacement and collection were performed every other day for 4 days, and the conditioned medium was collected, centrifuged and stocked at -80°C. For ICC analyses, astrocytes were plated at low density (3000 cells/cm^2^) in 18-well μ-Slides (Ibidi GmbH, Gräfelfing, Germany), cultured in PDGF medium for 3 days and fixed with 4% paraformaldehyde (PFA) in 0.1 M sodium phosphate buffer (PB). For RNA-seq analyses, immediately after sorting astrocytes were centrifuged, frozen and kept at -80° until RNA extraction.

### Immunofluorescence, confocal acquisitions and morphological analyses on CD49f- sorted hAstrocytes

For immunofluorescence staining, astrocytes were incubated for 24 h at 4 °C in a solution of 0.01 M PBS, pH 7.4, containing 0.5% Triton X-100, 2% normal donkey serum and primary antibodies (listed in Suppl. Table 6). Cells were then incubated for 2 h at room temperature, in a solution of 0.01 M PBS, pH 7.4, containing 1% normal donkey serum, 4′,6-diamidino-2- phenylindole dihydrochloride (DAPI; Fluka, Milan, Italy) and appropriate secondary antibodies (Suppl. Table 6). All the images were acquired with the inverted confocal microscope ZEN LSM800 (Zeiss, Oberkochen, Germany), using 20× magnification in a field of 319.45 × 319.45 µm, zstep = 1 µm. For each technical replicate (n = 3) of each cell line (n = 6), at least 15 astrocytes were analyzed, for a total of 193 CTRL astrocytes (C1: n=55; C2: n=72; C3: n=66), and 195 ADLD astrocytes (IT3.1: n=48; IT3.2: n=66; IT3.3: n=78). Of note, the quantification of GFAP, AQP4, and LMNB1 protein levels throughout immunofluorescence staining required a standardized procedure, in order to avoid technical bias. The same antibody aliquots were used for all the quantification, in order to avoid different specificities, and staining was carried out in parallel on the different samples. Images were acquired using the same confocal parameters, and pixel saturation was avoided, in order to appreciate the entire expression spectrum. Moreover, the images were not post-processed. The collected images were analysed with Fiji ImageJ software (Schindelin et al. 2012) to measure Integrated Density of LMNB1 staining (Figure 1E; Suppl. Figure 4D), assess LMNB1 protein pattern (Figure 1C, F-G, see Giorgio et al. 2019 for detailed categorization parameters), quantify cell, soma, and nuclear area, as well as GFAP and AQP4 levels (Suppl. Figure 2C-G).

### RNA isolation and quantitative Real Time PCR (qPCR)

Total RNA was extracted from hiPSCs and CD49f^+^ astrocytes using PureLink RNA Mini Kit (ThermoFisher, Carlsbad, CA, USA) and cDNA was generated using the High Capacity cDNA Reverse Transcription Kit (ThermoFisher, Carlsbad, CA, USA). The expression levels of *LMNB1*, *SOX2* (SRY-Box Transcription Factor 2), *POU5F1* (POU Class 5 Homeobox 1, OCT4), *PAX6* (Paired Box 6), *OLIG2* (Oligodendrocyte Transcription Factor 2), *NKX2-2* (NK2 Homeobox 2), *HMBS* (hydroxymethylbilane synthase), *18S* (18S rRNA) and *GAPDH* (Glyceraldehyde-3-Phosphate Dehydrogenase) were measured with predesigned TaqMan™ assays (Applied Biosystems, Carlsbad, CA, USA) and listed in Suppl. Table 7. Expression levels of *PTPRT* (Protein Tyrosine Phosphatase Receptor Type T), *GRM5* (Glutamate Metabotropic Receptor 5), *SHMT2* (Serine Hydroxymethyltransferase 2) and *ACTB* (Actin Beta) were analysed by combining the RealTime Ready Universal Probe Library (UPL, Roche Diagnostics, Monza, Italy) with the primers indicated in Suppl. Table 7. Reactions were carried out in triplicate and Real Time data were collected on the Applied Biosystems StepOnePlus Real-Time PCR System with StepOne™ Software. Data analysis was performed with Microsoft Excel (Microsoft Office 365). A relative quantification approach was used, according to the 2^-ΔΔCT^ method. *HMBS, GAPDH, 18S* or *ACTB* were used to normalize expression levels.

### Bulk RNA sequencing and bioinformatic analyses

Generation of RNA-Seq data was performed by GENARTIS srl (Verona, Italy), as follows. Total RNA was extracted as described above and RNA purity was measured at a NanoDrop Spectrophotometer (ThermoFisher, Carlsbad, CA, USA) while RNA integrity was assessed using the RNA 6000 Nano Kit on a Bioanalyzer (Agilent Technologies). All samples showed an RNA integrity number (RIN) >7. RNA samples (two experimental replicates/cell line, 12 samples in total) were quantified using the Qubit RNA BR Assay Kit (ThermoFisher, Carlsbad, CA, USA). RNA-Seq libraries were generated using the NEBNext Ultra II Directional RNA Library Prep Kit (Illumina) from 10ng of material after poly(A) capture and according to manufacturer’s instructions. Quality and size of RNA-Seq libraries were assessed by capillary electrophoretic analysis with the Agilent 4200 Tape station (Agilent Technologies). Libraries were quantified by real-time PCR against a standard curve with the KAPA Library Quantification Kit (KapaBiosystems, Wilmington, MA, USA). Libraries were pooled at equimolar concentration and sequenced on a NovaSeq6000 (Illumina) generating >20 million fragments in 150PE mode for each sample. Sequencing read trimming and quality of reads were assessed using FastQC software (http://www.bioinformatics.babraham.ac.uk/projects/fastqc/). Batch Correction was applied to avoid biases due to different differentiation experiments. Filtered reads (∼80% of trimmed reads are mapped against the transcriptome) were aligned to the Human reference genome GRCh38 (Ensembl) using STAR (v2.7.6a) with default parameters and quantMode TranscriptomeSAM option that output alignments translated into transcript coordinates. After reads mapping, the distribution of reads across known gene features, such as exons (CDS, 5’ UTR, 3’ UTR), introns and intergenic regions, was verified using the script read_distribution.py provided by RSeQC package (v3.0.1). Read counts on genes were quantified using RSEM (v.1.3.3). Gene-level abundance, estimated counts and gene length obtained with RSEM were summarized into a matrix using the R package tximport (v1.18.0) and the differential expression analysis was performed with DESeq2 (v1.30.0) integrating the ‘‘differentiation experiment’’ as variable in the model. To generate more accurate Log2 FoldChange estimates, the shrinkage of the Log2 FoldChange was performed applying the apeglm method. Gene Ontology (GO) enrichment analysis was performed using clusterProfiler, an R Package for comparing biological themes among gene clusters (Yu et al. 2012; Bioconductor version: Release 3.18.0). Differentially expressed genes (DEGs) with p adjusted value ≤ 0.05 and |log2FC| ≥ 0.5 were included in the analysis. In addition to strictly significant genes, a limited number of transcripts with adjusted p-values close to the significance threshold were retained based on consistent expression trends across cell lines/experiments and established relevance to astrocyte biology. FDR adjusted p value (q-value) <0.05 was used as a threshold for GO terms.

For Differentially Expressed Isoforms (DET) analysis, reads were aligned to the Human reference genome GRCh38 (Ensembl) using STAR two-pass (v2.7.6a) method and read counts on isoforms were quantified using Salmon. Isoforms-level abundance, estimated counts and gene length obtained with Salmon were summarized into a matrix using the R package tximport (v1.18.0) and the differential expression analysis was performed with EBseq. To generate more accurate Log2 FoldChange estimates, the shrinkage of the Log2 FoldChange was performed using the apeglm method.

For Differentially Alternative Spliced Genes (DASGs), the RNA sequencing output data were subjected to a computational analysis using the rMATS bioinformatics algorithm and Junction Read Counts (JC) were used to obtain informative reads for alternative splicing events. DASGs were categorized under five different spliced groups, namely (1) skipped exon (SE), (2) retained intron (RI), 3) alternative 3′ splice site (A3SS), 4) alternative 5′ splice site (A5SS) and 5) mutually exclusive exons (MXEs) (Suppl. Figure 3D). The percent spliced in (or inclusion [PSI], or Ψ) was determined for all genes/splicing events between the ADLD versus CTRL, on which basis delta Ψ (ΔΨ = ΨADLD – ΨCTRL) was determined for each gene/event. We selected Differentially Alternative Spliced Genes (DASGs) based on a False Discovery Rate (FDR) ≤ 0.05 and a |ΔΨ| > 0.1.

Gene Set Enrichment Analysis (GSEA) (version 4.1.0, the Broad Institute of MIT and Harvard; https://www.gsea-msigdb.org/ gsea/downloads.jsp) was performed and to select biologically-significant pathways we focused on gene-sets with maximal Enrichment Score (|ES|) > 0.3 and a q-value<0.25, according to GSEA guidelines.

### Magnetic Activating Cell Sorting (MACS) and primary murine OL cultures

MACS sorting was performed as described in Lorenzati et al. 2021. Briefly, after tissue dissociation with a papain + DNAseI solution (papain 1.5 mg/ml, l-cysteine 360 μg/ml, DNAseI 1000U/ml in MEM; all from Sigma-Aldrich, Saint Louis, MO, USA), mouse OPCs were enriched by positive selection using an anti-PDGFRα antibody conjugated to magnetic beads, according to the instructions of the manufacturer (Miltenyi Biotech GmbH, Bergisch Gladbach, DE). MACSorted OPCs were plated onto poly-d-lysine (1 μg/ml, Sigma-Aldrich, Saint Louis, MO, USA) coated 96-well plates (50000 cells/cm^2^) in a proliferative medium including Neurobasal, 1X B27 (Invitrogen, Milan, Italy), 2 mM l-glutamine (Sigma-Aldrich, Saint Louis, MO, USA), 10 ng/ml PDGF-BB and 10 ng/ml human bFGF (Miltenyi Biotech GmbH, Bergisch Gladbach, DE). Purity of the MACS-selected OPCs was verified by immunocytochemistry (more than 95% of the cells were NG2-positive (+) at 6 h post-plating). Cells were cultured 1DIV in proliferative medium and 2DIV in differentiative conditions, avoiding proliferative mitogens (i.e. PDGF-BB and bFGF). At DIV2 half media change was performed using Astrocyte-Conditioned Medium (ACM) from high-density cultures of CTRL, ADLD or shRNA ADLD astrocytes. Of note, for these experiments only CTRL2 and CTRL3 cell lines were used. Cells were monitored on the IncuCyte® Live Cell Analysis System (Sartorius, Gottingen, Germany) and scanned at 0h (after adding ACM), 6h and 16h. Cytotox dye was added to culture media to detect OL cell death at DIV2, together with ACM.

### Cultures of human induced OLs

Induced hOL cultures were generated as in (García-León et al. 2020). Cryopreserved stocks were gently thawed, centrifuged and plated at high density (50000 cells/cm^2^) into PO/Lam-coated 96-well plate and cultured with PDGF Medium with the addition of 1μg/mL of doxycycline. At DIV2 post plating half media change was performed using ACM from high-density cultures of CTRL, ADLD or shRNA ADLD astrocytes. Of note, for these experiments only two control cell lines were used. Cells were monitored on the IncuCyte® Live Cell Analysis System (Sartorius, USA) and scanned at 0h (after adding ACM), 4h, 8h and 12h. Cytotox dye was added to culture media to detect OL cell death at DIV2, together with ACM.

### Live Cell Imaging analysis on mOLs and hOLs

The assessment of m/hOL viability has been conducted thanks to the use of the Incucyte® Cytotox NIR Dye (Sartorius, Gottingen, Germany), formulated to detect cell membrane integrity disruption to quantify cell death. Cytotox NIR Object counts (8h and 16h vs 0h for mOLs; 4h, 8h and 12h vs 0h for hOLs) analysis of Incucyte® Imaging Analysis Software was used to detect cell death events over time (9 fields/well, 2-4 experiments/condition, CTRL or ADLD, SCR or shRNA, where applicable).

### Lentiviral transduction of Mixed Glial Cultures

Lentiviral transduction of ADLD Mixed Glial Cultures (DIV50 of differentiation) was used to target the ‘T’ allele (non-duplicated), in order to silence one of the three copies of *LMNB1*. Based on the results in (Giorgio et al. 2019), one ASP-siRNA (SNP position 4; the most efficient in terms of LMNB1 downregulation; see Giorgio et al. 2019) targeting the T allele was converted to generate a EGFP-tagged short- hairpin RNA expression vector and cloned into Recombinant Lentivirus particles (LV-ASP-T4 shRNA; pLV[shRNA]-EGFP:T2A:Puro-U6; viral titer 1.62x10^9 TU/mL; outsourced to Vector Builder). As negative control, we used a commercial EGFP-tagged scramble shRNA control lentivirus (scramble shRNA; viral titre 10.4 x 10^8 TU/ml, VB151023-10034; Vector Builder). The lentiviral particles produced were resuspended in Hank’s balanced salt solution buffer. Viral stocks were stored at -80°C until use. Cells were transduced with lentiviral particles (shRNA or SCRAMBLE) at a MOI of 10 in the presence of Polybrene 1:1000, for two times (the second infection was performed three days after the first one; Suppl. Figure 4A). Transduction efficiency was controlled two, seven and twenty days after transduction by counting the GFP^+^ over the total number of cells, reaching 80% of transduced cells (Suppl. Figure 4B). Before seeding, mixed glial cultures were FACS sorted for CD49f and GFP positivity, to collect only transduced astrocytes.

### Organotypic culture of cerebellar slices

Organotypic cerebellar slices were prepared from C57BL/6 mouse pups aged 7 postnatal days as already described (Hussain et al. 2011; Meffre et al. 2015). After decapitation, brains and cerebellum were dissected out in a solution of PBS and glucose 0.1% After removal of the meninges, 350 μm-thick cerebellar parasagittal slices were cut using a tissue chopper and transferred onto membranes of 30-mm Millipore culture inserts with 0.4-μm pore size (Millicell, Millipore, Bedford, MA, USA). Slices were maintained in culture in six- or twelve-well plates containing 1 mL or 600 μl of medium, respectively, at 35°C and 5% CO2. Serum-free culture medium was composed of DMEM-F12, 1% N2, 2% B27, 1% Glutamax 100X, 0.2% Fungizone, 1% Penicillin/Streptomycin. Cultures were maintained for 10 days in vitro (10DIV), and medium was changed every 3 days.

### Lysolecithin-induced demyelination and remyelination in organotypic cerebellar slice cultures

To induce demyelination, medium was removed after 10 DIV and fresh medium with lysolecithin (LPC; 0.5 mg/mL; Sigma, Saint Louis, MO) was added and incubated for 16h at 37°C following Birgbauer et al. (2004). After incubation, the LPC-containing medium was replaced with fresh medium. To identify the optimal timepoint for ACM application or astrocytes seeding, a time course analysis of LPC-induced myelin loss and spontaneous recovery was performed (Suppl. Figure 5A). The fraction of MBP^+^/CALB^+^ structures over CALB^+^ axons (see below) was used to monitor these changes, as decreased after LPC exposure, and subsequently increased, during bona fide recovery. Although this measure does not specifically identify newly formed myelin segments, its temporal dynamics supported its use as a proxy for myelin loss and recovery in this experimental setting. The loss of MBP^+^/CALB^+^ structures peaked at 16 h after LPC treatment, while recovery became apparent from 24 h. Accordingly, astrocytes or ACM were applied to organotypic slices 24 h after LPC treatment to evaluate their effects on post-lesion myelin recovery.

### ACM application and seeding of astrocytes on organotypic slices

24h post LPC washout, ACM was mixed with Matrigel to a final Matrigel concentration of 10% (v/v) and, subsequently, 10 μl of the ACM–Matrigel mixture was carefully applied onto each organotypic slice. Cerebellar slices were maintained in culture without changing the media for 48h. After 48h (72h after LPC treatment), slices were fixed with 4% PFA and washed with PBS for subsequent immunostaining and analysis. Representative Tile Scan images are shown in Suppl. Figure 5D.

For seedings, IT3.3 shRNA or IT3.3 SCR astrocytes were centrifuged at 3500 rpm for 5 minutes and resuspended in PDGF Medium with 10% Matrigel at a concentration of 25,000 cells/μL. Cells were seeded onto the slices (2 μl/slice) at 12DIV using a P10 pipette. Cell seeding was confirmed by epifluorescent microscopy. Slices were cultured for 48h before fixing for further analysis.

### Immunofluorescence, confocal acquisitions and imaging analyses on organotypic cerebellar slices

To perform the immunohistological staining, cerebellar slices were washed three times for 10 minutes in PBS, then permeabilized in PBS + Triton 0.5% for 2h. Then, the slices were incubated for 48h at 4°C in a solution of 0.01 M PBS, pH 7.4, containing 0.5% Triton X-100, 2% normal donkey serum and primary antibodies (listed in Suppl. Table 6). After 2 days, slices were washed three times for 10 minutes in PBS and incubated for 2 h at room temperature, in a solution of 0.01 M PBS, pH 7.4, containing 1% normal donkey serum, DAPI and appropriate secondary antibodies (Suppl. Table 5). Finally, cerebellar slices were mounted on glass slides and covered with a few drops of Mowiol before applying the coverslips. Of note, staining was carried out in parallel on the different samples.

All the images were acquired with ZEISS Axio Imager Z2 equipped with Apotome 3 (Carl Zeiss Microscopy GmbH, Jena, Germany) and 40× Plan-Neofluar objective (NA 0.75) within a region of interest (ROI) of 212.52 × 177.33 µm, zstep = 1 µm, zstack = 10. Sampling was rigorously performed in order to avoid technical biases: 3-5 region of interest (ROI) corresponding to the apices of different lobules were acquired for each slice (SR: 1 slice/timepoint, three experimental replicates; ACM: 1 slice/condition, three experimental replicates; astrocytes seeding: 1 slice/condition, three experimental replicates). In astrocytes seeding experiments, ROIs were selected according to the location of seeded astrocytes. Preference was given to granular layers in the apical region of the lobules when seeded astrocytes were present there. Otherwise, apices of lobules containing seeded astrocytes in the corresponding white matter were analyzed. Images were acquired using identical parameters across conditions, avoiding pixel saturation, and were not post-processed. Quantifications of MBP^+^/CALB^+^ fibers were performed with Fiji/ImageJ software (Schindelin et al. 2012) following Llufriu-Dabén et al. (2019), with some modifications. For each ROI, a straight line was traced perpendicular to the dominant direction of CALB^+^ axons and the number of CALB^+^ axons crossing the line was counted. Then, the same counting was performed for CALB^+^ axons adjacent to MBP^+^ structures. In Supplementary Figure 5B a Z projection of the SR condition (in Figure 4) is shown, with high magnification examples of CALB^+^ axons scored as MBP associated (myelinated axons, yellow asterisks) and CALB^+^ axons not MBP associated (red asterisks). The validation of this criterion is shown in Supplementary Figure 5C. The percentage of MBP^+^/CALB^+^ axons (myelinated axons) was calculated as the ratio of MBP^+^/CALB^+^ structures over total axons in each ROI. Results shown in Figure 4B and 5B are expressed as the percentages of MBP^+^/CALB^+^ axons in the investigated conditions.

### Single cell sequencing library preparation and bioinformatic analysis

ScRNA-seq was initially undertaken to resolve transcriptional consequences of LMNB1- targeting shRNA (Giorgio et al 2019) at single-cell resolution. Although this analysis did not reveal robust shRNA-specific transcriptional states, it remained informative by identifying convergent gene programs associated with *LMNB1* lowering across treated astrocytes. We therefore used the dataset to define shared LMNB1 dosage-responsive pathways rather than construct-specific effects.

For single cell library preparation, 100.000 cells of each genotype were collected and dissociated using trypsin/EDTA for 5 min at room temperature. Cells were then washed with PBS, and the resulting cell suspension was used to sort individual live cell in 96 well plate. Full length single cell RNA-seq was performed using a modified version of the Smart-seq2 protocol (Picelli et al. 2013) as in Proserpio et al. (2022). Briefly, individual cells were sorted into 96 well plates containing lysis buffer in presence of RNase inhibitor, dNTPs and oligodT. Reverse transcription of the polyadenylated RNA was performed with SuperScriptII and Template Switching Oligos. The resulting cDNA was amplified with 25 cycles of PCR and libraries were prepared for sequencing with miniaturized NexteraXT Illumina protocol. Libraries were sequenced on Illumina NextSeq 1000 System (single-end 100 bp reads), reaching a median of ∼ 300,000 generated reads per cell.

### Single cell RNA-seq data analysis

Following quality controls (performed with FastQC v0.11.2 (https://www.bioinformatics.babraham.ac.uk/projects/fastqc), sequencing reads were processed with TrimGalore! v0.5.0 (https://www.bioinformatics.babraham.ac.uk/projects/trim_galore) to perform quality and adapter trimming (parameters:–stringency 3 –q 20). Trimmed reads were next aligned to the human reference genome (GRCh38 Ensembl release 98) using STAR v2.7.1a with options: *–outFilterMultimapNmax 10, –outFilterMultimapScoreRange 1, – outFilterMismatchNmax 999, –outFilterMismatchNoverLmax 0.04*. Gene expression levels were quantified with featureCounts v1.6.1 (https://subread.sourceforge.net/, options: *-t exon -g gene_name*) using the GENCODE Release 32 annotation. Multi-mapped reads were excluded from quantification.

The following criteria were applied to exclude low-quality cells from subsequent analyses: <50,000 assigned reads; <2000 detected genes; more than 50% of reads assigned to mitochondrial genes. Gene expression counts were next analysed using the Seurat v4.0.1 package (Hao et al. 2021). Read counts were first coverage-normalized and log-transformed (*NormalizeData* function with default parameters); next, variance modelling for feature selection was carried out using the *FindVariableFeatures* function (parameters: *selection.method= ”vst”, nfeatures = 2000*), selecting the top 2000 variable genes for the subsequent analyses. The selected features were scaled and centred using the *ScaleData* function, which also allowed to regress-out unwanted sources of variability (i.e. library preparation batch, percentage of mitochondrial reads, by setting the parameter *vars.to.regress*).

### Statistical analyses

Graphics and statistical analyses were performed with Graphpad Prism version 7 (Graphpad software, San Diego, CA). The Shapiro–Wilk test was first applied to test for a normal distribution of the data. When normally distributed, unpaired Student’s t test (to compare two groups), One-way ANOVA, RM One-way ANOVA and Two-way ANOVA test (for multiple group comparisons) followed by Tukey’s/Sidak’s post hoc analysis were used. Statistics also included Chi-square test or Fisher’s exact test (to compare frequencies). In all instances, P < 0.05 was considered as statistically significant. Histograms represent mean ± standard error (SE). Statistical differences were indicated with *P<0.05, **P<0.01, ***P< 0.001, ****P < 0.0001. The list of the applied tests, F values and values for n, results of post hoc analyses are included in Suppl. Table 9. Supporting data are available within the article or can be provided upon request.

## Acknowledgments and Fundings

This work was supported by ELA International (Grant No. 2019-00612) and by local funds from the University of Turin. EG, PC, and SR were supported by the European Union – NextGenerationEU and by the Italian Ministry of University and Research (MUR) under the National Recovery and Resilience Plan (NRRP), project MNESYS (PE0000006), “A Multiscale Integrated Approach to the Study of the Nervous System in Health and Disease” (DN. 1553, 11 October 2022), and by the Italian Ministry of Health through the “Ricerca Finalizzata” program (Grant No. GR-2021-12373348). FJP was supported by a Marie Skłodowska-Curie Actions (MSCA) grant (Grant No. 101106517). We gratefully acknowledge the ADLD family members who participated in this study and Prof. Catherine Verfaillie for providing human induced oligodendrocytes (hOLs). Patent approved for the ASP-siRNA sequences n. 102017000121288 entitled ‘RNA interference mediated therapy for neurodegenerative diseases’ to EG and ABru.

## Ethics approval statement

The use of hiPSC lines was approved by the Bioethical Committee of the University of Turin. Animal experimentation was approved by the Italian Ministry of health.

## Author contributions

ML, AB and EG designed the study and supervised the project. ML, FJP, EG, AB wrote the manuscript; ML, AC, MR, MKA, FJP, ES, EN, GP, VP performed experiments; ML, AC, MR, FL, GT, VC analyzed data; JAGL provided induced human oligodendrocytes; AB, PC, SR and LC provided human cell samples; all authors contributed to the revision of the manuscript and approved the final version.

