## Supplementary Figures for "Functional validation of allele-specific LMNB1 silencing in patient-derived astrocytes as a therapeutic option for Autosomal Dominant Leukodystrophy"

**A**

| Cell Line | Age | Sex | Reprogramming method | Source |
|---|---|---|---|---|
| **ATCC-DYS0100** | Unknown | Unknown | Sendai (POU5F1, SOX2, KLF4, MYC) | Human foreskin fibroblast |
| **GIBCO TMOI001-A** | Unknown | Female | Episomal 8POU5F1, SOX2, KLF4, MYC, NANOG, LIN28, SV40 T) | CD34+ progenitors |
| **WTSII004-A** | 35-39 years old | Male | Sendai (POU5F1, SOX2, KLF4, MYC) | Human fibroblasts |
| **IT3.1** | 53 years old | Male | Sendai (POU5F1, SOX2, KLF4, MYC, NANOG, LIN28, SV40 T) | Human fibroblasts |
| **IT3.2** | 56 years old | Female | Sendai (POU5F1, SOX2, KLF4, MYC, NANOG, LIN28, SV40 T) | Human fibroblasts |
| **IT3.3** | 45 years old | Male | Sendai (POU5F1, SOX2, KLF4, MYC, NANOG, LIN28, SV40 T) | Human fibroblasts |

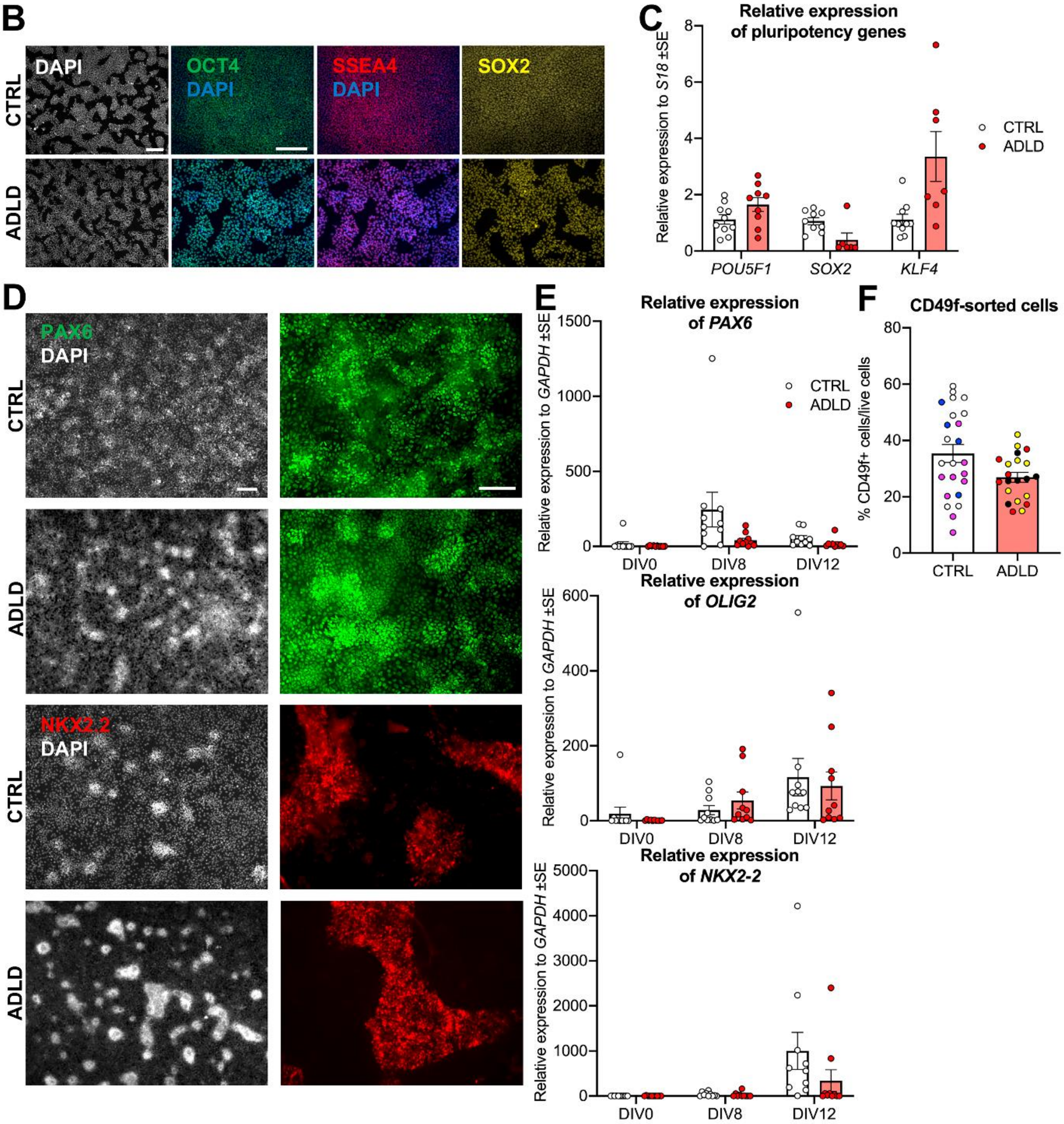


**Suppl. Figure 1. CTRL and ADLD hiPSCs similarly respond to gliogenic induction and generate astrocytes.** (A) Summary of the hiPSC lines used in this study, including (when available) donor age, sex, reprogramming method, and cell source. (B) Representative immunostaining showing that both CTRL and ADLD hiPSCs express pluripotency markers (OCT4 in green, SSEA4 in red, SOX2 in yellow).

(C) qRT-PCR quantifications of pluripotency markers (*POU5F1* codes for OCT4). (D) Differentiation of CTRL and ADLD hiPSCs into PAX6$^{+}$ neural progenitors (green, top panels) and NKX2.2$^{+}$ glial-committed cells (red, bottom panels). DAPI (white) counterstains nuclei and highlights the formation of densely packed 3D cellular aggregates, a key intermediate step in glial commitment. (E) qRT–PCR analysis of *PAX6*, *OLIG2*, and *NKX2-2* mRNA at key time points (DIV0, DIV8, DIV12) shows no significant differences between CTRL and ADLD cultures, indicating comparable differentiation kinetics. (F) Flow-cytometry sorting reveals similar yields of CD49f$^{+}$ astrocytes from CTRL and ADLD lines. Individual dots represent independent hiPSC lines (color-coded; ATCC DYS0100 - blue, GIBCO TMOi001-A - white, WTSIi004-A - pink, IT3.1 - red, IT3.2 - yellow, IT3.3 - black). OCT4, POU Class 5 Homeobox 1; SSEA4, Stage-Specific Embryonic Antigen 4; SOX2, SRY-box Transcription Factor 2; PAX6, Paired Box 6; NKX2-2, NK2 Homeobox 2; OLIG2, Oligodendrocyte Lineage Transcription Factor 2; CD49f, Integrin alpha-6; KLF4 (Kruppel-like factor 4). Statistics: Two-way ANOVA, Sidak's multiple comparison (E; n=3 to 4 independent experiments/cell line; 3 ADLD vs 3 CTRL cell lines); Unpaired two-tailed T-test (F; 3 ADLD vs 3 CTRL cell lines). Scale bar: 100 μm in (B, D).

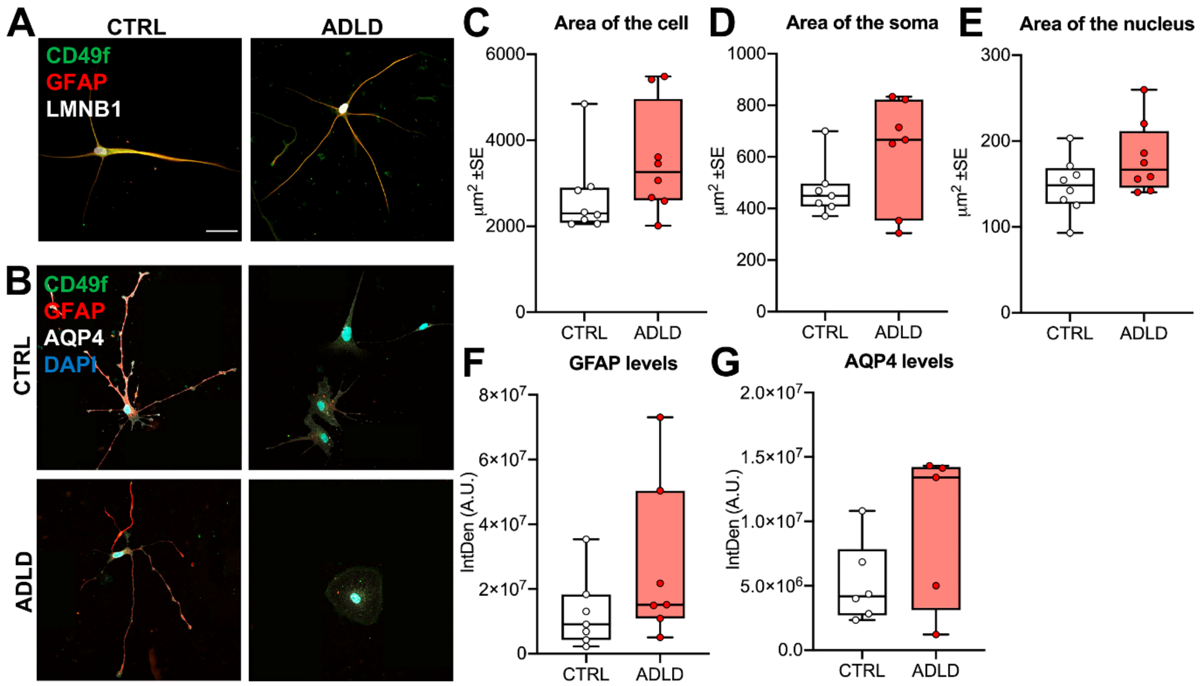


**Suppl. Figure 2. ADLD astrocytes display limited morphological and neurochemical alterations.** (A) Representative images of CTRL and ADLD astrocytes stained for CD49f (green), GFAP (red) and LMNB1 (white). (B) Representative images of CTRL and ADLD astrocytes decorated for CD49f (green), GFAP (red) and AQP4 (white). Nuclei are counterstained with DAPI. (C–G) Quantifications of morphological and marker-based parameters in CTRL and ADLD astrocytes. Measured features include total cell area (C), soma area (D), nuclear area (E), GFAP immunoreactivity (F), and AQP4 immunoreactivity (G). CD49f, Integrin alpha-6; GFAP, Glial Fibrillary Acid Protein; AQP4, Aquaporin 4; LMNB1, Lamin B1; AU, Arbitrary Units. Statistics: Unpaired two-tailed T-test (C-G, n= 2 to 3 independent experiments/cell line; 3 ADLD vs 3 CTRL cell lines); ns, not significant. Scale bar: 50 µm in (A, B).

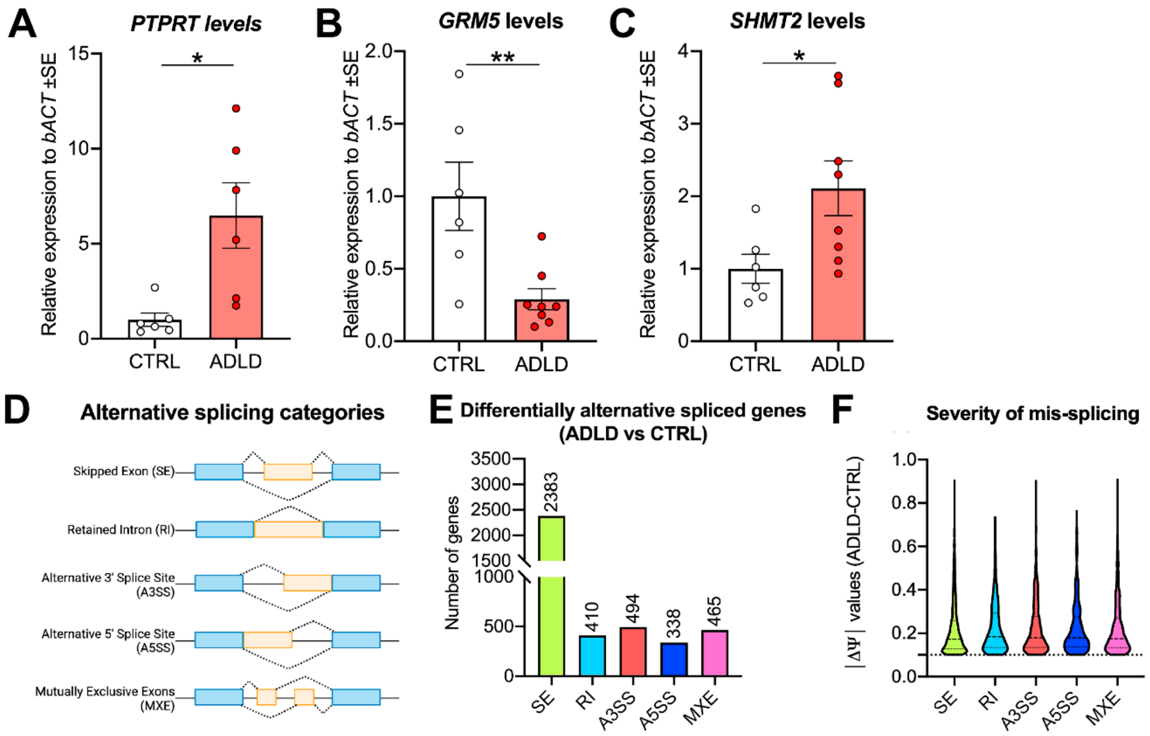


**Suppl. Figure 3. Analysis of transcriptional and alternative splicing alterations in ADLD astrocytes.** (A-C) qRT-PCR quantifications of relevant genes found dysregulated in bulk RNA-seq analysis: *PTPRT* (ECM organization), *GRM5* (Calcium Ion Homeostasis), *SHMT2* (cellular metabolism). (D) Investigated categories of Alternative Splicing. (E) Number of differentially spliced genes, detected under each splicing category, in ADLD astrocytes vs CTRL astrocytes, based on False Discovery Rate (FDR) $\leq 0.05$ and a $|\Delta\Psi| > 0.1$. (F) Severity of mis-splicing, i.e. the distribution of the differentially spliced events based on $\Delta\Psi$ values (see Methods section). *PTPRT*, Protein Tyrosine Phosphatase Receptor Type T; *GRM5,* Glutamate Metabotropic Receptor 5; *SHMT2*, Serine Hydroxymethyltransferase 2. Statistics: Unpaired T-test (A-C, n=2 to 3 independent experiments/cell line); *, $p<0.05$; **, $p<0.01$.

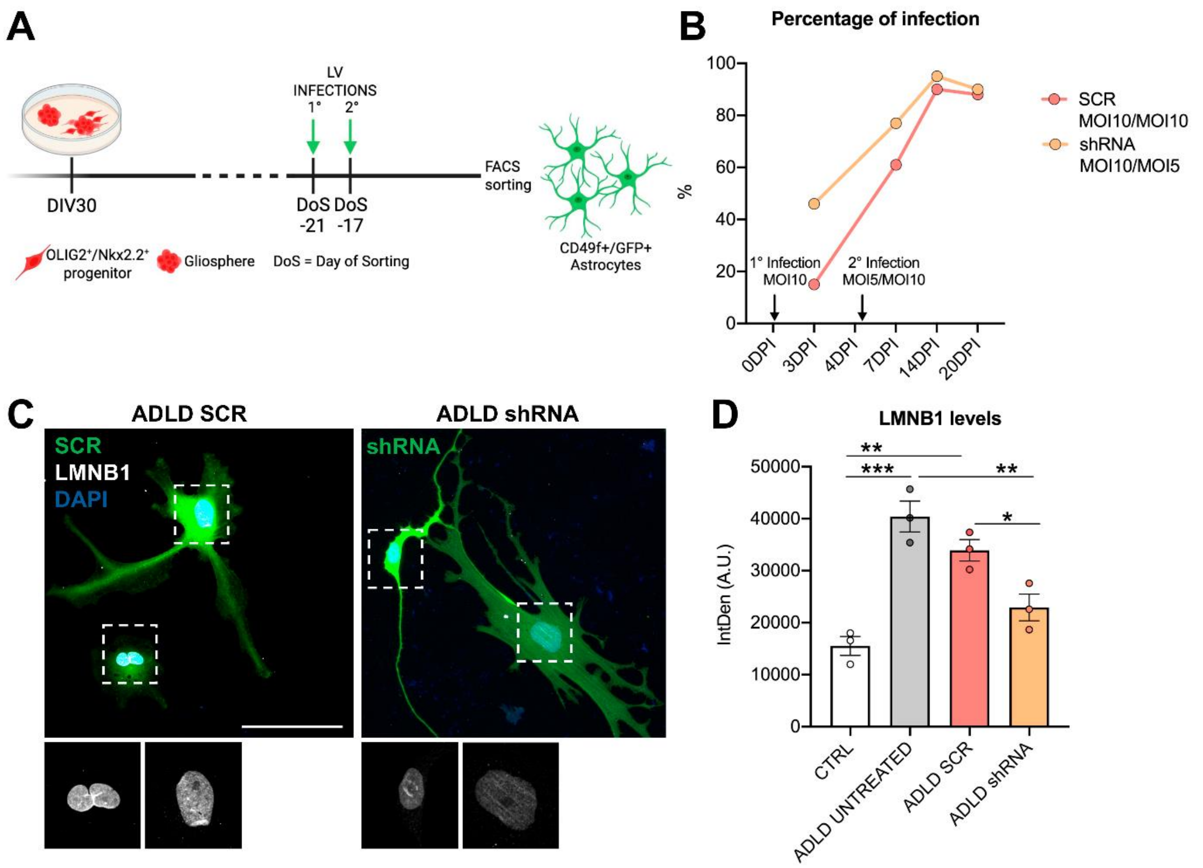


**Supplementary Figure 4. Experimental workflow and validation of LMNB1-targeting shRNA in ADLD astrocytes.** (A, B) Schematic representation of the differentiation protocol with the timing of LV infections (A), and representative percentage of infected cells over total cells (B), 21 (0 DPI) and 17 (5 DPI) days before sorting for GFP$^+$/CD49f$^+$ astrocytes. (C) Representative images of GFP$^+$ SCR (left) and shRNA (right) ADLD astrocytes three days after sorting, stained for LMNB1 (white). Insets show the presence of nuclear abnormalities in the SCR condition that virtually disappear in the shRNA condition. DAPI counterstains cell nuclei. (D) Quantification of LMNB1 protein abundance, showing a decrease in LMNB1 integrated density in shRNA condition. MOI, Multiplicity Of Infection; AU, Arbitrary Units; IntDen, Integrated Density. Statistics: One-way ANOVA, Tukey's multiple comparison (D; n=3 cell lines/condition). *, $p<0.05$; **, $p<0.01$; ***, $p<0.001$. Scale bar: 100 μm in (C).

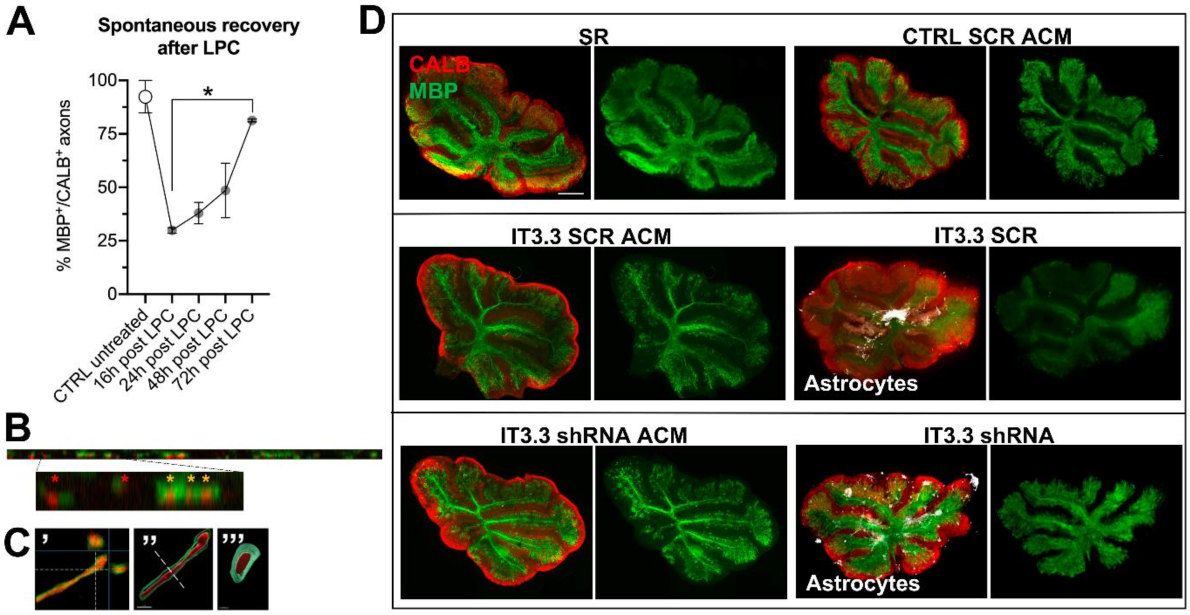


**Supplementary Figure 5. Dynamics of MBP/CALB-associated myelin recovery and whole-slice myelination pattern after LPC exposure.** (A) Time course of the spontaneous recovery (SR) of myelination occurring after LPC application on cerebellar organotypic cultures. (B) Z projection of the white dotted line in SR, shown as an example of the used approach. Higher-magnification images of the Z projection are shown to provide examples of CALB axons scored as MBP associated (yellow asterisks) or not (red asterisks). (C) Representative 2D (′) and 3D (″, lateral view; ‴, transverse view) reconstruction of a myelinated $CALB^+$ axon. (D) Representative whole-slice images of cerebellar organotypic cultures stained for CALB (red) and MBP (green) in SR, CTRL SCR/IT3.3 SCR/shRNA ACM (panels I-IV), and in IT3.3 SCR/shRNA (seeded, panels V, VI) conditions. Seeded $GFP^+$ astrocytes are in white. LPC, lysolecithin; SR, Spontaneous Recovery. RM One-way ANOVA, Tukey's multiple comparison (A; n=2 independent experiments). *, $p<0.05$. Scale bar: 3 μm in (C), 500 μm in (D).
