## Supplementary Table 1 for "Functional validation of allele-specific LMNB1 silencing in patient-derived astrocytes as a therapeutic option for Autosomal Dominant Leukodystrophy"

**Supplementary Table 1. Bulk RNAseq**

**Up-regulated Genes**

| Gene | log2FoldChange | padj |
|---|---|---|
| DPP6 | 8,443291958 | 0 |
| DNAH9 | 6,503021557 | 0 |
| NNAT | 8,44240758 | 0 |
| CES1 | 6,54669916 | 0 |
| PTPRT | 7,403562734 | 0 |
| POU3F4 | 6,735339608 | 0 |
| LOC728488 | 9,250445621 | 0 |
| CBS | 2,024821358 | 0 |
| CLDN6 | 2,941188321 | 0,000006 |
| NPIPB15 | 4,244362904 | 0,000172 |
| SHISA6 | 6,179293281 | 0,000216 |
| PCDHA11 | 2,809357503 | 0,000414 |
| ELF3 | 3,218429674 | 0,000620 |
| MAFIP | 6,185232502 | 0,000687 |
| FBLIM1 | 1,295538266 | 0,001081 |
| SYK | 4,499731346 | 0,001081 |
| FOXC1 | 1,461665612 | 0,001440 |
| RBM14-RBM4 | 1,974144659 | 0,001610 |
| MAFIP | 5,601812639 | 0,002167 |
| FOXD2-AS1 | 2,788754468 | 0,002293 |
| MGP | 2,276021556 | 0,002429 |
| SLC1A5 | 1,097053774 | 0,003381 |
| BLOC1S3 | 0,8173212357 | 0,003381 |
| VIRMA | 0,589313801 | 0,003408 |
| DNLZ | 0,8300664721 | 0,003682 |
| GUSBP14 | 1,303131554 | 0,003847 |
| DLK2 | 1,46506954 | 0,004602 |
| SCUBE2 | 1,153915727 | 0,005946 |
| CAT | 0,9215795372 | 0,007845 |
| ANK1 | 2,14299685 | 0,008297 |
| KHDRBS3 | 1,192160776 | 0,009724 |
| TNFRSF10A | 1,548403958 | 0,010774 |
| TMEM40 | 4,371792579 | 0,011846 |
| F2RL1 | 1,181569264 | 0,015654 |
| PDZD2 | 1,424723883 | 0,015771 |
| ADAMTSL3 | 2,598767822 | 0,016651 |
| TM4SF1 | 1,18024882 | 0,017440 |
| TMEM54 | 1,103971039 | 0,019970 |
| COL9A3 | 1,409488885 | 0,019970 |
| COL4A2 | 0,7769313493 | 0,019981 |
| HOXB9 | 1,759572088 | 0,021141 |
| HOXB6 | 1,417824725 | 0,021281 |
| SHF | 0,7166390089 | 0,022335 |
| SLC12A8 | 1,467361989 | 0,022487 |
| WNK4 | 1,401030706 | 0,023005 |
| ZNF726 | 2,463965126 | 0,027091 |
| IL4R | 1,247828811 | 0,031657 |
| GYPC | 1,128173763 | 0,031667 |
| HOXB8 | 1,228513881 | 0,034063 |
| OCLNP1 | 3,757604563 | 0,035299 |
| MSC-AS1 | 1,804469743 | 0,035299 |
| THY1 | 1,189100302 | 0,035728 |
| CAV1 | 1,047757443 | 0,036150 |
| WTIP | 0,7195533637 | 0,040331 |
| IGFBP3 | 1,344933309 | 0,040413 |
| ITGA3 | 0,9174178655 | 0,040936 |
| SHMT2 | 0,6206422951 | 0,042831 |
| NUP42 | 0,6592541996 | 0,045825 |
| COL13A1 | 1,060884633 | 0,045963 |
| SIGIRR | 0,9627320849 | 0,046788 |
| HOXB7 | 1,236648839 | 0,047069 |
| ARSJ | 1,084854128 | 0,048031 |
| CAV2 | 0,9522062306 | 0,048579 |
| TFPT | 0,6393871701 | 0,048579 |
| CCN4 | 1,752722841 | 0,049020 |
| PYCR1 | 0,7766556212 | 0,050539 |
| KIT | 1,246051067 | 0,051455 |

**Down-regulated Genes**

| Gene | log2FoldChange | padj |
|---|---|---|
| CBSL | -4,796089642 | 0 |
| RPS14P1 | -6,148323307 | 0 |
| SNHG10 | -2,037763307 | 0 |
| FUT9 | -2,189936265 | 0,000104 |
| CPS1 | -1,801033303 | 0,000218 |
| FGD4 | -1,079726504 | 0,000408 |
| PKD1P1 | -1,478469317 | 0,000620 |
| CAMK2B | -1,605053449 | 0,000697 |
| RPSAP58 | -1,70116534 | 0,000750 |
| FKBP9P1 | -2,421230783 | 0,000763 |
| PDCD6P1 | -1,71056184 | 0,000784 |
| THNSL2 | -1,47974588 | 0,001076 |
| RMST | -1,090115844 | 0,003845 |
| GRIN2A | -1,873899693 | 0,004660 |
| GRM5 | -2,040705628 | 0,005167 |
| PDE8B | -1,197848146 | 0,006185 |
| TRDN | -1,878399283 | 0,007320 |
| GTF2IRD2 | -1,049781184 | 0,009537 |
| TMEM74B | -1,968277458 | 0,010321 |
| ALG12 | -0,7829677167 | 0,010435 |
| FOLR1 | -1,568296681 | 0,010774 |
| LCA5L | -1,836203723 | 0,011161 |
| RGPD8 | -1,044088977 | 0,015185 |
| LYNX1 | -1,410614246 | 0,015512 |
| FAM238C | -1,924684481 | 0,015654 |
| WASH7P | -0,8785847043 | 0,016108 |
| CAPN3 | -1,585132953 | 0,016591 |
| P2RY1 | -1,262624842 | 0,017886 |
| SLC35E2A | -0,9631206872 | 0,019838 |
| LINC01001 | -1,363424691 | 0,020195 |
| NMRK1 | -1,194058936 | 0,020988 |
| COL4A5 | -0,762023573 | 0,021520 |
| AMH | -1,343317883 | 0,027091 |
| DNAH1 | -1,147404008 | 0,027091 |
| ACACB | -0,9245347351 | 0,027091 |
| GMDS-DT | -0,779361557 | 0,028302 |
| PRR22 | -1,783839306 | 0,030319 |
| LY6H | -0,9384811276 | 0,031657 |
| SUGT1P4-STR | -0,835678912 | 0,031667 |
| HERC2P2 | -1,106824412 | 0,032693 |
| EFCC1 | -0,99288496 | 0,035728 |
| ZC2HC1C | -0,9021295124 | 0,039537 |
| NT5M | -0,8756001596 | 0,039537 |
| POMZP3 | -0,8268426736 | 0,039616 |
| TOGARAM2 | -1,28778875 | 0,040102 |
| GPM6B | -0,8881226013 | 0,040413 |
| CFAP46 | -0,7945713372 | 0,040413 |
| CSRNP3 | -1,03214176 | 0,044891 |
| GPR35 | -0,9941641302 | 0,045963 |
| SLC47A1 | -1,325392527 | 0,048031 |
| TRIM52-AS1 | -0,6530353746 | 0,048579 |
| LRAT | -1,060304611 | 0,050439 |
| AQP4 | -1,014435711 | 0,050530 |
| IFI27 | -1,195752696 | 0,050593 |
| ADORA2A | -0,9139699168 | 0,050593 |
| LINC00467 | -0,8744652566 | 0,050593 |
| TDRP | -0,6072916926 | 0,051851 |
| ERBB4 | -0,7708833024 | 0,052767 |
