## Supplementary Table 2 for "Functional validation of allele-specific LMNB1 silencing in patient-derived astrocytes as a therapeutic option for Autosomal Dominant Leukodystrophy"

**Supplementary Table 2 A. GO Analysis - Biological Process**

| ID | Description | BgRatio | pvalue | p.adjust | qvalue | geneID |
|---|---|---|---|---|---|---|
| GO:0051480 | regulation of cytosolic calcium ion concentration | 373/21081 | 0,0000005007493053 | 0,001200296085 | 0,001081091395 | RYR2/TRDN/GRIN2A/GPR35/P2RY1/GRM5/F2RL1/THY1/GRM1/CAV1/CAV2/CAPN3/ATP2B3 |
| GO:0055074 | calcium ion homeostasis | 496/21081 | 0,00001138362433 | 0,00720498272 | 0,006489436163 | RYR2/TRDN/GRIN2A/GPR35/P2RY1/GRM5/F2RL1/THY1/GRM1/CAV1/CAV2/CAPN3/ATP2B3 |
| GO:1904062 | regulation of cation transmembrane transport | 365/21081 | 0,00001550835939 | 0,00720498272 | 0,006489436163 | VAMP2/RYR2/SHISA6/TRDN/GRIN2A/GPR35/THY1/DPP6/WNK4/CAV1/CAPN3 |
| GO:0046942 | carboxylic acid transport | 368/21081 | 0,0000167303161 | 0,00720498272 | 0,006489436163 | SLC35D2/VAMP2/CES1/LCN12/SYK/SLC47A1/FOLR1/SLC1A5/ACACB/SLC11A1/NOS2 |
| GO:0015849 | organic acid transport | 371/21081 | 0,00001803500055 | 0,00720498272 | 0,006489436163 | SLC35D2/VAMP2/CES1/LCN12/SYK/SLC47A1/FOLR1/SLC1A5/ACACB/SLC11A1/NOS2 |
| GO:0051712 | positive regulation of killing of cells of other organism | 11/21081 | 0,00003563511497 | 0,01220248151 | 0,01099061967 | SYK/F2RL1/NOS2 |
| GO:0006563 | L-serine metabolic process | 12/21081 | 0,00004730091632 | 0,01239257523 | 0,01116183466 | CBSL/SHMT2/CBS |
| GO:0032412 | regulation of ion transmembrane transporter activity | 276/21081 | 0,00005170035559 | 0,01239257523 | 0,01116183466 | VAMP2/RYR2/SHISA6/TRDN/GRIN2A/GPR35/GRM5/WNK4/CAV1 |
| GO:0009071 | serine family amino acid catabolic process | 13/21081 | 0,00006121615926 | 0,01327990959 | 0,01196104541 | SHMT2/CBS/THNSL2 |
| GO:0030198 | extracellular matrix organization | 442/21081 | 0,00008798443618 | 0,01344739964 | 0,01211190157 | COL13A1/COL4A5/ELF3/ADAMTSL3/COL4A2/CAV1/CAV2/COL9A3/FOXC1/GPM6B/ITGA3 |
| GO:0043062 | extracellular structure organization | 443/21081 | 0,00008976153285 | 0,01344739964 | 0,01211190157 | COL13A1/COL4A5/ELF3/ADAMTSL3/COL4A2/CAV1/CAV2/COL9A3/FOXC1/GPM6B/ITGA3 |
| GO:0032409 | regulation of transporter activity | 302/21081 | 0,0001029424069 | 0,01451487937 | 0,01307336697 | VAMP2/RYR2/SHISA6/TRDN/GRIN2A/GPR35/GRM5/WNK4/CAV1 |
| GO:0038063 | collagen-activated tyrosine kinase receptor signaling pathway | 18/21081 | 0,0001707905616 | 0,02154657769 | 0,01940672808 | COL4A5/SYK/COL4A2 |
| GO:0060401 | cytosolic calcium ion transport | 192/21081 | 0,0001826755264 | 0,02189366184 | 0,01971934235 | RYR2/TRDN/GRIN2A/THY1/CAV1/CAPN3/ATP2B3 |
| GO:0009070 | serine family amino acid biosynthetic process | 19/21081 | 0,0002019081191 | 0,02304636959 | 0,02075757154 | CBSL/SHMT2/CBS |
| GO:0010524 | positive regulation of calcium ion transport into cytosol | 54/21081 | 0,0003339697958 | 0,0363875273 | 0,03277378235 | TRDN/THY1/CAV1/CAPN3 |
| GO:0010959 | regulation of metal ion transport | 438/21081 | 0,0003689174499 | 0,03844761423 | 0,03462927642 | VAMP2/RYR2/TRDN/GPR35/THY1/DPP6/WNK4/CAV1/CAPN3/CAMK2B |
| GO:1990778 | protein localization to cell periphery | 366/21081 | 0,0004276389529 | 0,04125378573 | 0,03715675934 | VAMP2/SHISA6/GRIN2A/P2RY1/DPP6/LGI1/CAV1/ANK1/ITGA3 |
| GO:0007200 | phospholipase C-activating G protein-coupled receptor signaling pathway | 107/21081 | 0,000512443082 | 0,04661400385 | 0,04198463953 | GPR35/P2RY1/GRM5/F2RL1/GRM1 |
| GO:0086064 | cell communication by electrical coupling involved in cardiac conduction | 26/21081 | 0,0005250638732 | 0,04661400385 | 0,04198463953 | RYR2/TRDN/CAV1 |
| GO:0048705 | skeletal system morphogenesis | 234/21081 | 0,0006026642014 | 0,04981331347 | 0,04486621695 | HOXB7/COL13A1/HOXB9/HOXB8/MGP/HOXB6/FOXC1 |

**Supplementary Table 2 B. Manual Annotation of DEGs**

**UPREGULATED**

| | | | |
|---|---|---|---|
| ***CAT*** | Cellular Metabolism | Antioxidant enzyme that catalyses the decomposition of hydrogen peroxide | Layton et al., 2010 |
| ***CBS*** | Cellular Metabolism | Is the first enzyme in the transsulfuration pathway and catalyzes the conversion of serine and homocysteine to cystathionine, a substrate for cysteine synthesis | Ide et al., 2019 |
| ***CES1*** | Cellular Metabolism | Is involved in the hydrolysis of ester group-containing xenobiotic and endobiotic compounds | Gan et al., 2022 |
| ***DNLZ*** | Cellular Metabolism | Involved in mitochondrial protein import | Vu et al., 2012 |
| ***IGFBP3*** | Cellular Metabolism | Insulin-like growth factor binding protein that modulates the bioactivity and availability of insulin-like growth factor | Zhang et al., 2023 |
| ***PYCR1*** | Cellular Metabolism | Mitochondrial enzyme that catalyzes the NAD(P)H-dependent conversion of pyrroline-5-carboxylate to proline | Alaqbi et al., 2022 |
| ***SHMT2*** | Cellular Metabolism | Mitochondrial form of a pyridoxal phosphate-dependent enzyme that catalyzes the reversible reaction of serine and tetrahydrofolate to glycine and 5,10-methylene tetrahydrofolate, primarily responsible for glycine synthesis | Escande-Beillard et al., 2020 |
| ***FOXC1*** | RNA Processing | Transcription factor | Mishra et al., 2016 |
| ***KHDRBS3*** | RNA Processing | Involved in mRNA splicing | Somrit et al., 2024 |
| ***POU3F4*** | RNA Processing | Chromatin remodeling regulator | Zhang et al., 2024 |
| ***TFPT*** | RNA Processing | Chromatin remodeling regulator | Irie et al., 2000 |
| ***VIRMA*** | RNA Processing | Involved in mRNA splicing | Zou et al., 2024 |
| ***WTIP*** | RNA Processing | Gene transcriptional regulators | Hou et al., 2010 |
| ***ZNF726*** | RNA Processing | Gene transcriptional regulators | Chang et al., 2023 |
| ***CCN4*** | Immune Response | Secreted cysteine-rich matricellular protein produced by activating the Wnt/β-catenin pathway | Wang et al., 2012 |
| ***IL4R*** | Immune Response | Type I cytochine receptor | Hanuscheck et al., 2022 |
| ***SIGIRR*** | Immune Response | Member of the TLR/IL-1R family and a negative regulator of the inflammation | Watson et al., 2010 |
| ***SYK*** | Immune Response | It is involved in B-cell receptor and T-cell receptor signaling | Ennerfelt et al., 2022 |

**DOWNREGULATED**

| | | | |
|---|---|---|---|
| ***CFAP46*** | Microtubule Organization | Plays a role in cilium movement | Renard et al., 2019 |
| ***DNAH1*** | Microtubule Organization | Encodes an inner arm heavy chain dynein attached to the peripheral microtubule doublets in cilia and flagellum | Wilken et al., 2024 |
| ***POMZP3*** | Microtubule Organization | Nuclear pore membrane protein | Ge et al., 2019 |
| ***TOGARAM2*** | Microtubule Organization | Involved in microtubule cytoskeleton organization | Patterson et al., 2024 |
| ***ACACB*** | Cellular Metabolism | Mitochondrial enzyme that catalyzes the carboxylation of acetyl-CoA to malonyl-CoA and plays a central role in fatty acid metabolism | Westbrook et al. 2022 |
| ***CBSL*** | Cellular Metabolism | Involved in hydrogen sulfide biosynthesis II and Peptide chain elongation | Dey et al., 2023 |
| ***CPS1*** | Cellular Metabolism | Mitochondrial enzyme that catalyzes synthesis of carbamoyl phosphate from ammonia and bicarbonate | Nissim et al. 2011 |
| ***FUT9*** | Cellular Metabolism | Is responsible for the synthesis of Lewis X carbohydrate epitope, a marker for pluripotent or multipotent tissue-specific stem cells | Abdullah et al., 2022 |
| ***NT5M*** | Cellular Metabolism | Mitochondrial enzyme that dephosphorylates the 5'- and 2'(3')-phosphates of uracil and thymine deoxyribonucleotides, protecting mitochondrial DNA replication from excess dTTP | Ramòn et al. 2021 |
