## Supplementary Table 3 for "Functional validation of allele-specific LMNB1 silencing in patient-derived astrocytes as a therapeutic option for Autosomal Dominant Leukodystrophy"

**Supplementary Table 3. GSEA Analysis - Molecular Function**

| ID | Description | setSize | enrichmentScore | NES | qvalues | core_enrichment |
|---|---|---|---|---|---|---|
| GO:0030594 | neurotransmitter receptor activity | 101 | -0,8372714629 | -1,954573577 | 0,09102121715 | GRM1/GRIN2A/GRM5 |
| GO:0008066 | glutamate receptor activity | 27 | -0,9242045677 | -1,937328918 | 0,09102121715 | GRM1/GRIN2A/GRM5 |
| GO:0030547 | receptor inhibitor activity | 27 | -0,9125167254 | -1,91282872 | 0,09102121715 | LYPD1/IGSF1/LY6H/LYNX1 |
| GO:0001637 | G protein-coupled chemoattractant receptor activity | 19 | -0,9401745547 | -1,903034794 | 0,09102121715 | CXCR4/GPR35 |
| GO:0004950 | chemokine receptor activity | 19 | -0,9401745547 | -1,903034794 | 0,09102121715 | CXCR4/GPR35 |
| GO:0016595 | glutamate binding | 10 | -0,9666612363 | -1,804484576 | 0,09102121715 | GCLC/SLC1A3/CPS1 |
| GO:0045028 | G protein-coupled purinergic nucleotide receptor activity | 10 | -0,96555292 | -1,802415661 | 0,09102121715 | GPR34/P2RY1 |
| GO:0008391 | arachidonic acid monooxygenase activity | 11 | -0,9669412739 | -1,795415615 | 0,1171519103 | CYP2C8/CYP2E1 |
| GO:0045294 | alpha-catenin binding | 10 | 0,9776914045 | 1,786577122 | 0,09102121715 | PTPRT |
| GO:0045296 | cadherin binding | 327 | 0,7770282358 | 1,78663398 | 0,09102121715 | PTPRT/EPCAM/PPL/CAPG/TJP2/CNN2/EHD1/ANXA2/TWF2/PLCB3/TBC1D10A/RPL14/LASP1/MYO1B/LRRC59/EHD4/RPL23A/PDLIM1/PFN1/DBNL/EIF4H/MYH9/IQGAP1/RARS1/ATIC/SND1/PKM/MRE11/PSMB6/RPS2/ENO1/RANGAP1/NIBAN2/SLC3A2/PLIN3/FLNA/TLN1/YKT6/EEF2/CLIC1/SH3GL1/FASN/BSG/FSCN1/CDH10/CALD1/CAST/RAB10/CAPZB/DBN1/LRRFIP1/PCBP1/KIF5B/RAN/EEF1D/RTN4/PPFIBP1/GIPC1/AHSA1/HSP90AB1/PPME1/RSL1D1/ABCF3/EZR/PFKP/ITGB1/TAGLN2/HSPA5/S100A11/KLC2/EPS8L2/EIF2A/CAPZA1/EPHA2/RANBP1/ATXN2L/CCT8/STK38/CHMP4B/RPL15/LDHA/RAB1A/CDH8/EPS8L1/NUMB/SNX5/RPL29/RACK1/SERBP1/UBFD1/NOP56/P2RX4/BZW1/VCL/RPL7A/ERC1/PSEN1/LIMA1/SPTBN2/DNAJB1/CDH2/VAPA/PLEC/NUDC/MB21D2/CDH11/RPL34/CDC42EP1/CDH1/NOTCH3/CGN/PROM1/CTNNAL1/TES/CDH13/UNC45A/SEPTIN9/MICALL1/GCN1/VASP/RPL24/EEF1G/F11R/ALDOA/GLOD4/TRPC4/BZW2/SNX9/PKP4/RPL6/CDH4/LYPLA2/TMPO/MPRIP/VAPB/PPP1CA/STK24/SWAP70/EPS15L1 |
| GO:0045295 | gamma-catenin binding | 12 | 0,9552059375 | 1,799937187 | 0,2331923115 | PTPRT |
| GO:0015459 | potassium channel regulator activity | 53 | 0,9267856845 | 1,963872222 | 0,09102121715 | DPP6/WNK4/CAV1 |
