## Supplementary Table 4 for "Functional validation of allele-specific LMNB1 silencing in patient-derived astrocytes as a therapeutic option for Autosomal Dominant Leukodystrophy"

**Supplementary Table 4 A. DETs**

**Up-regulated**

| Gene | Transcript | log2FoldChange |
|---|---|---|
| MALAT1 | ENST00000692086 | 19,3613 |
| DPP6 | ENST00000404039 | 12,0502 |
| XIST | ENST00000429829 | 10,4822 |
| MEG3 | ENST00000398461 | 8,3755 |
| XIST | ENST00000650366 | 8,2835 |
| ST14 | ENST00000278742 | 8,2403 |
| MAFIP | ENST00000400754 | 8,1878 |
| NNAT | ENST00000649451 | 7,7447 |
| DPP6 | ENST00000377770 | 7,7328 |
| NNAT | ENST00000346199 | 7,4951 |
| CMAHP | ENST00000471416 | 7,3602 |
| DNAH9 | ENST00000579828 | 7,3023 |
| PADI1 | ENST00000375471 | 7,2473 |
| PEG3 | ENST00000649428 | 7,0998 |
| MEG3 | ENST00000452120 | 7,0501 |
| HOXA11 | ENST00000006015 | 6,9417 |
| MEG3 | ENST00000398460 | 6,7828 |
| CES1 | ENST00000360526 | 6,7666 |
| MRPL42 | ENST00000552938 | 6,7461 |
| ABI1 | ENST00000376139 | 6,6549 |
| PAQR6 | ENST00000492619 | 6,6351 |
| SHISA6 | ENST00000409168 | 6,6155 |
| KRT19 | ENST00000471565 | 6,5463 |
| DNAH9 | ENST00000579406 | 6,5141 |
| IGF2BP2 | ENST00000457616 | 6,5061 |
| COL5A1 | ENST00000618395 | 6,4894 |
| PMEPA1 | ENST00000395819 | 6,4852 |
| SBF2 | ENST00000533661 | 6,451 |
| ADAMTS7P4 | ENST00000561209 | 6,4408 |
| ALKBH8 | ENST00000393100 | 6,44 |
| NEIL2 | ENST00000403422 | 6,4174 |
| CXXC1 | ENST00000587170 | 6,3419 |
| RRM2 | ENST00000652660 | 6,2918 |
| AKT2 | ENST00000578615 | 6,2781 |
| ANKRD13D | ENST00000504186 | 6,2513 |
| DCTD | ENST00000510370 | 6,2489 |
| MMP19 | ENST00000547685 | 6,2247 |
| SEC11C | ENST00000591406 | 6,2156 |
| RAD54B | ENST00000463267 | 6,1869 |
| NAXD | ENST00000470164 | 6,169 |
| SEC31A | ENST00000443462 | 6,1639 |
| MTMR9LP | ENST00000688352 | 6,1582 |
| CCDC15 | ENST00000529051 | 6,1574 |
| KIF2C | ENST00000452259 | 6,146 |
| TMEM191C | ENST00000417708 | 6,1364 |
| SRSF6 | ENST00000668808 | 6,1231 |
| NSMCE4A | ENST00000481320 | 6,123 |
| ADARB1 | ENST00000629643 | 6,1228 |
| COL20A1 | ENST00000415763 | 6,0636 |
| TPD52L1 | ENST00000527711 | 6,0619 |
| TCF7L2 | ENST00000543371 | 6,0589 |
| SMARCA4 | ENST00000644737 | 6,0371 |
| POLR1B | ENST00000537335 | 6,0324 |
| PSCA | ENST00000301258 | 6,0208 |
| NSD2 | ENST00000508299 | 6,0199 |
| INPP5F | ENST00000637174 | 6,0108 |
| PEG3 | ENST00000648694 | 5,9916 |
| USP14 | ENST00000578786 | 5,9521 |
| GRIA2 | ENST00000510854 | 5,9494 |
| LERFS | ENST00000445604 | 5,9149 |
| SULT1E1 | ENST00000226444 | 5,905 |
| PRR15L | ENST00000300557 | 5,9045 |

**Down-regulated**

| Gene | Transcript | log2FoldChange |
|---|---|---|
| ARL4A | ENST00000651779 | -1,001 |
| NPC2 | ENST00000557510 | -1,005 |
| EWSR1 | ENST00000483629 | -1,0066 |
| DBP | ENST00000222122 | -1,0077 |
| C12orf29 | ENST00000550333 | -1,0079 |
| ATG3 | ENST00000402314 | -1,0106 |
| BTN3A3 | ENST00000244519 | -1,0154 |
| PSMA6 | ENST00000554457 | -1,0156 |
| PRPS1 | ENST00000372418 | -1,018 |
| STIM2 | ENST00000477474 | -1,0185 |
| RPL7L1 | ENST00000602561 | -1,0197 |
| EOLA2-DT | ENST00000671125 | -1,0229 |
| HNRNPH1 | ENST00000442819 | -1,0246 |
| MAGI2-AS3 | ENST00000452320 | -1,0333 |
| IP6K2 | ENST00000491686 | -1,0351 |
| ZNF839 | ENST00000442396 | -1,0363 |
| NBEAL2 | ENST00000477412 | -1,0428 |
| CREBZF | ENST00000682836 | -1,0435 |
| TET2 | ENST00000305737 | -1,0467 |
| ADAMTS13 | ENST00000371929 | -1,0517 |
| TBCD | ENST00000574886 | -1,0554 |
| ANKRD36C | ENST00000295246 | -1,0573 |
| MTCH2 | ENST00000534074 | -1,061 |
| BIVM | ENST00000651808 | -1,065 |
| TMBIM4 | ENST00000544599 | -1,0655 |
| SAMD14 | ENST00000330175 | -1,0664 |
| CRTC3 | ENST00000268184 | -1,067 |
| WASH7P | ENST00000488147 | -1,0697 |
| DNAJB2 | ENST00000477917 | -1,0707 |
| WDR12 | ENST00000467777 | -1,075 |
| PTCH1 | ENST00000375290 | -1,0827 |
| TMEM94 | ENST00000585105 | -1,0839 |
| FGF13 | ENST00000315930 | -1,0847 |
| CYB5R1 | ENST00000497655 | -1,0951 |
| MDM1 | ENST00000411698 | -1,0956 |
| NUTF2 | ENST00000587481 | -1,0958 |
| ZNF276 | ENST00000289816 | -1,0979 |
| MTX1 | ENST00000368376 | -1,0998 |
| PHF21A | ENST00000686153 | -1,1135 |
| MTFR1L | ENST00000466284 | -1,1154 |
| LINC01011 | ENST00000605901 | -1,1159 |
| DDX19A | ENST00000566574 | -1,1176 |
| USP51 | ENST00000500968 | -1,1203 |
| STX16-NPEPL1 | ENST00000413559 | -1,121 |
| EOLA2-DT | ENST00000425602 | -1,1216 |
| TMCC1-DT | ENST00000605830 | -1,1222 |
| STAU2 | ENST00000522695 | -1,1225 |
| PPP1R9A | ENST00000456331 | -1,1274 |
| WHAMMP1 | ENST00000561563 | -1,1305 |
| MAN1B1 | ENST00000682881 | -1,1306 |
| BAIAP2 | ENST00000576995 | -1,132 |
| COPZ2 | ENST00000584955 | -1,1373 |
| TRMT112 | ENST00000308774 | -1,1394 |
| FLNB | ENST00000682868 | -1,151 |
| PCDHA12 | ENST00000398631 | -1,1544 |
| CAPN1 | ENST00000527897 | -1,1574 |
| LACTB2 | ENST00000276590 | -1,1618 |
| NCOR1 | ENST00000582357 | -1,1618 |
| DENND1A | ENST00000373620 | -1,1649 |
| SPARC | ENST00000524277 | -1,1651 |
| CIRBP | ENST00000593093 | -1,1705 |
| C11orf1 | ENST00000260276 | -1,1716 |

| | | |
|---|---|---|
| FAM131B | ENST00000521347 | 5,9016 |
| TBCD | ENST00000683184 | 5,8587 |
| MBNL1 | ENST00000545754 | 5,8465 |
| POLK | ENST00000506928 | 5,8429 |
| AGPAT4 | ENST00000366911 | 5,8014 |
| SCNN1A | ENST00000228916 | 5,7832 |
| UBE2V2 | ENST00000520595 | 5,7801 |
| KIFBP | ENST00000635779 | 5,7498 |
| GTF3C3 | ENST00000409364 | 5,7208 |
| C7orf57 | ENST00000539619 | 5,7197 |
| HIF1A | ENST00000539097 | 5,6982 |
| RRP1 | ENST00000471909 | 5,6773 |
| PITPNM3 | ENST00000421306 | 5,6709 |
| ABLIM3 | ENST00000506113 | 5,6704 |
| CAST | ENST00000511097 | 5,6592 |
| PNPLA6 | ENST00000646984 | 5,651 |
| PRMT1 | ENST00000610806 | 5,6477 |
| ZFYVE21 | ENST00000554255 | 5,644 |
| FMO1 | ENST00000402921 | 5,6422 |
| NOTCH1 | ENST00000680133 | 5,633 |
| GRIK2 | ENST00000683546 | 5,6277 |
| MBNL1 | ENST00000357472 | 5,6168 |
| C6orf132 | ENST00000696229 | 5,6157 |
| SEMA4B | ENST00000559247 | 5,6149 |
| PRKN | ENST00000366897 | 5,6106 |
| BRD3 | ENST00000494743 | 5,6094 |
| NPHP3-ACAD1 | ENST00000632629 | 5,6051 |
| S100A6 | ENST00000462951 | 5,5751 |
| ICA1 | ENST00000455539 | 5,5723 |
| ZNF662 | ENST00000328199 | 5,5654 |
| PPP4C | ENST00000567828 | 5,5597 |
| PROM2 | ENST00000403131 | 5,5403 |
| PMP22 | ENST00000676221 | 5,5378 |
| HINT1 | ENST00000513012 | 5,5343 |
| TK1 | ENST00000592126 | 5,5176 |
| ATP6V0D1 | ENST00000565835 | 5,507 |
| FKBP11 | ENST00000551094 | 5,5068 |
| APPL1 | ENST00000464446 | 5,4902 |
| TMEM79 | ENST00000463670 | 5,4776 |
| COL1A2 | ENST00000464916 | 5,4722 |
| MUC16 | ENST00000601404 | 5,4695 |
| SORD | ENST00000558789 | 5,4661 |
| BNC2 | ENST00000486514 | 5,4325 |
| KIDINS220 | ENST00000691349 | 5,4226 |
| PLXNA4 | ENST00000496550 | 5,4144 |
| AP2B1 | ENST00000588116 | 5,4002 |
| SLC38A2 | ENST00000547252 | 5,3907 |
| MAP4K4 | ENST00000425019 | 5,3802 |
| RAD50 | ENST00000651249 | 5,3787 |
| CCDC117 | ENST00000448492 | 5,3768 |
| SLC39A11 | ENST00000583715 | 5,374 |
| IRF3 | ENST00000377135 | 5,3732 |
| TIAM1 | ENST00000286827 | 5,3701 |
| TRAF3IP1 | ENST00000409739 | 5,3685 |
| TCF4 | ENST00000637169 | 5,3671 |
| C12orf43 | ENST00000508193 | 5,3628 |
| DHDDS | ENST00000525165 | 5,3554 |
| ABCC9 | ENST00000544039 | 5,3447 |
| OPA1 | ENST00000646793 | 5,3366 |
| PRXL2B | ENST00000444521 | 5,3305 |
| FN1 | ENST00000323926 | 5,3299 |
| CHMP2A | ENST00000600804 | 5,323 |
| S100A9 | ENST00000368738 | 5,303 |
| NFE2L3 | ENST00000606261 | 5,2985 |
| MLPH | ENST00000338530 | 5,2953 |

| | | |
|---|---|---|
| NF1 | ENST00000495910 | -1,1717 |
| CSNK1E | ENST00000494610 | -1,1727 |
| PCDHB10 | ENST00000239446 | -1,1756 |
| DCLK2 | ENST00000411937 | -1,1877 |
| EFCAB7 | ENST00000461039 | -1,1877 |
| EOLA1-DT | ENST00000618757 | -1,1971 |
| LRRC45 | ENST00000583302 | -1,1985 |
| DXO | ENST00000473976 | -1,2019 |
| LAMTOR5 | ENST00000531779 | -1,2039 |
| TSPOAP1 | ENST00000580669 | -1,2066 |
| SYT4 | ENST00000255224 | -1,2075 |
| LMAN2 | ENST00000502721 | -1,2088 |
| CCDC66 | ENST00000476142 | -1,2094 |
| LINC00476 | ENST00000660645 | -1,2134 |
| TKT | ENST00000472528 | -1,2248 |
| STX4 | ENST00000468583 | -1,2274 |
| SLC35A3 | ENST00000638929 | -1,2316 |
| PPP1R26 | ENST00000356818 | -1,2331 |
| COTL1 | ENST00000567786 | -1,2368 |
| PITPNM2 | ENST00000280562 | -1,2393 |
| MAP2 | ENST00000447185 | -1,2408 |
| CANX | ENST00000679471 | -1,2454 |
| CDC42 | ENST00000344548 | -1,2478 |
| TOB1-AS1 | ENST00000416263 | -1,2489 |
| UNC45A | ENST00000487875 | -1,2493 |
| PROSER3 | ENST00000646935 | -1,2519 |
| NELFE | ENST00000492185 | -1,2612 |
| ZNF608 | ENST00000513985 | -1,262 |
| POT1-AS1 | ENST00000449642 | -1,2687 |
| OSBP2 | ENST00000332585 | -1,2693 |
| AHI1 | ENST00000681754 | -1,2694 |
| NKPD1 | ENST00000686631 | -1,2714 |
| GREB1L | ENST00000580732 | -1,2743 |
| PLXNB1 | ENST00000470525 | -1,2744 |
| GPR85 | ENST00000610164 | -1,2769 |
| KLC1 | ENST00000380038 | -1,2771 |
| TMEM254-AS1 | ENST00000412298 | -1,2782 |
| DLG5-AS1 | ENST00000632919 | -1,2809 |
| NUDT7 | ENST00000437314 | -1,2836 |
| TP53BP1 | ENST00000476454 | -1,2859 |
| GOLGA8R | ENST00000327271 | -1,2867 |
| NUTM2D | ENST00000412718 | -1,2901 |
| CLASRP | ENST00000585615 | -1,2951 |
| ZFP41 | ENST00000520584 | -1,2962 |
| MEN1 | ENST00000377326 | -1,2963 |
| C10orf95-AS1 | ENST00000473970 | -1,2967 |
| NSMF | ENST00000371468 | -1,3027 |
| HDHD2 | ENST00000588183 | -1,3054 |
| MLH1 | ENST00000673741 | -1,3083 |
| C4orf47 | ENST00000378850 | -1,3115 |
| IRS4-AS1 | ENST00000436013 | -1,3118 |
| DPH7 | ENST00000479650 | -1,3198 |
| GBF1 | ENST00000678486 | -1,3213 |
| BABAM1 | ENST00000594247 | -1,3236 |
| RMST | ENST00000663411 | -1,3258 |
| SAFB | ENST00000591991 | -1,326 |
| ETFBKMT | ENST00000357721 | -1,3281 |
| LYRM2 | ENST00000524153 | -1,3295 |
| TRIM52-AS1 | ENST00000507434 | -1,3362 |
| GET1 | ENST00000466787 | -1,3364 |
| KCNA5 | ENST00000252321 | -1,341 |
| REPS1 | ENST00000492787 | -1,3582 |
| POMGNT1 | ENST00000690377 | -1,3586 |
| RAB7A | ENST00000675712 | -1,3615 |
| GABARAP | ENST00000571129 | -1,3682 |

| | | |
|---|---|---|
| TSPY26P | ENST00000476365 | 5,2942 |
| MEG3 | ENST00000522771 | 5,2858 |
| MYEOV | ENST00000441339 | 5,2759 |
| SMTN | ENST00000612341 | 5,272 |
| AKR7A2 | ENST00000481966 | 5,268 |
| SLC16A4 | ENST00000437429 | 5,267 |
| ITGA11 | ENST00000423218 | 5,2668 |
| GAL3ST1 | ENST00000402369 | 5,2631 |
| SRSF11 | ENST00000460795 | 5,2587 |
| DNAJC17 | ENST00000560645 | 5,2516 |
| ATR | ENST00000666943 | 5,2465 |
| HAP1 | ENST00000455021 | 5,2293 |
| BCL3 | ENST00000444487 | 5,2282 |
| ACADM | ENST00000680743 | 5,2275 |
| NKIRAS2 | ENST00000587337 | 5,2201 |
| S100A13 | ENST00000392622 | 5,213 |
| POU3F4 | ENST00000644024 | 5,2014 |
| PKD1 | ENST00000564313 | 5,1909 |
| XPC | ENST00000452172 | 5,1899 |
| LAMB1 | ENST00000677994 | 5,1746 |
| SLC39A11 | ENST00000579732 | 5,1746 |
| NDE1 | ENST00000674588 | 5,1675 |
| ABTB1 | ENST00000468137 | 5,1572 |
| ESRP2 | ENST00000565858 | 5,148 |
| PTHLH | ENST00000545234 | 5,1473 |
| STAU2 | ENST00000518981 | 5,1463 |
| GK5 | ENST00000472759 | 5,1283 |
| DNAH9 | ENST00000262442 | 5,1272 |
| PITRM1 | ENST00000677001 | 5,1253 |
| DRAM2 | ENST00000461449 | 5,1234 |
| LRRC37B | ENST00000578674 | 5,1136 |
| LINC02210 | ENST00000592428 | 5,1069 |
| MRPL27 | ENST00000508200 | 5,0935 |
| DYSF | ENST00000258104 | 5,085 |
| SPG7 | ENST00000645886 | 5,0831 |
| LRRC34 | ENST00000522526 | 5,0827 |
| RSRC1 | ENST00000480119 | 5,0716 |
| ATXN2L | ENST00000564162 | 5,0674 |
| ITSN2 | ENST00000427234 | 5,065 |
| MYLK | ENST00000687709 | 5,0585 |
| ASAP2 | ENST00000484590 | 5,0506 |
| VCAN | ENST00000502527 | 5,0427 |
| NRXN1 | ENST00000401669 | 5,0392 |
| RBM17 | ENST00000418631 | 5,0295 |
| MTCH1 | ENST00000695068 | 5,0243 |
| DICER1-AS1 | ENST00000692703 | 5,0231 |
| HYDIN2 | ENST00000650199 | 5,0141 |
| USP14 | ENST00000400266 | 5,0113 |
| HTR2B | ENST00000258400 | 5,0089 |
| ZCCHC4 | ENST00000505412 | 5,0084 |
| CNN2 | ENST00000565096 | 5,0045 |
| CMKLR1 | ENST00000412676 | 4,9967 |
| PHF11 | ENST00000426879 | 4,9773 |
| TRIM2 | ENST00000676458 | 4,9705 |
| LINC01141 | ENST00000693117 | 4,9582 |
| HGS | ENST00000677109 | 4,955 |
| CREB3L2 | ENST00000456390 | 4,943 |
| LINC00877 | ENST00000668168 | 4,9415 |
| GUK1 | ENST00000366721 | 4,9389 |
| MAP3K7CL | ENST00000545939 | 4,9181 |
| PLAGL1 | ENST00000625622 | 4,9121 |
| KRT15 | ENST00000254043 | 4,912 |
| SERPINB2 | ENST00000299502 | 4,9118 |
| CD44 | ENST00000526669 | 4,9108 |
| AMIGO2 | ENST00000429635 | 4,9032 |

| | | |
|---|---|---|
| LRRC51 | ENST00000642510 | -1,3688 |
| BPTF | ENST00000582467 | -1,3737 |
| ISCU | ENST00000539593 | -1,375 |
| COQ7 | ENST00000561858 | -1,3774 |
| CDAN1 | ENST00000563604 | -1,3777 |
| PRRT2 | ENST00000567659 | -1,3798 |
| NAPA | ENST00000597274 | -1,3927 |
| PCMTD1 | ENST00000521344 | -1,3937 |
| CFAP58 | ENST00000369704 | -1,3994 |
| BAG6 | ENST00000439687 | -1,4014 |
| BSCL2 | ENST00000470529 | -1,4026 |
| PARD3B | ENST00000622699 | -1,4117 |
| CELSR3 | ENST00000498057 | -1,4118 |
| SMG1P3 | ENST00000522841 | -1,4129 |
| TMEM68 | ENST00000522576 | -1,4132 |
| ZNF213-AS1 | ENST00000654482 | -1,4134 |
| FAM184A | ENST00000475529 | -1,4155 |
| YPEL4 | ENST00000524592 | -1,4226 |
| SLC36A4 | ENST00000527503 | -1,4227 |
| ADRB2 | ENST00000305988 | -1,4263 |
| PIGP | ENST00000399098 | -1,4282 |
| BEX2 | ENST00000372677 | -1,429 |
| FBXO48 | ENST00000377957 | -1,4294 |
| ZNF140 | ENST00000536790 | -1,4348 |
| KIF5A | ENST00000675433 | -1,4392 |
| BTD | ENST00000417015 | -1,4457 |
| PPIL3 | ENST00000392283 | -1,4492 |
| UROS | ENST00000650472 | -1,4494 |
| STARD3 | ENST00000584850 | -1,4507 |
| SBF1 | ENST00000689177 | -1,4555 |
| IVD | ENST00000491554 | -1,4607 |
| SMUG1 | ENST00000682136 | -1,4614 |
| HEY1 | ENST00000523531 | -1,4679 |
| WASHC2C | ENST00000537517 | -1,4689 |
| GOLGA3 | ENST00000545875 | -1,4714 |
| NPIPB9 | ENST00000357796 | -1,4723 |
| EEF1AKMT2 | ENST00000652548 | -1,4752 |
| MRPS25 | ENST00000253686 | -1,4811 |
| NEPRO | ENST00000486271 | -1,4811 |
| NKD2 | ENST00000274150 | -1,4844 |
| TRIM2 | ENST00000676305 | -1,4855 |
| TPGS1 | ENST00000588278 | -1,4964 |
| WASH5P | ENST00000631796 | -1,4967 |
| KAT6B | ENST00000649463 | -1,5021 |
| PANTR1 | ENST00000661120 | -1,503 |
| SENP7 | ENST00000394091 | -1,5051 |
| DNAJC7 | ENST00000674337 | -1,5122 |
| CEP70 | ENST00000484888 | -1,517 |
| RNH1 | ENST00000534797 | -1,5196 |
| CFAP92 | ENST00000645291 | -1,5203 |
| GEM | ENST00000396194 | -1,5238 |
| SERPING1 | ENST00000677856 | -1,5284 |
| CLYBL | ENST00000376355 | -1,5299 |
| NTN5 | ENST00000270235 | -1,5341 |
| SDHAP3 | ENST00000652578 | -1,5351 |
| MORF4L1 | ENST00000558746 | -1,5352 |
| ACSF3 | ENST00000614302 | -1,5373 |
| SPAG8 | ENST00000495667 | -1,5382 |
| CTSK | ENST00000678725 | -1,5389 |
| TMEM94 | ENST00000578624 | -1,5422 |
| DYNC2LI1 | ENST00000482738 | -1,5423 |
| AKNA | ENST00000312033 | -1,5472 |
| DEPTOR | ENST00000286234 | -1,5498 |
| CASC2 | ENST00000454857 | -1,5523 |
| NGLY1 | ENST00000493324 | -1,5563 |

| | | |
|---|---|---|
| NBN | ENST00000613033 | 4,9029 |
| MINK1 | ENST00000453408 | 4,9012 |
| WIZ | ENST00000673675 | 4,8999 |
| LARP1B | ENST00000507377 | 4,8978 |
| SLCO4A1 | ENST00000217159 | 4,8967 |
| MFSD9 | ENST00000421966 | 4,8836 |
| GINS1 | ENST00000696808 | 4,874 |
| TNS4 | ENST00000254051 | 4,8632 |
| XPO1 | ENST00000676771 | 4,8576 |
| LRPPRC | ENST00000682885 | 4,8482 |
| ACO2 | ENST00000679264 | 4,8443 |
| RGS11 | ENST00000359740 | 4,8438 |
| ANKRD13B | ENST00000488766 | 4,8349 |
| FERMT1 | ENST00000478194 | 4,834 |
| NOC4L | ENST00000535343 | 4,829 |
| ERGIC3 | ENST00000416206 | 4,8265 |
| PRSS8 | ENST00000317508 | 4,8233 |
| DHX37 | ENST00000544745 | 4,8171 |
| GRP | ENST00000256857 | 4,8163 |
| DCAF8 | ENST00000475733 | 4,8067 |
| TRAPPC9 | ENST00000648948 | 4,802 |
| ZNF330 | ENST00000506302 | 4,7992 |
| PID1 | ENST00000409462 | 4,7988 |
| TMEM107 | ENST00000417073 | 4,7965 |
| PCDHGA7 | ENST00000518325 | 4,7927 |
| TNK2 | ENST00000672548 | 4,7918 |
| MTFR1L | ENST00000526894 | 4,7542 |
| PYCR1 | ENST00000584848 | 4,7533 |
| PHAX | ENST00000513813 | 4,7499 |
| ODF2 | ENST00000688016 | 4,7383 |
| HNF1B | ENST00000617811 | 4,7305 |
| ADAM15 | ENST00000360674 | 4,7272 |
| LINC01638 | ENST00000418271 | 4,7247 |
| PLK2 | ENST00000617412 | 4,7182 |
| LHX8 | ENST00000356261 | 4,7175 |
| VRK2 | ENST00000412104 | 4,7167 |
| RRP1 | ENST00000475534 | 4,7159 |
| FKBP15 | ENST00000689480 | 4,7099 |
| ARHGAP33 | ENST00000588248 | 4,7043 |
| ZC3H14 | ENST00000318308 | 4,7022 |
| BIN3 | ENST00000522687 | 4,7002 |
| DEPDC7 | ENST00000427755 | 4,6967 |
| MKS1 | ENST00000393120 | 4,6886 |
| BASP1-AS1 | ENST00000667677 | 4,6863 |
| SMYD2 | ENST00000460580 | 4,6855 |
| COQ8B | ENST00000677517 | 4,6848 |
| RPLP0P2 | ENST00000490750 | 4,6785 |
| GPR17 | ENST00000486700 | 4,6729 |
| STON1-GTF2A | ENST00000470560 | 4,6611 |
| NECAP1 | ENST00000450991 | 4,6572 |
| EPS8L1 | ENST00000201647 | 4,6547 |
| WIPF1 | ENST00000679041 | 4,6534 |
| BDKRB2 | ENST00000542454 | 4,6463 |
| BFAR | ENST00000566710 | 4,644 |
| CHRAC1 | ENST00000518971 | 4,6375 |
| KLK10 | ENST00000358789 | 4,6346 |
| CTSV | ENST00000681330 | 4,6343 |
| TARS2 | ENST00000369054 | 4,6312 |
| TUBGCP2 | ENST00000368562 | 4,6299 |
| NOL11 | ENST00000581966 | 4,6296 |
| CDK5RAP2 | ENST00000688923 | 4,6137 |
| TTC37 | ENST00000508181 | 4,6087 |
| ZBED2 | ENST00000317012 | 4,5962 |
| HOXC6 | ENST00000394331 | 4,5958 |
| COA1 | ENST00000395879 | 4,5957 |

| | | |
|---|---|---|
| MIR29B2CHG | ENST00000608023 | -1,5576 |
| PARP10 | ENST00000526007 | -1,5579 |
| ERCC3 | ENST00000462306 | -1,56 |
| EIF4H | ENST00000678438 | -1,5609 |
| MFSD10 | ENST00000503596 | -1,5622 |
| SNED1 | ENST00000310397 | -1,5691 |
| EPS8 | ENST00000645775 | -1,5701 |
| TTLL4 | ENST00000465558 | -1,5704 |
| PABPC4 | ENST00000677548 | -1,5706 |
| MTLN | ENST00000611969 | -1,576 |
| TMEM70 | ENST00000517439 | -1,5764 |
| DDR1 | ENST00000504152 | -1,5797 |
| BRWD1-AS2 | ENST00000603064 | -1,5802 |
| LHFPL1 | ENST00000371968 | -1,5812 |
| SUOX | ENST00000394109 | -1,5819 |
| SPG7 | ENST00000647476 | -1,5939 |
| SERF1B | ENST00000380750 | -1,5962 |
| BAZ2A | ENST00000549763 | -1,5969 |
| ECI2 | ENST00000380120 | -1,5974 |
| PICALM | ENST00000528398 | -1,5985 |
| CAST | ENST00000421689 | -1,6004 |
| PDE8B | ENST00000264917 | -1,6049 |
| YJU2B | ENST00000540216 | -1,6071 |
| YPEL5 | ENST00000495673 | -1,6105 |
| KDM1A | ENST00000602503 | -1,6183 |
| CARS2 | ENST00000375781 | -1,6202 |
| LINC01001 | ENST00000526704 | -1,6237 |
| CDC25C | ENST00000503022 | -1,6255 |
| FAM131A | ENST00000639617 | -1,6255 |
| GTF2IRD2B | ENST00000611835 | -1,6259 |
| SREK1 | ENST00000522214 | -1,6292 |
| GTPBP10 | ENST00000257659 | -1,6293 |
| HHLA3 | ENST00000463058 | -1,6394 |
| SPG7 | ENST00000562775 | -1,6401 |
| JAM2 | ENST00000471689 | -1,6412 |
| RINL | ENST00000593424 | -1,6418 |
| LIPT1 | ENST00000651691 | -1,6463 |
| WDR54 | ENST00000482605 | -1,6475 |
| DYNC2H1 | ENST00000533027 | -1,6477 |
| RAB3GAP2 | ENST00000688035 | -1,6499 |
| ZNF286A | ENST00000481540 | -1,6549 |
| FAM219A | ENST00000379081 | -1,656 |
| TNPO1 | ENST00000680533 | -1,6587 |
| PAK1 | ENST00000525542 | -1,66 |
| INTS6-AS1 | ENST00000601034 | -1,6627 |
| PTGDR2 | ENST00000332539 | -1,6636 |
| IST1 | ENST00000545388 | -1,6637 |
| TTC34 | ENST00000401095 | -1,6637 |
| FAM214A | ENST00000562351 | -1,6681 |
| KLC4 | ENST00000467906 | -1,6744 |
| CCDC144NL-AS1 | ENST00000692309 | -1,6745 |
| TCEAL7 | ENST00000332431 | -1,6751 |
| TSPOAP1-AS1 | ENST00000667382 | -1,6752 |
| BBS1 | ENST00000527959 | -1,6826 |
| DECR2 | ENST00000465166 | -1,686 |
| NEPRO | ENST00000474311 | -1,6864 |
| RAB33B-AS1 | ENST00000610159 | -1,6877 |
| ACADVL | ENST00000583312 | -1,6886 |
| HYOU1 | ENST00000694928 | -1,6972 |
| CXCL2 | ENST00000508487 | -1,6973 |
| HECTD1 | ENST00000556224 | -1,6975 |
| C16orf95-DT | ENST00000602282 | -1,6983 |
| KBTBD11-AS1 | ENST00000662506 | -1,7014 |
| NRSN2 | ENST00000608736 | -1,7025 |
| ME2 | ENST00000639663 | -1,7044 |

| | | |
|---|---|---|
| ARMC9 | ENST00000682027 | 4,5875 |
| RBM14-RBM4 | ENST00000500635 | 4,5759 |
| ZMYND11 | ENST00000602682 | 4,5723 |
| SETMAR | ENST00000425863 | 4,5697 |
| CSPP1 | ENST00000677052 | 4,5693 |
| NRM | ENST00000495946 | 4,5643 |
| GATA6-AS1 | ENST00000650323 | 4,5642 |
| ANPEP | ENST00000558740 | 4,563 |
| ME2 | ENST00000382927 | 4,5582 |
| KCTD7 | ENST00000638540 | 4,5494 |
| ERC1 | ENST00000536573 | 4,5465 |
| NDUFA13 | ENST00000606722 | 4,5425 |
| ATP13A3-DT | ENST00000665891 | 4,5373 |
| PHGDH | ENST00000641811 | 4,52 |
| MARCHF8 | ENST00000476962 | 4,5182 |
| PDCD2L | ENST00000585821 | 4,5106 |
| VTCN1 | ENST00000328189 | 4,5041 |
| CRTC2 | ENST00000303569 | 4,4997 |
| CFH | ENST00000470918 | 4,4982 |
| ABCF1 | ENST00000475993 | 4,4835 |
| RNF217-AS1 | ENST00000687689 | 4,4814 |
| MEG3 | ENST00000554639 | 4,4777 |
| STOML1 | ENST00000564777 | 4,4727 |
| GDNF | ENST00000326524 | 4,4632 |
| MLH1 | ENST00000673899 | 4,4616 |
| H19 | ENST00000417089 | 4,4517 |
| ANKRD28 | ENST00000439830 | 4,4444 |
| JTB | ENST00000471173 | 4,443 |
| VPS26B | ENST00000525095 | 4,4394 |
| NAIPP1 | ENST00000504546 | 4,4371 |
| ACSS2 | ENST00000473172 | 4,4354 |
| SLC12A6 | ENST00000675289 | 4,4351 |
| MLKL | ENST00000306247 | 4,432 |
| UBL7 | ENST00000564488 | 4,431 |
| PSME3IP1 | ENST00000568671 | 4,4308 |
| SIPA1 | ENST00000394224 | 4,4303 |
| GABRA2 | ENST00000514090 | 4,4235 |
| YWHAH | ENST00000397492 | 4,4094 |
| CHMP2B | ENST00000471660 | 4,406 |
| LHX9 | ENST00000367391 | 4,402 |
| C2orf72 | ENST00000477463 | 4,3933 |
| NDUFA4L2 | ENST00000555173 | 4,3901 |
| MAN1B1 | ENST00000684336 | 4,3784 |
| FRMD8 | ENST00000531296 | 4,3733 |
| RCC2 | ENST00000474892 | 4,3731 |
| NTRK2 | ENST00000687596 | 4,373 |
| TMEM94 | ENST00000584383 | 4,3686 |
| RPL28 | ENST00000426763 | 4,3669 |
| ABHD10 | ENST00000491580 | 4,3656 |
| NR2C1 | ENST00000393101 | 4,3654 |
| EEF1A2 | ENST00000642899 | 4,3629 |
| C1RL-AS1 | ENST00000536679 | 4,3569 |
| TRAP1 | ENST00000571538 | 4,3558 |
| CCDC82 | ENST00000679960 | 4,3456 |
| MYLK | ENST00000687848 | 4,3445 |
| WDR45 | ENST00000471338 | 4,3367 |
| FBXO44 | ENST00000376760 | 4,3359 |
| DGKA | ENST00000402956 | 4,3346 |
| CDC27 | ENST00000531206 | 4,3341 |
| TOM1L1 | ENST00000572158 | 4,3308 |
| MRPS7 | ENST00000584678 | 4,3297 |
| CLGN | ENST00000414773 | 4,3269 |
| TROAP | ENST00000546776 | 4,3236 |
| POLD4 | ENST00000528087 | 4,3194 |
| NKAPD1 | ENST00000532163 | 4,3183 |

| | | |
|---|---|---|
| LYPLA1 | ENST00000618914 | -1,7065 |
| KIF13A | ENST00000358380 | -1,7164 |
| LITAF | ENST00000571976 | -1,7171 |
| EMC2 | ENST00000517593 | -1,7182 |
| CENPT | ENST00000564128 | -1,72 |
| GSTZ1 | ENST00000555208 | -1,7208 |
| SF3B2 | ENST00000529994 | -1,7211 |
| TTC1 | ENST00000682220 | -1,7253 |
| COPA | ENST00000696211 | -1,7284 |
| MHENCR | ENST00000449500 | -1,7296 |
| PNCK | ENST00000340888 | -1,73 |
| COA1 | ENST00000446564 | -1,7302 |
| KIAA0930 | ENST00000498418 | -1,7333 |
| VRK3 | ENST00000596814 | -1,7339 |
| ACAD11 | ENST00000469042 | -1,7409 |
| MARCHF8 | ENST00000453424 | -1,7474 |
| PXN-AS1 | ENST00000667017 | -1,7532 |
| FBXO41 | ENST00000295133 | -1,7541 |
| ALG11 | ENST00000649651 | -1,756 |
| RPL23AP49 | ENST00000638439 | -1,7612 |
| COL4A5 | ENST00000483338 | -1,7635 |
| LRWD1 | ENST00000485417 | -1,7676 |
| TBCD | ENST00000576603 | -1,7728 |
| CLK1 | ENST00000434813 | -1,7748 |
| EIF4G2 | ENST00000524932 | -1,7764 |
| FBF1 | ENST00000586631 | -1,7856 |
| EFTUD2 | ENST00000586276 | -1,7869 |
| ARAP1 | ENST00000544721 | -1,7882 |
| ARL4C | ENST00000339728 | -1,7897 |
| NEK6 | ENST00000373596 | -1,7908 |
| ABCD4 | ENST00000474270 | -1,7938 |
| STIM2 | ENST00000463501 | -1,7953 |
| LINC00643 | ENST00000334389 | -1,7988 |
| RARS2 | ENST00000685219 | -1,8009 |
| STRADA | ENST00000582026 | -1,8009 |
| LINC00963 | ENST00000661101 | -1,8059 |
| FCF1P2 | ENST00000415284 | -1,812 |
| PLCXD3 | ENST00000377801 | -1,8146 |
| ACBD6 | ENST00000440959 | -1,8164 |
| OR7E14P | ENST00000530490 | -1,8166 |
| GSDMC | ENST00000522273 | -1,818 |
| NT5C2 | ENST00000675164 | -1,8203 |
| LINC02256 | ENST00000693695 | -1,8261 |
| CBR1 | ENST00000530908 | -1,8262 |
| FUZ | ENST00000531017 | -1,8301 |
| CCDC187 | ENST00000638797 | -1,8321 |
| ACAD9 | ENST00000505602 | -1,8362 |
| FUBP1 | ENST00000489495 | -1,8426 |
| RPL5 | ENST00000644549 | -1,8427 |
| CHORDC1 | ENST00000530765 | -1,8441 |
| ZBTB44 | ENST00000525842 | -1,8449 |
| RUNDC3A | ENST00000225441 | -1,8453 |
| NSMAF | ENST00000427130 | -1,8455 |
| MANSC1 | ENST00000545735 | -1,8502 |
| SNHG21 | ENST00000558174 | -1,8525 |
| CTNNB1 | ENST00000643865 | -1,8579 |
| PPP6R1 | ENST00000586690 | -1,8581 |
| EIF4A1 | ENST00000584054 | -1,8587 |
| ZNF419 | ENST00000415379 | -1,8589 |
| FCGBP | ENST00000616721 | -1,8613 |
| HHAT | ENST00000261458 | -1,8613 |
| EHBP1 | ENST00000405482 | -1,8636 |
| BDNF-AS | ENST00000650703 | -1,8679 |
| SPPL3 | ENST00000545209 | -1,8718 |
| PRMT5 | ENST00000556043 | -1,879 |

| | | |
|---|---|---|
| HAGHL | ENST00000561546 | 4,3163 |
| VPS53 | ENST00000680641 | 4,3014 |
| PABPC4 | ENST00000677860 | 4,2996 |
| LIMS2 | ENST00000410011 | 4,2921 |
| HTRA2 | ENST00000484881 | 4,2704 |
| SSRP1 | ENST00000526696 | 4,2619 |
| HTD2 | ENST00000474660 | 4,2618 |
| COPS7B | ENST00000413197 | 4,2617 |
| TRIM2 | ENST00000674786 | 4,2538 |
| BDNF-AS | ENST00000530686 | 4,2536 |
| BCL3 | ENST00000487394 | 4,2527 |
| PRIM2 | ENST00000672107 | 4,2459 |
| OXA1L | ENST00000358043 | 4,245 |
| PIEZO1 | ENST00000566414 | 4,2443 |
| WASH5P | ENST00000631994 | 4,2397 |
| CDK14 | ENST00000436577 | 4,2379 |
| TFPT | ENST00000391758 | 4,235 |
| ACAA1 | ENST00000484284 | 4,2346 |
| KIAA1755 | ENST00000460881 | 4,2303 |
| LYRM9 | ENST00000508862 | 4,2282 |
| THBS2 | ENST00000649844 | 4,2209 |
| CTSB | ENST00000678598 | 4,2197 |
| FZD10 | ENST00000229030 | 4,2192 |
| ZNF7 | ENST00000529819 | 4,2191 |
| MMP11 | ENST00000437086 | 4,2183 |
| PHACTR2 | ENST00000402863 | 4,218 |
| NLGN4X | ENST00000275857 | 4,2161 |
| SBF1 | ENST00000691792 | 4,2132 |
| DHRS12 | ENST00000461948 | 4,213 |
| MIDN | ENST00000591446 | 4,2129 |
| LRRC49 | ENST00000560980 | 4,2091 |
| SRP54 | ENST00000678477 | 4,1968 |
| COL9A3 | ENST00000462700 | 4,1949 |
| ZNF471 | ENST00000591537 | 4,1945 |
| NPAS3 | ENST00000357798 | 4,1901 |
| ERP44 | ENST00000691823 | 4,1861 |
| PRPSAP2 | ENST00000573253 | 4,1836 |
| ITPA | ENST00000483354 | 4,1822 |
| TMEM175 | ENST00000515492 | 4,1725 |
| PPFIBP1 | ENST00000540114 | 4,1702 |
| SUSD1 | ENST00000355396 | 4,165 |
| NKAPD1 | ENST00000280352 | 4,1618 |
| SNAI3-AS1 | ENST00000687510 | 4,1598 |
| HADHA | ENST00000643063 | 4,1524 |
| ZNF561 | ENST00000495503 | 4,149 |
| TMEM40 | ENST00000435218 | 4,1489 |
| INTS1 | ENST00000482994 | 4,1453 |
| SPATA20 | ENST00000512416 | 4,1418 |
| CSMD1 | ENST00000521646 | 4,1413 |
| CLASP1 | ENST00000452274 | 4,1387 |
| UTS2B | ENST00000340524 | 4,1368 |
| HLX | ENST00000549319 | 4,1331 |
| PPIP5K1 | ENST00000381885 | 4,1331 |
| CEP41 | ENST00000676243 | 4,1225 |
| CHKB-DT | ENST00000652967 | 4,1185 |
| POLE | ENST00000672742 | 4,118 |
| PRPF40A | ENST00000493468 | 4,1165 |
| LLGL2 | ENST00000392550 | 4,1153 |
| RPL31 | ENST00000409038 | 4,1123 |
| LCN9 | ENST00000277526 | 4,1083 |
| IFIT2 | ENST00000611722 | 4,1041 |
| PPP6R1 | ENST00000589343 | 4,1041 |
| POMT2 | ENST00000684549 | 4,0959 |
| GIT1 | ENST00000579937 | 4,0953 |
| TBX1 | ENST00000649276 | 4,0936 |

| | | |
|---|---|---|
| ABI2 | ENST00000430418 | -1,8809 |
| DIS3L2 | ENST00000433430 | -1,8814 |
| ETFA | ENST00000559075 | -1,8826 |
| RAD51C | ENST00000461271 | -1,8879 |
| L3MBTL1 | ENST00000648715 | -1,889 |
| FMO2 | ENST00000209929 | -1,8955 |
| AKR7A3 | ENST00000361640 | -1,9007 |
| WAPL-DT | ENST00000428940 | -1,905 |
| PTPN18 | ENST00000483617 | -1,9051 |
| LINC00665 | ENST00000666182 | -1,9053 |
| DNM2 | ENST00000590787 | -1,9113 |
| LRRC39 | ENST00000370137 | -1,9132 |
| ACSL6 | ENST00000379244 | -1,9176 |
| RAB35 | ENST00000534951 | -1,9176 |
| DMD | ENST00000359836 | -1,924 |
| RBBP4 | ENST00000414241 | -1,9284 |
| LAMA3 | ENST00000313654 | -1,9359 |
| DPAGT1 | ENST00000684252 | -1,9364 |
| DOC2GP | ENST00000639052 | -1,9369 |
| ZNF79 | ENST00000342483 | -1,9379 |
| REPS1 | ENST00000478483 | -1,9396 |
| ERC2 | ENST00000288221 | -1,9442 |
| SMARCC2 | ENST00000549757 | -1,9444 |
| CLASP2 | ENST00000494261 | -1,9447 |
| ITPR1 | ENST00000649144 | -1,9448 |
| PABIR3 | ENST00000644752 | -1,9449 |
| WBP2 | ENST00000587374 | -1,945 |
| MIB2 | ENST00000506488 | -1,9492 |
| RPS20 | ENST00000676918 | -1,9525 |
| MRPL46 | ENST00000558660 | -1,9538 |
| STARD4 | ENST00000509887 | -1,9563 |
| CRAT | ENST00000458362 | -1,9618 |
| CORO6 | ENST00000469090 | -1,9672 |
| PDCD6P1 | ENST00000690916 | -1,9685 |
| AHI1 | ENST00000681718 | -1,9701 |
| KDM4C | ENST00000381306 | -1,9727 |
| TMEM138 | ENST00000691720 | -1,9865 |
| GET1-SH3BGR | ENST00000647779 | -1,9911 |
| ASXL1 | ENST00000646367 | -1,9928 |
| FTO | ENST00000460382 | -1,9952 |
| AP4M1 | ENST00000422582 | -1,9974 |
| BCO2 | ENST00000460924 | -1,9987 |
| KCNH5 | ENST00000322893 | -2,0023 |
| ZGRF1 | ENST00000445413 | -2,003 |
| TMEM147-AS1 | ENST00000444728 | -2,0069 |
| RAB6D | ENST00000623617 | -2,0077 |
| MAMDC2-AS1 | ENST00000414515 | -2,011 |
| SNHG5 | ENST00000656092 | -2,0215 |
| NRBP2 | ENST00000530347 | -2,0237 |
| BEX5 | ENST00000333643 | -2,0241 |
| TNFSF13 | ENST00000380535 | -2,0358 |
| BBS5 | ENST00000392663 | -2,0433 |
| TMEM161B-DT | ENST00000667127 | -2,0572 |
| HSP90B1 | ENST00000680370 | -2,061 |
| PMPCB | ENST00000444457 | -2,0625 |
| S100PBP | ENST00000356689 | -2,0665 |
| C11orf91 | ENST00000379011 | -2,0706 |
| NMRK1 | ENST00000376808 | -2,0815 |
| TK2 | ENST00000451102 | -2,082 |
| PLA2G6 | ENST00000663895 | -2,0823 |
| PRMT3 | ENST00000330796 | -2,0852 |
| TTC8 | ENST00000556567 | -2,0852 |
| AMER2 | ENST00000357816 | -2,0889 |
| GALE | ENST00000481736 | -2,0916 |
| ARHGEF7 | ENST00000491775 | -2,0922 |

| | | |
|---|---|---|
| TTC21B | ENST00000681167 | 4,0936 |
| ADAMTSL2 | ENST00000651351 | 4,0932 |
| PTPRT | ENST00000612229 | 4,0915 |
| MIPOL1 | ENST00000396294 | 4,0904 |
| CYP2D7 | ENST00000610593 | 4,0869 |
| SGK1 | ENST00000367858 | 4,0862 |
| DEF8 | ENST00000561741 | 4,085 |
| METTL22 | ENST00000563037 | 4,0849 |
| HRAS | ENST00000479482 | 4,0847 |
| LRG1 | ENST00000306390 | 4,0837 |
| CAPZA1 | ENST00000476936 | 4,0778 |
| CCAR2 | ENST00000523349 | 4,0688 |
| OSBPL2 | ENST00000358053 | 4,0659 |
| DHCR7 | ENST00000529990 | 4,0412 |
| SSBP4 | ENST00000598159 | 4,0411 |
| ITGB4 | ENST00000584939 | 4,0402 |
| PANTR1 | ENST00000413121 | 4,0388 |
| IL32 | ENST00000525377 | 4,0369 |
| CSDE1 | ENST00000686781 | 4,0346 |
| CCDC137 | ENST00000575223 | 4,0289 |
| STING1 | ENST00000652543 | 4,0261 |
| RAMP2 | ENST00000589683 | 4,0234 |
| NDUFS2 | ENST00000678911 | 4,0224 |
| CTSB | ENST00000678929 | 4,0107 |
| AKAP8 | ENST00000537303 | 4,007 |
| SNAP23 | ENST00000349777 | 4,0018 |
| POMT2 | ENST00000555134 | 4,0012 |
| RBM14 | ENST00000460762 | 4,0008 |
| CKMT2-AS1 | ENST00000501927 | 4,0002 |
| LOXL2 | ENST00000520617 | 3,9999 |
| VEGFA | ENST00000518538 | 3,9968 |
| IRF9 | ENST00000324076 | 3,9876 |
| STING1 | ENST00000514542 | 3,9876 |
| NKX2-5 | ENST00000329198 | 3,9813 |
| E2F4 | ENST00000569573 | 3,981 |
| PRDM8 | ENST00000504452 | 3,9756 |
| CDK5RAP3 | ENST00000584525 | 3,9718 |
| FBXW7 | ENST00000604316 | 3,962 |
| CDCA7 | ENST00000695918 | 3,9603 |
| LMNB1 | ENST00000460265 | 3,9556 |
| PLCB4 | ENST00000473151 | 3,9531 |
| LARGE1 | ENST00000676126 | 3,9511 |
| CNTN4 | ENST00000397459 | 3,9495 |
| WARS1 | ENST00000557135 | 3,9475 |
| EPB41L3 | ENST00000580647 | 3,9451 |
| RSRC2 | ENST00000525332 | 3,9427 |
| SLITRK4 | ENST00000338017 | 3,9393 |
| ZNF133 | ENST00000622607 | 3,9359 |
| RPL23AP82 | ENST00000463325 | 3,9285 |
| WDR19 | ENST00000509560 | 3,9201 |
| RAPSN | ENST00000352508 | 3,9188 |
| GK5 | ENST00000460544 | 3,9174 |
| SCN1B | ENST00000596348 | 3,9096 |
| SEMA4F | ENST00000420077 | 3,9041 |
| TMEM191B | ENST00000614395 | 3,9033 |
| ST3GAL3 | ENST00000647503 | 3,9022 |
| ATF4 | ENST00000674835 | 3,9011 |
| PTCH1 | ENST00000687744 | 3,9007 |
| SLC2A11 | ENST00000345044 | 3,8973 |
| FZD10-AS1 | ENST00000509760 | 3,8893 |
| WARS1 | ENST00000555031 | 3,8883 |
| CEP120 | ENST00000675330 | 3,8817 |
| MBD1 | ENST00000347968 | 3,8801 |
| ST3GAL3 | ENST00000644922 | 3,8787 |

| | | |
|---|---|---|
| CCDC144NL-AS1 | ENST00000417232 | -2,0943 |
| CCDC144NL-AS1 | ENST00000443508 | -2,0979 |
| CTSB | ENST00000677418 | -2,0979 |
| NAA30 | ENST00000555166 | -2,0983 |
| METTL25B | ENST00000517871 | -2,0996 |
| TMEM161B-DT | ENST00000657972 | -2,1017 |
| CCDC82 | ENST00000679856 | -2,1036 |
| ZNF24 | ENST00000261332 | -2,1092 |
| TCTN1 | ENST00000680512 | -2,1112 |
| PAFAH1B1 | ENST00000675385 | -2,1156 |
| PDXK | ENST00000498040 | -2,118 |
| ERMAP | ENST00000487556 | -2,1258 |
| NADK2 | ENST00000404560 | -2,1289 |
| ZNF875 | ENST00000589218 | -2,1294 |
| KIF21A | ENST00000541463 | -2,1309 |
| B3GAT1-DT | ENST00000528482 | -2,1314 |
| SENP7 | ENST00000348610 | -2,1316 |
| SNX24 | ENST00000513881 | -2,1323 |
| EFHC1 | ENST00000481466 | -2,1361 |
| KLHL7-DT | ENST00000419813 | -2,1384 |
| OLMALINC | ENST00000690014 | -2,1416 |
| ARMCX5 | ENST00000246174 | -2,1501 |
| MID1 | ENST00000479925 | -2,1502 |
| NUMB | ENST00000556772 | -2,154 |
| ZKSCAN5 | ENST00000326775 | -2,1545 |
| CYTH1 | ENST00000589297 | -2,159 |
| DZANK1 | ENST00000377630 | -2,1667 |
| RBM23 | ENST00000556687 | -2,1667 |
| TADA1 | ENST00000467021 | -2,1732 |
| AKAP8 | ENST00000680336 | -2,1738 |
| STX18-AS1 | ENST00000670249 | -2,1775 |
| MROH1 | ENST00000326134 | -2,1783 |
| LRRC23 | ENST00000431207 | -2,1785 |
| RMST | ENST00000658877 | -2,1958 |
| NLRC5 | ENST00000688547 | -2,1961 |
| MGAT1 | ENST00000505682 | -2,1973 |
| CPT2 | ENST00000637252 | -2,1995 |
| CEP41 | ENST00000475282 | -2,2024 |
| COQ8B | ENST00000679130 | -2,2024 |
| RAB18 | ENST00000683797 | -2,2029 |
| ZBTB48 | ENST00000498342 | -2,2129 |
| RNF130 | ENST00000523487 | -2,2134 |
| GLDC | ENST00000639318 | -2,2152 |
| VPS51 | ENST00000529180 | -2,2207 |
| CC2D2B | ENST00000646931 | -2,2243 |
| TAF6 | ENST00000687641 | -2,2246 |
| DGKG | ENST00000265022 | -2,2249 |
| EXOSC10 | ENST00000485606 | -2,2356 |
| SAP30-DT | ENST00000666010 | -2,2373 |
| SUPT6H | ENST00000581510 | -2,2417 |
| PRR18 | ENST00000529616 | -2,2434 |
| BRAT1 | ENST00000421712 | -2,244 |
| CFAP92 | ENST00000669741 | -2,247 |
| SBNO1 | ENST00000267176 | -2,2479 |
| BOLA3-DT | ENST00000691938 | -2,2482 |
| CBFA2T2 | ENST00000375279 | -2,2528 |
| TEX9 | ENST00000558127 | -2,2569 |
| CCDC43 | ENST00000457422 | -2,2584 |
| SOX6 | ENST00000533870 | -2,2621 |
| JKAMP | ENST00000554721 | -2,264 |
| CYLD | ENST00000569891 | -2,2658 |
| RAB30-DT | ENST00000669459 | -2,2719 |
| MAN2C1 | ENST00000566256 | -2,2727 |
| LRRC8A | ENST00000259324 | -2,2752 |

| | | |
|---|---|---|
| NDUFA4L2 | ENST00000393825 | 3,8509 |
| C3orf62 | ENST00000479673 | 3,8447 |
| SLC12A2-DT | ENST00000668265 | 3,8425 |
| RHBDF1 | ENST00000448893 | 3,838 |
| DPF1 | ENST00000614244 | 3,8346 |
| SNX17 | ENST00000537606 | 3,8313 |
| TGFBI | ENST00000604555 | 3,8299 |
| AADAT | ENST00000353187 | 3,8284 |
| ATG7 | ENST00000435760 | 3,822 |
| LMO7 | ENST00000482116 | 3,8105 |
| ATG9B | ENST00000639579 | 3,8058 |
| SORBS2 | ENST00000492104 | 3,8025 |
| ARL6IP1 | ENST00000562234 | 3,8018 |
| PSME3 | ENST00000543428 | 3,801 |
| MIR4435-2HG | ENST00000371162 | 3,7981 |
| SERHL2 | ENST00000416156 | 3,7935 |
| STRA6 | ENST00000572975 | 3,7923 |
| ZNF160 | ENST00000429604 | 3,7915 |
| SLC18B1 | ENST00000650136 | 3,7898 |
| AKAP9 | ENST00000679821 | 3,7867 |
| DPP10 | ENST00000393147 | 3,7803 |
| BARX2 | ENST00000281437 | 3,7768 |
| ARL6IP5 | ENST00000470936 | 3,7722 |
| SNX22 | ENST00000557789 | 3,7722 |
| SLC35A3 | ENST00000640600 | 3,7672 |
| NUP160 | ENST00000694866 | 3,7654 |
| LOXL1-AS1 | ENST00000568229 | 3,757 |
| HAND2-AS1 | ENST00000502896 | 3,7532 |
| MCRS1 | ENST00000548596 | 3,7516 |
| CCDC142 | ENST00000290418 | 3,7498 |
| NEGR1 | ENST00000467479 | 3,7476 |
| HOXB7 | ENST00000467314 | 3,7417 |
| DDB1 | ENST00000680250 | 3,7389 |
| TUT7 | ENST00000375957 | 3,7381 |
| THOC5 | ENST00000472164 | 3,7282 |
| IL1RAP | ENST00000439062 | 3,7275 |
| CARS1 | ENST00000531387 | 3,7228 |
| ADAMTSL1 | ENST00000546040 | 3,7204 |
| RFESD | ENST00000511684 | 3,7182 |
| FN1 | ENST00000498719 | 3,7179 |
| ACO2 | ENST00000678688 | 3,7148 |
| MALAT1 | ENST00000618227 | 3,7103 |
| TRIM16 | ENST00000577326 | 3,709 |
| STMN2 | ENST00000518491 | 3,7088 |
| ANO4 | ENST00000392977 | 3,7075 |
| L2HGDH | ENST00000555423 | 3,6972 |
| GNAO1 | ENST00000640560 | 3,6947 |
| RHBDF2 | ENST00000590168 | 3,6905 |
| TMPRSS2 | ENST00000678743 | 3,684 |
| IDS | ENST00000490775 | 3,6838 |
| TMEM237 | ENST00000471318 | 3,6828 |
| GRN | ENST00000592783 | 3,6813 |
| PPP1R16B | ENST00000373331 | 3,6789 |
| PDLIM5 | ENST00000513341 | 3,6761 |
| TSC1 | ENST00000644097 | 3,6758 |
| DKC1 | ENST00000696587 | 3,6756 |
| CYB561A3 | ENST00000536452 | 3,6683 |
| UBXN4 | ENST00000490163 | 3,6622 |
| ATG4B | ENST00000405546 | 3,6551 |
| FLOT2 | ENST00000465427 | 3,6541 |
| ADA | ENST00000696078 | 3,6528 |
| DNAH9 | ENST00000396001 | 3,6509 |
| HEXA | ENST00000565873 | 3,6502 |
| DHRS2 | ENST00000611765 | 3,6494 |
| HSPH1 | ENST00000445273 | 3,6461 |

| | | |
|---|---|---|
| SERGEF | ENST00000528200 | -2,2767 |
| ZNF23 | ENST00000576258 | -2,2767 |
| CDKN3 | ENST00000555837 | -2,2888 |
| CINP | ENST00000541568 | -2,289 |
| CLPTM1 | ENST00000588274 | -2,2901 |
| CLASP2 | ENST00000487200 | -2,2914 |
| COQ8B | ENST00000676960 | -2,2921 |
| SYT3 | ENST00000600079 | -2,2975 |
| MTO1 | ENST00000370305 | -2,3055 |
| MID1 | ENST00000380787 | -2,3101 |
| GPNMB | ENST00000409458 | -2,3106 |
| DPP7 | ENST00000491807 | -2,3107 |
| KIFBP | ENST00000627949 | -2,3126 |
| MORC2 | ENST00000215862 | -2,3165 |
| EAF1 | ENST00000449565 | -2,3171 |
| SATB1 | ENST00000417717 | -2,3208 |
| KCTD17 | ENST00000483389 | -2,3241 |
| ALG12 | ENST00000492791 | -2,325 |
| EMC10 | ENST00000601780 | -2,3271 |
| SAMD12 | ENST00000524796 | -2,3278 |
| PDE6B-AS1 | ENST00000489312 | -2,3283 |
| CFAP276 | ENST00000369942 | -2,3294 |
| PTPN2 | ENST00000591115 | -2,3332 |
| HPS4 | ENST00000496385 | -2,3335 |
| TIAM1 | ENST00000541036 | -2,3383 |
| PCGF2 | ENST00000618506 | -2,34 |
| TP73-AS1 | ENST00000648600 | -2,3444 |
| CRIM1 | ENST00000426856 | -2,3482 |
| HDAC8 | ENST00000373554 | -2,3483 |
| CEP63 | ENST00000683138 | -2,3501 |
| SERF2 | ENST00000409617 | -2,3516 |
| SELENBP1 | ENST00000426705 | -2,3527 |
| RESF1 | ENST00000543763 | -2,3536 |
| ME2 | ENST00000640967 | -2,3589 |
| CACTIN | ENST00000248420 | -2,362 |
| ANKRD28 | ENST00000498524 | -2,3682 |
| NAV3 | ENST00000644176 | -2,369 |
| ALG11 | ENST00000680793 | -2,3698 |
| ABCB8 | ENST00000462605 | -2,3701 |
| CXCL2 | ENST00000510048 | -2,3705 |
| GGCT | ENST00000426081 | -2,3781 |
| CCDC93 | ENST00000474546 | -2,3837 |
| RNASEK | ENST00000546395 | -2,3879 |
| UBE2D3 | ENST00000503282 | -2,3895 |
| RABGAP1L | ENST00000392064 | -2,3979 |
| ELMOD3 | ENST00000466467 | -2,4003 |
| ZCCHC8 | ENST00000672018 | -2,4004 |
| E4F1 | ENST00000562589 | -2,4005 |
| CELF5 | ENST00000292672 | -2,4094 |
| TTC28-AS1 | ENST00000454996 | -2,412 |
| ZNF213-AS1 | ENST00000573447 | -2,4152 |
| MGAT5 | ENST00000468758 | -2,4215 |
| EMX2 | ENST00000616794 | -2,4235 |
| QDPR | ENST00000507439 | -2,4243 |
| ATPSCKMT | ENST00000280330 | -2,4253 |
| THNSL2 | ENST00000343544 | -2,4317 |
| POLR2C | ENST00000567982 | -2,4391 |
| ZNF235 | ENST00000291182 | -2,4392 |
| CARS2 | ENST00000535516 | -2,4394 |
| ZNF350-AS1 | ENST00000658402 | -2,4425 |
| NAA10 | ENST00000477882 | -2,4459 |
| TRDMT1 | ENST00000495022 | -2,4577 |
| ATP1B3 | ENST00000484727 | -2,4618 |
| SELENBP1 | ENST00000493560 | -2,4691 |
| SPTSSB | ENST00000620149 | -2,4754 |

| | | |
|---|---|---|
| CTSA | ENST00000678939 | 3,6448 |
| TNPO2 | ENST00000589149 | 3,643 |
| TRIM61 | ENST00000508856 | 3,6295 |
| MDK | ENST00000533952 | 3,6291 |
| IL6ST | ENST00000381287 | 3,6276 |
| MYO7B | ENST00000494959 | 3,6275 |
| NBPF26 | ENST00000620612 | 3,6265 |
| PHTF1 | ENST00000472612 | 3,6226 |
| ANKRD36 | ENST00000452478 | 3,6204 |
| FGF1 | ENST00000359370 | 3,6127 |
| THOC3 | ENST00000513006 | 3,6124 |
| CTSC | ENST00000677796 | 3,6119 |
| COL16A1 | ENST00000468459 | 3,6116 |
| OCLNP1 | ENST00000445744 | 3,6077 |
| REPIN1 | ENST00000467980 | 3,6045 |
| HADH | ENST00000640201 | 3,6012 |
| DOK5 | ENST00000491469 | 3,5965 |
| PRPF39 | ENST00000556782 | 3,5954 |
| CTSF | ENST00000677365 | 3,5939 |
| CTSC | ENST00000677106 | 3,5924 |
| ZNF692 | ENST00000495731 | 3,5922 |
| TAF6 | ENST00000692408 | 3,588 |
| SAMD11 | ENST00000622503 | 3,5872 |
| GPD2 | ENST00000496190 | 3,5867 |
| GRIA3 | ENST00000620581 | 3,5845 |
| TACSTD2 | ENST00000371225 | 3,5816 |
| SETD2 | ENST00000685505 | 3,5805 |
| FAM162A | ENST00000692876 | 3,5782 |
| ZNF668 | ENST00000539836 | 3,5713 |
| SAR1A | ENST00000373242 | 3,5652 |
| H19 | ENST00000447298 | 3,5638 |
| TAF2 | ENST00000685876 | 3,5618 |
| FYN | ENST00000491885 | 3,559 |
| KDM1B | ENST00000297792 | 3,5584 |
| SGCD | ENST00000517913 | 3,5558 |
| MTA3 | ENST00000407270 | 3,5553 |
| MYRF | ENST00000537318 | 3,5539 |
| MNT | ENST00000575374 | 3,552 |
| ATP1A1 | ENST00000440951 | 3,5503 |
| ATXN2L | ENST00000570200 | 3,5498 |
| RPS6KC1 | ENST00000366959 | 3,5421 |
| BMS1P12 | ENST00000428213 | 3,5365 |
| COPZ1 | ENST00000552218 | 3,5328 |
| PSME2 | ENST00000559056 | 3,5281 |
| GLI1 | ENST00000543426 | 3,5265 |
| SEC16B | ENST00000495165 | 3,5243 |
| TFRC | ENST00000420415 | 3,5176 |
| NKIRAS2 | ENST00000479407 | 3,513 |
| DDX20 | ENST00000680936 | 3,5122 |
| SMPD4 | ENST00000433118 | 3,5121 |
| RBM14-RBM4 | ENST00000511114 | 3,5119 |
| YIPF3 | ENST00000503972 | 3,5115 |
| RPLP0 | ENST00000228306 | 3,5108 |
| ADGRF4 | ENST00000327753 | 3,5019 |
| RAPGEFL1 | ENST00000469209 | 3,497 |
| CCDC82 | ENST00000680049 | 3,4958 |
| MGP | ENST00000545199 | 3,4933 |
| ACP1 | ENST00000439645 | 3,4905 |
| CNDP2 | ENST00000584768 | 3,49 |
| RACK1 | ENST00000512968 | 3,4894 |
| APTX | ENST00000309615 | 3,4879 |
| PLAT | ENST00000678083 | 3,4876 |
| SLC35B1 | ENST00000513508 | 3,4814 |
| ANK1 | ENST00000347528 | 3,4797 |
| TCOF1 | ENST00000513346 | 3,4796 |

| | | |
|---|---|---|
| RAF1 | ENST00000460610 | -2,4755 |
| FKBP9P1 | ENST00000455909 | -2,4787 |
| DDX51 | ENST00000329073 | -2,4814 |
| GPAA1 | ENST00000528073 | -2,4817 |
| CEP290 | ENST00000672647 | -2,4849 |
| MIR99AHG | ENST00000665504 | -2,4849 |
| CAMSAP3 | ENST00000595692 | -2,4857 |
| DRG2 | ENST00000497744 | -2,486 |
| GPM6A | ENST00000280187 | -2,4879 |
| BBOX1 | ENST00000263182 | -2,4889 |
| EPM2A-DT | ENST00000452617 | -2,4948 |
| DENND2B | ENST00000530438 | -2,4954 |
| NME2 | ENST00000514264 | -2,4977 |
| C1QTNF7 | ENST00000444304 | -2,5111 |
| ARMCX4 | ENST00000442270 | -2,5115 |
| HSPB11 | ENST00000371377 | -2,5136 |
| INTS8 | ENST00000523206 | -2,5136 |
| GRB10 | ENST00000401949 | -2,5161 |
| CCDC191 | ENST00000481358 | -2,5163 |
| CAMK2B | ENST00000457475 | -2,5181 |
| ZNF283 | ENST00000650832 | -2,5205 |
| TRIM16 | ENST00000649191 | -2,5232 |
| STAG2 | ENST00000428941 | -2,5241 |
| CCZ1B | ENST00000468078 | -2,5267 |
| TRAPPC12 | ENST00000417243 | -2,5267 |
| PCDH7 | ENST00000543491 | -2,5309 |
| BCKDHA | ENST00000538423 | -2,5409 |
| TNFRSF9 | ENST00000377507 | -2,5519 |
| MTHFR | ENST00000641909 | -2,5531 |
| NRXN3 | ENST00000335750 | -2,5542 |
| RANBP3 | ENST00000591092 | -2,5601 |
| BORCS8-MEF2B | ENST00000354191 | -2,5659 |
| KIAA1958 | ENST00000536272 | -2,5697 |
| STX18-AS1 | ENST00000507244 | -2,5713 |
| MCPH1-AS1 | ENST00000661193 | -2,5718 |
| DLGAP1 | ENST00000400150 | -2,5737 |
| PIGQ | ENST00000443147 | -2,5749 |
| PSMD6 | ENST00000475036 | -2,5773 |
| CHD4 | ENST00000646145 | -2,5783 |
| ODF3B | ENST00000463472 | -2,5809 |
| GYG2 | ENST00000381163 | -2,5817 |
| NFYC | ENST00000456393 | -2,5826 |
| PLIN4 | ENST00000633942 | -2,5833 |
| KATNAL2 | ENST00000686215 | -2,5907 |
| CA2 | ENST00000520127 | -2,5959 |
| RPL26L1 | ENST00000519239 | -2,5997 |
| ATP2B3 | ENST00000684004 | -2,6068 |
| COMMD10 | ENST00000515539 | -2,6072 |
| AFF1 | ENST00000307808 | -2,6121 |
| MRPS25 | ENST00000695388 | -2,6153 |
| C21orf58 | ENST00000472607 | -2,6161 |
| LINC00899 | ENST00000609737 | -2,6169 |
| PMP22 | ENST00000674651 | -2,6192 |
| BRF1 | ENST00000379937 | -2,6219 |
| NPIPB5 | ENST00000415654 | -2,6237 |
| DDX39A | ENST00000592927 | -2,6266 |
| NR2F1-AS1 | ENST00000607195 | -2,632 |
| H4C8 | ENST00000634956 | -2,6322 |
| NR2C2AP | ENST00000537399 | -2,6334 |
| WDR90 | ENST00000552728 | -2,6357 |
| PTPMT1 | ENST00000426530 | -2,637 |
| DXO | ENST00000478221 | -2,6382 |
| DHX58 | ENST00000251642 | -2,6388 |
| MAP9 | ENST00000311277 | -2,642 |
| FPGT | ENST00000467578 | -2,6431 |

| | | |
|---|---|---|
| CBWD1 | ENST00000314367 | 3,4794 |
| PHC2 | ENST00000486897 | 3,4783 |
| RAPGEF3 | ENST00000395360 | 3,469 |
| EFL1 | ENST00000696335 | 3,4687 |
| LTB4R2 | ENST00000533293 | 3,4643 |
| DLK2 | ENST00000372485 | 3,4592 |
| NSD2 | ENST00000382891 | 3,4582 |
| PARP10 | ENST00000524918 | 3,456 |
| GMCL1 | ENST00000468386 | 3,4542 |
| POU2F1 | ENST00000367865 | 3,4511 |
| ALKAL1 | ENST00000358543 | 3,4509 |
| DTNBP1 | ENST00000462989 | 3,4509 |
| SGCE | ENST00000646559 | 3,4265 |
| TMPRSS5 | ENST00000545579 | 3,4252 |
| CKAP2 | ENST00000490903 | 3,4236 |
| LINC00844 | ENST00000666624 | 3,4224 |
| FBN1 | ENST00000561429 | 3,4151 |
| HNRNPA2B1 | ENST00000679124 | 3,4135 |
| TNNI1 | ENST00000367312 | 3,412 |
| ACAN | ENST00000559004 | 3,4113 |
| PTBP1 | ENST00000587136 | 3,4101 |
| SLK | ENST00000474260 | 3,4076 |
| ETNK1 | ENST00000335148 | 3,4045 |
| PCNP | ENST00000470490 | 3,4031 |
| HEXA | ENST00000684602 | 3,3966 |
| PISD | ENST00000382151 | 3,395 |
| TRAP1 | ENST00000574494 | 3,3902 |
| SMC4 | ENST00000484799 | 3,384 |
| FAM76B | ENST00000545813 | 3,3833 |
| CDKL3 | ENST00000520693 | 3,3782 |
| NAA16 | ENST00000403412 | 3,3776 |
| TP63 | ENST00000354600 | 3,3768 |
| CTHRC1 | ENST00000520337 | 3,3738 |
| TK2 | ENST00000567357 | 3,3714 |
| ELP1 | ENST00000676416 | 3,3702 |
| ROBO3 | ENST00000524971 | 3,3663 |
| EMID1 | ENST00000430127 | 3,3658 |
| SERPINB7 | ENST00000398019 | 3,3647 |
| C4B | ENST00000647698 | 3,359 |
| PCOLCE2 | ENST00000470795 | 3,3568 |
| HGS | ENST00000677044 | 3,3535 |
| TMEM138 | ENST00000540194 | 3,3475 |
| ELMOD3 | ENST00000409344 | 3,3468 |
| MBP | ENST00000397865 | 3,3433 |
| KIF5C | ENST00000678133 | 3,3413 |
| TBCB | ENST00000586868 | 3,3393 |
| MED13L | ENST00000648762 | 3,3373 |
| ZNF438 | ENST00000361310 | 3,3352 |
| ANKS3 | ENST00000592190 | 3,333 |
| RNF220 | ENST00000372247 | 3,3319 |
| ANKRD22 | ENST00000371930 | 3,3289 |
| ATP2B1 | ENST00000550716 | 3,3267 |
| MX1 | ENST00000417963 | 3,3264 |
| GCNT1 | ENST00000376730 | 3,3207 |
| FAM98C | ENST00000588262 | 3,3197 |
| HRH1 | ENST00000438284 | 3,3169 |
| ATM | ENST00000452508 | 3,3154 |
| AGPS | ENST00000679994 | 3,3126 |
| METTL8 | ENST00000463392 | 3,3097 |
| COL8A1 | ENST00000463753 | 3,3082 |
| AURKB | ENST00000534871 | 3,3071 |
| ARHGAP11A | ENST00000563330 | 3,3067 |
| MRPL28 | ENST00000450882 | 3,3065 |
| HACE1 | ENST00000517424 | 3,3063 |
| FBXW4 | ENST00000664783 | 3,3042 |

| | | |
|---|---|---|
| CREB3L4 | ENST00000368600 | -2,6526 |
| SORBS2 | ENST00000480146 | -2,6561 |
| C18orf32 | ENST00000582392 | -2,6573 |
| PCCA | ENST00000636366 | -2,6647 |
| NPEPPS | ENST00000527298 | -2,6669 |
| SYNGR3 | ENST00000562045 | -2,6717 |
| GPT | ENST00000527165 | -2,6748 |
| FAM76B | ENST00000541418 | -2,6797 |
| DLG1 | ENST00000664564 | -2,683 |
| ZNF501 | ENST00000396048 | -2,6842 |
| INTS14 | ENST00000569491 | -2,6884 |
| LYNX1 | ENST00000614491 | -2,6887 |
| TMEM161B-DT | ENST00000654762 | -2,6891 |
| CTTNBP2 | ENST00000441556 | -2,6967 |
| GGA1 | ENST00000406772 | -2,6988 |
| BBS1 | ENST00000529955 | -2,7028 |
| PCMTD1-DT | ENST00000656129 | -2,7179 |
| PDCD6P1 | ENST00000689007 | -2,7246 |
| TMEM74B | ENST00000429036 | -2,725 |
| GAS5 | ENST00000693030 | -2,7259 |
| CHD9 | ENST00000561869 | -2,7281 |
| ADAMTS10 | ENST00000593913 | -2,7319 |
| SLC25A20 | ENST00000430379 | -2,7336 |
| TMEM106C | ENST00000615597 | -2,7364 |
| FKTN | ENST00000675736 | -2,7381 |
| NONO | ENST00000677014 | -2,7385 |
| TRIM66 | ENST00000646038 | -2,7481 |
| DCAF7 | ENST00000688972 | -2,7494 |
| WNT2B | ENST00000369684 | -2,7533 |
| CHCHD10 | ENST00000401675 | -2,7609 |
| CRCP | ENST00000360415 | -2,7706 |
| FNBP4 | ENST00000544590 | -2,7762 |
| HAX1 | ENST00000483970 | -2,7762 |
| MUTYH | ENST00000483127 | -2,7791 |
| SEL1L3 | ENST00000513416 | -2,7833 |
| MAMSTR | ENST00000594582 | -2,7881 |
| ZMYND11 | ENST00000402736 | -2,7884 |
| AGK | ENST00000629555 | -2,7969 |
| SNHG10 | ENST00000658932 | -2,7998 |
| USP33 | ENST00000370793 | -2,8003 |
| GTF2H2C | ENST00000512736 | -2,8034 |
| PDZK1 | ENST00000417171 | -2,8097 |
| AHI1 | ENST00000327035 | -2,8219 |
| SERPINI1 | ENST00000295777 | -2,8337 |
| NMBR | ENST00000258042 | -2,8366 |
| ASNSD1 | ENST00000607062 | -2,8479 |
| FBXO34 | ENST00000680658 | -2,848 |
| BTF3-DT | ENST00000607001 | -2,8559 |
| CNOT8 | ENST00000403027 | -2,8598 |
| CYHR1 | ENST00000526887 | -2,863 |
| HYDIN | ENST00000541601 | -2,8651 |
| CLDN2 | ENST00000540876 | -2,867 |
| C1R | ENST00000545466 | -2,8679 |
| TELO2 | ENST00000497339 | -2,8709 |
| AMDHD2 | ENST00000565963 | -2,8798 |
| TCF7 | ENST00000342854 | -2,8919 |
| SLC24A1 | ENST00000544319 | -2,8935 |
| CTSL | ENST00000676466 | -2,8947 |
| UBE2V1 | ENST00000557021 | -2,8951 |
| TAF1D | ENST00000534770 | -2,8977 |
| LINC01126 | ENST00000623515 | -2,901 |
| PGAP6 | ENST00000475348 | -2,9045 |
| ING4 | ENST00000484795 | -2,9065 |
| YKT6 | ENST00000677436 | -2,9107 |
| GGT5 | ENST00000263112 | -2,9196 |

| | | |
|---|---|---|
| IL4R | ENST00000170630 | 3,2978 |
| LYSMD4 | ENST00000479791 | 3,2973 |
| PISD | ENST00000442379 | 3,2949 |
| CEMIP | ENST00000559966 | 3,2935 |
| GCC2-AS1 | ENST00000322353 | 3,2925 |
| LSR | ENST00000354900 | 3,292 |
| PEAR1 | ENST00000469390 | 3,2918 |
| GRIN3A | ENST00000361820 | 3,2909 |
| ITGB1BP1 | ENST00000497105 | 3,2864 |
| RPL13A | ENST00000474171 | 3,2818 |
| POLE2 | ENST00000554396 | 3,2783 |
| CHEK2 | ENST00000348295 | 3,2781 |
| NMNAT2 | ENST00000464047 | 3,2773 |
| DDX54 | ENST00000546898 | 3,2722 |
| CDON | ENST00000683597 | 3,2653 |
| MTSS1 | ENST00000520094 | 3,2628 |
| STYXL1 | ENST00000359697 | 3,261 |
| LPIN3 | ENST00000632009 | 3,2601 |
| PVT1 | ENST00000661830 | 3,2561 |
| APPL2 | ENST00000539978 | 3,2532 |
| H19 | ENST00000439725 | 3,2511 |
| SRCIN1 | ENST00000622190 | 3,2487 |
| SULT1B1 | ENST00000310613 | 3,2484 |
| GRHPR | ENST00000494290 | 3,2477 |
| MTMR11 | ENST00000482025 | 3,246 |
| TAF1A-AS1 | ENST00000668515 | 3,2459 |
| SLC35F5 | ENST00000447673 | 3,245 |
| SP6 | ENST00000342234 | 3,2443 |
| MRPS27 | ENST00000457646 | 3,2407 |
| SCG3 | ENST00000542355 | 3,2393 |
| LINC00665 | ENST00000591372 | 3,2356 |
| HOPX | ENST00000553379 | 3,2348 |
| TMEM54 | ENST00000482771 | 3,2331 |
| CD55 | ENST00000343420 | 3,231 |
| NPEPPS | ENST00000678461 | 3,2223 |
| TPM1 | ENST00000559397 | 3,2161 |
| CDC25A | ENST00000437972 | 3,2125 |
| HDHD5 | ENST00000155674 | 3,2112 |
| CC2D2A | ENST00000652443 | 3,2068 |
| ZDHHC16 | ENST00000414567 | 3,2001 |
| EDC3 | ENST00000566219 | 3,1984 |
| GPR89A | ENST00000465185 | 3,196 |
| AZIN2 | ENST00000481886 | 3,1905 |
| ACSBG1 | ENST00000558130 | 3,1904 |
| GRIK2 | ENST00000413795 | 3,1901 |
| MRPL4 | ENST00000588502 | 3,1893 |
| FAM104A | ENST00000581110 | 3,1883 |
| SCAMP4 | ENST00000409472 | 3,1857 |
| METTL22 | ENST00000567295 | 3,1851 |
| BNC1 | ENST00000345382 | 3,1845 |
| FBLIM1 | ENST00000441801 | 3,1839 |
| IGF2BP2 | ENST00000461957 | 3,1732 |
| SETD4 | ENST00000469482 | 3,1673 |
| GPAA1P2 | ENST00000606848 | 3,1619 |
| BDNF | ENST00000525528 | 3,1618 |
| MUTYH | ENST00000456914 | 3,1611 |
| CEP104 | ENST00000675677 | 3,1604 |
| SIGIRR | ENST00000526395 | 3,1588 |
| PYCR1 | ENST00000405481 | 3,1579 |
| TBC1D3G | ENST00000569055 | 3,1518 |
| NCOR1 | ENST00000464381 | 3,1498 |
| MMP28 | ENST00000616383 | 3,1438 |
| CBS | ENST00000461686 | 3,1396 |
| SLC37A4 | ENST00000638360 | 3,1392 |
| OPRL1 | ENST00000355631 | 3,1369 |

| | | |
|---|---|---|
| PHKA2 | ENST00000481718 | -2,9233 |
| DCAF11 | ENST00000558914 | -2,926 |
| RGS9 | ENST00000582940 | -2,9271 |
| HHLA3 | ENST00000689607 | -2,9284 |
| ACR | ENST00000216139 | -2,9304 |
| ANK2 | ENST00000683893 | -2,938 |
| IFIH1 | ENST00000649554 | -2,9406 |
| PPT2-EGFL8 | ENST00000585246 | -2,9439 |
| SELENOT | ENST00000480740 | -2,9449 |
| TMEM8B | ENST00000650015 | -2,953 |
| CTC1 | ENST00000449476 | -2,954 |
| SLC35D2 | ENST00000490599 | -2,9544 |
| ZNF603P | ENST00000405613 | -2,9578 |
| SMIM2-AS1 | ENST00000659169 | -2,9614 |
| TIMM50 | ENST00000595286 | -2,9663 |
| ELMO2 | ENST00000396391 | -2,9693 |
| ABCB9 | ENST00000392439 | -2,9704 |
| NECAP1 | ENST00000640099 | -2,9736 |
| FADS1 | ENST00000536991 | -2,9762 |
| BTD | ENST00000383778 | -2,9922 |
| DDX58 | ENST00000679771 | -2,9932 |
| DMD | ENST00000681026 | -3,0019 |
| ERLEC1 | ENST00000692745 | -3,0032 |
| SPOP | ENST00000347630 | -3,0046 |
| EPM2A | ENST00000638554 | -3,007 |
| KCTD13 | ENST00000563955 | -3,0115 |
| ENKUR | ENST00000483339 | -3,0327 |
| ZFP69B | ENST00000469416 | -3,0417 |
| CCDC30 | ENST00000666893 | -3,0444 |
| MID1 | ENST00000687008 | -3,0467 |
| MACF1 | ENST00000524432 | -3,0478 |
| EIF2AK2 | ENST00000679507 | -3,0479 |
| PHLDB1 | ENST00000527898 | -3,0514 |
| EMC9 | ENST00000560600 | -3,0521 |
| EP400P1 | ENST00000389560 | -3,0664 |
| PIGX | ENST00000451319 | -3,0685 |
| SGCE | ENST00000472326 | -3,0697 |
| GLIDR | ENST00000667968 | -3,0755 |
| DMAC2L | ENST00000554438 | -3,0803 |
| THUMPD3-AS1 | ENST00000665171 | -3,0854 |
| CD109 | ENST00000437994 | -3,0868 |
| ZNF599 | ENST00000673678 | -3,089 |
| INTS11 | ENST00000533916 | -3,0899 |
| INTS11 | ENST00000467408 | -3,0909 |
| MIA2 | ENST00000348007 | -3,0922 |
| BOC | ENST00000488486 | -3,0923 |
| ETNK1 | ENST00000672951 | -3,0968 |
| NR2F1-AS1 | ENST00000671514 | -3,099 |
| HID1 | ENST00000532894 | -3,0993 |
| FHL1 | ENST00000420362 | -3,0994 |
| CFAP206 | ENST00000489338 | -3,1122 |
| NCKAP1L | ENST00000293373 | -3,1165 |
| ABHD16A | ENST00000492084 | -3,1256 |
| FAM86B3P | ENST00000691228 | -3,1282 |
| DNM1P41 | ENST00000623634 | -3,1305 |
| HIF3A | ENST00000600383 | -3,1328 |
| ACSL6 | ENST00000489047 | -3,1349 |
| CCT6B | ENST00000577307 | -3,1381 |
| NSUN5P1 | ENST00000687920 | -3,1399 |
| TCF4 | ENST00000629387 | -3,1461 |
| GLB1L | ENST00000424620 | -3,1462 |
| ABHD11 | ENST00000222800 | -3,152 |
| MAP1LC3B | ENST00000564638 | -3,1521 |
| TUBGCP3 | ENST00000649778 | -3,1521 |
| LRRC75B | ENST00000404045 | -3,1575 |

| | | |
|---|---|---|
| POSTN | ENST00000497145 | 3,1353 |
| ZNF138 | ENST00000359735 | 3,1301 |
| FBXO42 | ENST00000444116 | 3,1278 |
| DNMT1 | ENST00000676868 | 3,1241 |
| EIF4G1 | ENST00000676453 | 3,116 |
| IL1B | ENST00000263341 | 3,1126 |
| CAMK2D | ENST00000509907 | 3,1099 |
| NOX4 | ENST00000263317 | 3,1049 |
| LRRC32 | ENST00000407242 | 3,1047 |
| LCP1 | ENST00000674665 | 3,1019 |
| ULK4P2 | ENST00000569682 | 3,0994 |
| USP53 | ENST00000514305 | 3,09 |
| EDAR | ENST00000258443 | 3,088 |
| EBF3 | ENST00000355311 | 3,0851 |
| TRIM37 | ENST00000393065 | 3,0794 |
| ABCA13 | ENST00000484268 | 3,0764 |
| MPND | ENST00000601877 | 3,0749 |
| ALKBH6 | ENST00000590666 | 3,0692 |
| LINC01730 | ENST00000613868 | 3,0684 |
| ERMP1 | ENST00000688283 | 3,0639 |
| CSDE1 | ENST00000483407 | 3,0579 |
| RPP40 | ENST00000468105 | 3,0558 |
| DDHD2 | ENST00000528613 | 3,055 |
| AFDN | ENST00000366806 | 3,0526 |
| DMPK | ENST00000682898 | 3,0488 |
| WDR91 | ENST00000497853 | 3,0448 |
| SYNM | ENST00000560674 | 3,0408 |
| SLITRK3 | ENST00000241274 | 3,034 |
| ADGRA1 | ENST00000392606 | 3,0309 |
| NCAPH | ENST00000435349 | 3,0306 |
| TMEM132C | ENST00000435159 | 3,0299 |
| NGEF | ENST00000424488 | 3,0253 |
| SLITRK2 | ENST00000370490 | 3,0244 |
| C1orf112 | ENST00000359326 | 3,0241 |
| ARHGEF28 | ENST00000296799 | 3,024 |
| RYR3 | ENST00000415757 | 3,0202 |
| AJAP1 | ENST00000378191 | 3,0128 |
| LIMS2 | ENST00000426981 | 3,0119 |
| MESD | ENST00000560244 | 3,0085 |
| NOL10 | ENST00000695470 | 3,0084 |
| RAB3GAP2 | ENST00000685286 | 3,0075 |
| LRRC32 | ENST00000464145 | 3,0015 |
| LINC01561 | ENST00000623529 | 3 |
| RNASEH2B | ENST00000642721 | 2,9935 |
| CCDC57 | ENST00000578187 | 2,9894 |
| HEXIM2 | ENST00000591070 | 2,9875 |
| RNF114 | ENST00000623528 | 2,9829 |
| PABPC4 | ENST00000525751 | 2,9777 |
| TSPAN6 | ENST00000614008 | 2,9767 |
| SP140L | ENST00000396563 | 2,9761 |
| OCEL1 | ENST00000601529 | 2,9756 |
| LBR | ENST00000424022 | 2,9703 |
| TAF6 | ENST00000487115 | 2,9574 |
| APBB2 | ENST00000509446 | 2,9546 |
| UCHL5 | ENST00000367450 | 2,9542 |
| COL13A1 | ENST00000683993 | 2,9483 |
| TSFM | ENST00000457189 | 2,9457 |
| DCTN2 | ENST00000550576 | 2,9445 |
| RPA2 | ENST00000373909 | 2,9412 |
| C18orf21 | ENST00000269194 | 2,9318 |
| PAK5 | ENST00000378423 | 2,9307 |
| GLTP | ENST00000537066 | 2,9145 |
| PIGS | ENST00000577594 | 2,9141 |
| PRR5L | ENST00000389693 | 2,9137 |
| GDPD5 | ENST00000531759 | 2,9119 |

| | | |
|---|---|---|
| EHBP1-AS1 | ENST00000429952 | -3,1595 |
| PPID | ENST00000512699 | -3,177 |
| MAP3K3 | ENST00000584573 | -3,1774 |
| KIFC3 | ENST00000565270 | -3,1777 |
| FBXL2 | ENST00000451636 | -3,1816 |
| FAM71E1 | ENST00000600100 | -3,1824 |
| NT5M | ENST00000478373 | -3,1858 |
| DARS2 | ENST00000650297 | -3,189 |
| ATG13 | ENST00000528145 | -3,1896 |
| IMMP1L | ENST00000532624 | -3,1933 |
| WDR55 | ENST00000520764 | -3,1948 |
| H1-4 | ENST00000304218 | -3,2004 |
| IMP4 | ENST00000477375 | -3,2039 |
| SOX6 | ENST00000396356 | -3,2101 |
| LURAP1L | ENST00000489107 | -3,2113 |
| ZNF204P | ENST00000687798 | -3,218 |
| LSP1P4 | ENST00000692653 | -3,2223 |
| ADAM15 | ENST00000526491 | -3,2263 |
| MOCS1 | ENST00000373195 | -3,2263 |
| VDAC2 | ENST00000460044 | -3,2334 |
| PLEKHA5 | ENST00000510738 | -3,2372 |
| NDRG2 | ENST00000557676 | -3,2482 |
| CDH7 | ENST00000323011 | -3,2538 |
| NRL | ENST00000396997 | -3,2542 |
| ASXL1 | ENST00000644168 | -3,2579 |
| SH3GLB2 | ENST00000372554 | -3,266 |
| CYBC1 | ENST00000584791 | -3,2747 |
| TNFAIP3 | ENST00000420009 | -3,2763 |
| TMBIM6 | ENST00000550445 | -3,2838 |
| GALR1 | ENST00000299727 | -3,2889 |
| ERI2 | ENST00000564349 | -3,2901 |
| LINC01278 | ENST00000669201 | -3,2951 |
| ELAVL4 | ENST00000371827 | -3,2959 |
| CXCL2 | ENST00000296031 | -3,2986 |
| NARF | ENST00000579198 | -3,2991 |
| RPS24 | ENST00000475468 | -3,2991 |
| CA13 | ENST00000522631 | -3,2998 |
| COL9A2 | ENST00000488463 | -3,305 |
| ZNF668 | ENST00000426488 | -3,3051 |
| KAT7 | ENST00000512616 | -3,3072 |
| BTBD1 | ENST00000379403 | -3,3112 |
| MCPH1 | ENST00000687720 | -3,3154 |
| IZUMO4 | ENST00000395301 | -3,3223 |
| MYCBP2 | ENST00000695090 | -3,3306 |
| RBM23 | ENST00000553738 | -3,3308 |
| GGT7 | ENST00000470952 | -3,3324 |
| ASL | ENST00000380839 | -3,3411 |
| RPL31 | ENST00000441435 | -3,3522 |
| SLC41A3 | ENST00000504118 | -3,3557 |
| FBXO42 | ENST00000456164 | -3,377 |
| FRG1HP | ENST00000692434 | -3,3783 |
| GMDS-DT | ENST00000654526 | -3,3803 |
| SNHG17 | ENST00000654183 | -3,381 |
| ERBB2 | ENST00000406381 | -3,3855 |
| ITPRID1 | ENST00000615280 | -3,3911 |
| ACBD5 | ENST00000375905 | -3,3975 |
| FAM166B | ENST00000399742 | -3,4018 |
| POMZP3 | ENST00000454397 | -3,4104 |
| TERF1 | ENST00000678997 | -3,4114 |
| RNF4 | ENST00000502316 | -3,4186 |
| AASS | ENST00000460376 | -3,4213 |
| RRAGB | ENST00000414239 | -3,4325 |
| FGF22 | ENST00000586042 | -3,4403 |
| LENG8 | ENST00000462541 | -3,4406 |
| ABCA8 | ENST00000430352 | -3,4425 |

| | | |
|---|---|---|
| LINC01836 | ENST00000606242 | 2,9102 |
| ATP10A | ENST00000674138 | 2,9097 |
| KCNIP4 | ENST00000359001 | 2,9035 |
| LRRC17 | ENST00000498487 | 2,9013 |
| CBY1 | ENST00000492537 | 2,8999 |
| ETFDH | ENST00000684611 | 2,8983 |
| DLAT | ENST00000679614 | 2,8928 |
| NFYC | ENST00000425457 | 2,886 |
| TEDC2 | ENST00000483320 | 2,885 |
| ZNF280D | ENST00000559237 | 2,8844 |
| ICE2 | ENST00000560895 | 2,8843 |
| MVD | ENST00000568133 | 2,8823 |
| KMT2B | ENST00000685168 | 2,8815 |
| WWOX | ENST00000355860 | 2,8802 |
| LINC02610 | ENST00000409070 | 2,8798 |
| TAF1D | ENST00000527169 | 2,8791 |
| HMOX2 | ENST00000572812 | 2,8789 |
| SLC6A15 | ENST00000450363 | 2,8727 |
| NECTIN4 | ENST00000368012 | 2,8702 |
| KIAA1755 | ENST00000487506 | 2,866 |
| ZNF726 | ENST00000531821 | 2,8634 |
| MTFP1 | ENST00000407550 | 2,8613 |
| ANO7 | ENST00000481071 | 2,8545 |
| SZT2 | ENST00000562955 | 2,8545 |
| BICDL1 | ENST00000397558 | 2,8544 |
| PXYLP1 | ENST00000511968 | 2,8507 |
| WDR46 | ENST00000468157 | 2,8482 |
| FAM78B | ENST00000456900 | 2,848 |
| RBM19 | ENST00000553232 | 2,8437 |
| DNAAF3 | ENST00000526003 | 2,8352 |
| CD9 | ENST00000538418 | 2,8347 |
| FGGY | ENST00000371210 | 2,8316 |
| CREM | ENST00000337656 | 2,8236 |
| LINC00342 | ENST00000660641 | 2,8215 |
| HJURP | ENST00000432087 | 2,8183 |
| RUSC1 | ENST00000497930 | 2,8164 |
| DISP1 | ENST00000674736 | 2,816 |
| GMPR2 | ENST00000558007 | 2,8154 |
| ACSL6 | ENST00000652375 | 2,815 |
| TYK2 | ENST00000524462 | 2,8114 |
| TBC1D2 | ENST00000375064 | 2,8111 |
| HSP90B1 | ENST00000550595 | 2,8052 |
| SEPTIN7P2 | ENST00000690171 | 2,7956 |
| PDE4DIP | ENST00000695772 | 2,7953 |
| ZCCHC9 | ENST00000438268 | 2,7941 |
| IMPDH2 | ENST00000677480 | 2,7911 |
| TM7SF3 | ENST00000545303 | 2,7859 |
| HSD11B1L | ENST00000583928 | 2,7832 |
| TYSND1 | ENST00000479086 | 2,7815 |
| NRN1 | ENST00000244766 | 2,7771 |
| HGS | ENST00000678866 | 2,7765 |
| ELMOD3 | ENST00000440462 | 2,7696 |
| SLC37A3 | ENST00000473707 | 2,7644 |
| ZNF667-AS1 | ENST00000685306 | 2,7636 |
| TMEM143 | ENST00000600816 | 2,7572 |
| SLC16A3 | ENST00000584781 | 2,757 |
| CTSH | ENST00000676850 | 2,7503 |
| CHRNA5 | ENST00000394802 | 2,7443 |
| TMEM219 | ENST00000569445 | 2,7354 |
| SP2 | ENST00000637314 | 2,7266 |
| MRAS | ENST00000621127 | 2,7257 |
| ENTPD6 | ENST00000463734 | 2,7211 |
| FAM193B | ENST00000504130 | 2,7192 |
| TOMM40L | ENST00000367988 | 2,7177 |
| OXCT1-AS1 | ENST00000508458 | 2,7163 |

| | | |
|---|---|---|
| NSUN5P2 | ENST00000444583 | -3,4437 |
| ZNF559 | ENST00000605750 | -3,4484 |
| ZNF384 | ENST00000535485 | -3,4486 |
| ZC2HC1C | ENST00000238686 | -3,4488 |
| LSP1P4 | ENST00000689711 | -3,451 |
| ZNF335 | ENST00000475002 | -3,4511 |
| TAF11 | ENST00000689560 | -3,4524 |
| AAAS | ENST00000546562 | -3,4544 |
| RPS6KC1 | ENST00000543470 | -3,4582 |
| ZNF189 | ENST00000259395 | -3,4677 |
| IQCK | ENST00000308214 | -3,4723 |
| ISY1 | ENST00000393292 | -3,4782 |
| PRRC2B | ENST00000684694 | -3,479 |
| COPS7B | ENST00000412922 | -3,4821 |
| BCLAF1 | ENST00000527759 | -3,4831 |
| SOCS2 | ENST00000622746 | -3,4831 |
| RNF121 | ENST00000530058 | -3,4862 |
| POLR2J3 | ENST00000503564 | -3,4918 |
| CALB1 | ENST00000469032 | -3,492 |
| PRKAR2A-AS1 | ENST00000416209 | -3,4946 |
| CCT8 | ENST00000470450 | -3,4977 |
| CEPT1 | ENST00000498239 | -3,499 |
| ATP7B | ENST00000448424 | -3,5 |
| ATF7 | ENST00000591397 | -3,502 |
| SPTBN4 | ENST00000598249 | -3,5034 |
| NUDT2 | ENST00000618590 | -3,5042 |
| SNHG14 | ENST00000658853 | -3,5105 |
| HSD17B10 | ENST00000684251 | -3,5192 |
| GALNTL6 | ENST00000506823 | -3,5365 |
| ARHGAP21 | ENST00000482792 | -3,541 |
| DAPK1 | ENST00000469067 | -3,5428 |
| STK39 | ENST00000487143 | -3,5447 |
| UBR7 | ENST00000554232 | -3,5544 |
| ACAD11 | ENST00000481970 | -3,5604 |
| HDLBP | ENST00000460826 | -3,5688 |
| GLRA2 | ENST00000218075 | -3,5787 |
| GPBP1L1 | ENST00000480083 | -3,5876 |
| WWOX | ENST00000406884 | -3,5931 |
| LINC01128 | ENST00000659124 | -3,5972 |
| CTSA | ENST00000606066 | -3,6081 |
| SEZ6 | ENST00000544224 | -3,6152 |
| AFAP1L1 | ENST00000296721 | -3,6153 |
| NAPA | ENST00000597271 | -3,6196 |
| CENPV | ENST00000472570 | -3,6221 |
| FANCC | ENST00000375305 | -3,6298 |
| ITGA6 | ENST00000475302 | -3,6384 |
| THAP5 | ENST00000493722 | -3,6385 |
| METTL27 | ENST00000297873 | -3,6413 |
| SMYD2 | ENST00000484459 | -3,6496 |
| NPIPB5 | ENST00000424340 | -3,6499 |
| MTO1 | ENST00000522205 | -3,6643 |
| ANK2 | ENST00000682211 | -3,6797 |
| RUNDC3A | ENST00000590834 | -3,6912 |
| SETD9 | ENST00000628593 | -3,6924 |
| KEAP1 | ENST00000592478 | -3,6994 |
| CCDC144NL-AS1 | ENST00000577537 | -3,7034 |
| HLA-F | ENST00000484704 | -3,7076 |
| GALT | ENST00000473506 | -3,7238 |
| ARMH1 | ENST00000418779 | -3,7307 |
| BRD9 | ENST00000493082 | -3,7415 |
| EMSY-DT | ENST00000572035 | -3,7638 |
| TSPEAR-AS2 | ENST00000354333 | -3,7664 |
| ZFX | ENST00000379177 | -3,7673 |
| TBC1D23 | ENST00000471273 | -3,7783 |
| ENTPD1-AS1 | ENST00000671110 | -3,7885 |

| | | |
|---|---|---|
| CMBL | ENST00000506821 | 2,7158 |
| MUC3A | ENST00000414964 | 2,7148 |
| NUDT9 | ENST00000440591 | 2,7068 |
| ARHGEF35 | ENST00000688754 | 2,6981 |
| RGS9 | ENST00000637818 | 2,6973 |
| MIR100HG | ENST00000690198 | 2,6963 |
| SPTBN4 | ENST00000392023 | 2,6929 |
| RAB3IP | ENST00000550647 | 2,688 |
| RYK | ENST00000480381 | 2,6829 |
| PROSER3 | ENST00000396908 | 2,6825 |
| SGK3 | ENST00000345714 | 2,6798 |
| EPS15L1 | ENST00000535753 | 2,6656 |
| MAP3K2-DT | ENST00000692842 | 2,6636 |
| RECQL5 | ENST00000580707 | 2,6611 |
| ZNF696 | ENST00000521537 | 2,6555 |
| HHIP | ENST00000434550 | 2,6495 |
| LINC00662 | ENST00000655567 | 2,645 |
| LRRC28 | ENST00000561276 | 2,6449 |
| TRMT2A | ENST00000464535 | 2,6411 |
| KCNMA1 | ENST00000639716 | 2,629 |
| FN1 | ENST00000480737 | 2,6225 |
| CACNB3 | ENST00000550391 | 2,6196 |
| ZNF532 | ENST00000592452 | 2,6122 |
| SPRTN | ENST00000008440 | 2,612 |
| ANXA2 | ENST00000559780 | 2,6034 |
| TSEN15 | ENST00000644815 | 2,6015 |
| MED15 | ENST00000406969 | 2,5904 |
| ENGASE | ENST00000578419 | 2,5902 |
| PGM2 | ENST00000505986 | 2,5823 |
| MBD2 | ENST00000583046 | 2,5804 |
| BAIAP2 | ENST00000572073 | 2,5794 |
| TM2D3 | ENST00000560910 | 2,572 |
| PDXDC2P-NPI | ENST00000525331 | 2,5699 |
| ZNF667-AS1 | ENST00000693493 | 2,5686 |
| SLITRK4 | ENST00000596188 | 2,5673 |
| UNC5B-AS1 | ENST00000447119 | 2,5673 |
| RANBP1 | ENST00000411892 | 2,5632 |
| EXOSC7 | ENST00000468667 | 2,5609 |
| RNF10 | ENST00000543675 | 2,5599 |
| ASPSCR1 | ENST00000578236 | 2,5593 |
| WDR5B-DT | ENST00000667395 | 2,5567 |
| ZNF419 | ENST00000426954 | 2,5514 |
| TRIP11 | ENST00000555105 | 2,5469 |
| CYREN | ENST00000617987 | 2,5461 |
| OR2A9P | ENST00000461471 | 2,5434 |
| CERS5 | ENST00000547800 | 2,5309 |
| ZNF667-AS1 | ENST00000654200 | 2,5302 |
| SOCS2 | ENST00000549887 | 2,5286 |
| GABRG3 | ENST00000333743 | 2,5236 |
| AMT | ENST00000465925 | 2,5163 |
| SETDB1 | ENST00000525956 | 2,5134 |
| SRA1 | ENST00000602875 | 2,5112 |
| RARA | ENST00000475125 | 2,5104 |
| TPRKB | ENST00000497464 | 2,5075 |
| ARRDC2 | ENST00000379656 | 2,5049 |
| ZC3H13 | ENST00000428921 | 2,5039 |
| ZC2HC1A | ENST00000519307 | 2,5035 |
| ZSCAN9 | ENST00000425468 | 2,5033 |
| THUMPD3 | ENST00000441127 | 2,5027 |
| RGS4 | ENST00000531057 | 2,5025 |
| TPM4 | ENST00000647037 | 2,5013 |
| HOXD9 | ENST00000249499 | 2,4999 |
| PRCD | ENST00000587289 | 2,4959 |
| ALG11 | ENST00000679544 | 2,4923 |
| TPGS2 | ENST00000590500 | 2,4883 |

| | | |
|---|---|---|
| CMPK2 | ENST00000256722 | -3,7896 |
| SEC16A | ENST00000684901 | -3,7913 |
| MAPK10 | ENST00000641493 | -3,7958 |
| ALDH3A2 | ENST00000573505 | -3,8018 |
| ALS2 | ENST00000680441 | -3,8149 |
| CARMIL2 | ENST00000334583 | -3,8149 |
| GNG8 | ENST00000693335 | -3,8167 |
| TMEM234 | ENST00000344461 | -3,8203 |
| MPV17 | ENST00000233545 | -3,8226 |
| IFT43 | ENST00000554026 | -3,826 |
| GARS1 | ENST00000676403 | -3,8375 |
| MADD | ENST00000395336 | -3,8402 |
| LYRM1 | ENST00000568820 | -3,8453 |
| SUN2 | ENST00000405018 | -3,8457 |
| GYG1 | ENST00000627418 | -3,8483 |
| CPAMD8 | ENST00000597335 | -3,8503 |
| PLAGL1 | ENST00000649211 | -3,8508 |
| MARK3 | ENST00000560417 | -3,8592 |
| AFF2 | ENST00000370460 | -3,8598 |
| HOGA1 | ENST00000370647 | -3,8605 |
| METTL8 | ENST00000612742 | -3,8607 |
| TDRKH | ENST00000368824 | -3,8635 |
| SCFD1 | ENST00000553278 | -3,8831 |
| MYO5A | ENST00000687172 | -3,8877 |
| REXO5 | ENST00000348433 | -3,8883 |
| MTR | ENST00000679569 | -3,889 |
| PACSIN1 | ENST00000538621 | -3,9022 |
| MTCH1 | ENST00000695045 | -3,9143 |
| MED12 | ENST00000690250 | -3,9158 |
| GGA1 | ENST00000411501 | -3,9179 |
| SPIN3 | ENST00000639007 | -3,9212 |
| TTLL3 | ENST00000493241 | -3,9228 |
| GGCX | ENST00000691348 | -3,9254 |
| BANCR | ENST00000690153 | -3,9418 |
| AURKA | ENST00000422322 | -3,9461 |
| SGK3 | ENST00000521198 | -3,9465 |
| CKMT1B | ENST00000441322 | -3,949 |
| BSCL2 | ENST00000684609 | -3,9507 |
| MRS2 | ENST00000274747 | -3,9642 |
| PRKD2 | ENST00000597390 | -3,9667 |
| RIDA | ENST00000522791 | -3,9759 |
| ADNP | ENST00000621696 | -3,9791 |
| PAK6 | ENST00000560346 | -3,9793 |
| LINC01088 | ENST00000513153 | -3,9831 |
| ALDH16A1 | ENST00000593417 | -3,9835 |
| NRXN1 | ENST00000625672 | -3,9896 |
| FAM86B2 | ENST00000685726 | -3,9916 |
| PRR22 | ENST00000419421 | -3,9986 |
| RBM23 | ENST00000346528 | -3,9989 |
| DMXL1 | ENST00000514595 | -4,0072 |
| BRD9 | ENST00000490814 | -4,0095 |
| COMMD5 | ENST00000543949 | -4,0124 |
| SIPA1L2 | ENST00000674749 | -4,0142 |
| DNAI1 | ENST00000442556 | -4,0395 |
| DHCR7 | ENST00000527316 | -4,0402 |
| UBE2A | ENST00000630695 | -4,0448 |
| CDK7 | ENST00000629350 | -4,0457 |
| KIFC3 | ENST00000565753 | -4,0548 |
| DDX5 | ENST00000577922 | -4,0549 |
| PPP1R12B | ENST00000391959 | -4,0564 |
| CD302 | ENST00000429078 | -4,0587 |
| JMJD6 | ENST00000591460 | -4,0628 |
| POLG2 | ENST00000578997 | -4,0746 |
| CABP1 | ENST00000316803 | -4,0776 |
| STK36 | ENST00000462031 | -4,0813 |

| | | |
|---|---|---|
| MFSD1 | ENST00000651862 | 2,4869 |
| ZNF93 | ENST00000592160 | 2,4798 |
| CAPZB | ENST00000433834 | 2,4763 |
| KLC1 | ENST00000557686 | 2,4741 |
| GTF2H2B | ENST00000508065 | 2,4727 |
| PDLIM1 | ENST00000490391 | 2,4678 |
| MRPS25 | ENST00000695395 | 2,4659 |
| MTRR | ENST00000506115 | 2,4644 |
| PGLS-DT | ENST00000596643 | 2,4643 |
| ZBTB17 | ENST00000479282 | 2,4573 |
| PHETA1 | ENST00000361483 | 2,4566 |
| DZANK1 | ENST00000358866 | 2,448 |
| GNA12 | ENST00000471281 | 2,4478 |
| B3GNTL1 | ENST00000571954 | 2,4471 |
| H19 | ENST00000412788 | 2,4458 |
| IL32 | ENST00000525228 | 2,4444 |
| MGST3 | ENST00000461308 | 2,443 |
| ECPAS | ENST00000374383 | 2,4349 |
| LAMB1 | ENST00000679244 | 2,434 |
| PTPN6 | ENST00000399448 | 2,428 |
| MIR4435-2HG | ENST00000666144 | 2,4262 |
| TLE1P1 | ENST00000422965 | 2,4237 |
| DNAH14 | ENST00000486657 | 2,4222 |
| ARID4B | ENST00000494543 | 2,4166 |
| ZNF84 | ENST00000438628 | 2,4135 |
| CIRBP | ENST00000589235 | 2,4131 |
| ZNF414 | ENST00000594748 | 2,404 |
| SLC16A1-AS1 | ENST00000416193 | 2,3961 |
| NAMPT | ENST00000681631 | 2,3923 |
| DTWD1 | ENST00000557968 | 2,3922 |
| IFT122 | ENST00000692321 | 2,3916 |
| QPCTL | ENST00000366382 | 2,3886 |
| PISD | ENST00000474017 | 2,3864 |
| MELK | ENST00000541717 | 2,3835 |
| CLDN6 | ENST00000328796 | 2,3832 |
| DAP3 | ENST00000471214 | 2,3832 |
| GALNT1 | ENST00000589189 | 2,3767 |
| CTDP1 | ENST00000299543 | 2,3712 |
| GAS5 | ENST00000450589 | 2,364 |
| SP100 | ENST00000494508 | 2,3637 |
| AAMP | ENST00000489767 | 2,3601 |
| HNRNPL | ENST00000595804 | 2,3544 |
| CAST | ENST00000514845 | 2,3537 |
| CEP63 | ENST00000684170 | 2,3535 |
| PNPLA2 | ENST00000526083 | 2,3505 |
| MAP4K2 | ENST00000489952 | 2,3504 |
| NPHP1 | ENST00000675752 | 2,3487 |
| BACE2 | ENST00000463674 | 2,3482 |
| LAMC3 | ENST00000462567 | 2,3467 |
| DYNC1I2 | ENST00000410079 | 2,3448 |
| MKNK2 | ENST00000586828 | 2,3397 |
| RN7SL731P | ENST00000584329 | 2,3356 |
| MTCL1 | ENST00000517570 | 2,3332 |
| MIR9-1HG | ENST00000471156 | 2,3328 |
| POLE4 | ENST00000485527 | 2,3301 |
| ADCY8 | ENST00000377928 | 2,33 |
| FXN | ENST00000396366 | 2,3291 |
| PLSCR3 | ENST00000576201 | 2,3276 |
| APTX | ENST00000673487 | 2,3233 |
| ZFAT | ENST00000518408 | 2,3179 |
| LYPD6B | ENST00000280115 | 2,3164 |
| KRT80 | ENST00000466011 | 2,3143 |
| H19 | ENST00000414790 | 2,3132 |
| ING4 | ENST00000486287 | 2,3131 |
| ADSL | ENST00000679904 | 2,3063 |

| | | |
|---|---|---|
| RBCK1 | ENST00000382214 | -4,0908 |
| APOL4 | ENST00000332987 | -4,0932 |
| SNX1 | ENST00000559401 | -4,0968 |
| ZNF181 | ENST00000448715 | -4,1039 |
| WAC | ENST00000651598 | -4,1076 |
| VPS28 | ENST00000531924 | -4,1132 |
| TPCN1 | ENST00000547955 | -4,119 |
| FLII | ENST00000478416 | -4,1209 |
| SNX21 | ENST00000372541 | -4,123 |
| ERCC2 | ENST00000391941 | -4,1284 |
| SRGAP2 | ENST00000604925 | -4,1374 |
| FAM149A | ENST00000514829 | -4,1412 |
| SAFB | ENST00000589863 | -4,1413 |
| TRPV3 | ENST00000381913 | -4,1473 |
| FAM216B | ENST00000313851 | -4,1493 |
| NPTX1 | ENST00000535681 | -4,1497 |
| RGS9 | ENST00000577595 | -4,1502 |
| WT1 | ENST00000379079 | -4,1544 |
| NCOA3 | ENST00000371997 | -4,1581 |
| DNAJB1 | ENST00000601533 | -4,1591 |
| AMDHD2 | ENST00000569219 | -4,1605 |
| KMT2D | ENST00000683543 | -4,1647 |
| SERTAD3 | ENST00000392028 | -4,1733 |
| ZNF195 | ENST00000399602 | -4,1742 |
| NDRG2 | ENST00000554104 | -4,1815 |
| SETD6 | ENST00000427443 | -4,1928 |
| EXT2 | ENST00000684039 | -4,1934 |
| DKKL1 | ENST00000221498 | -4,1984 |
| PORCN | ENST00000682661 | -4,2009 |
| HSD11B1L | ENST00000581423 | -4,2011 |
| ALG3 | ENST00000462735 | -4,2113 |
| PLA2G6 | ENST00000664587 | -4,2227 |
| RMND1 | ENST00000684715 | -4,2247 |
| EGR4 | ENST00000436467 | -4,227 |
| STX1A | ENST00000462135 | -4,2282 |
| NPIPB4 | ENST00000682606 | -4,2369 |
| ALS2 | ENST00000681303 | -4,2407 |
| AKAP8L | ENST00000680103 | -4,2428 |
| MOV10 | ENST00000468624 | -4,2477 |
| ENTPD3-AS1 | ENST00000425156 | -4,2525 |
| TRAP1 | ENST00000574175 | -4,2577 |
| SNHG15 | ENST00000577700 | -4,2578 |
| EXOSC8 | ENST00000685643 | -4,2592 |
| CD99P1 | ENST00000685470 | -4,2598 |
| CHRNA1 | ENST00000409323 | -4,2658 |
| COX18 | ENST00000295890 | -4,272 |
| GCC2 | ENST00000688612 | -4,272 |
| POR | ENST00000447222 | -4,2732 |
| CHRFAM7A | ENST00000299847 | -4,2735 |
| KIF1A | ENST00000648680 | -4,2815 |
| B3GNTL1 | ENST00000572977 | -4,2871 |
| SUSD2 | ENST00000463101 | -4,2888 |
| SCML1 | ENST00000380041 | -4,2939 |
| DYNLT4 | ENST00000675259 | -4,2941 |
| HSPA9 | ENST00000508003 | -4,3042 |
| TNNT2 | ENST00000476888 | -4,3087 |
| SEL1L3 | ENST00000512286 | -4,3108 |
| LINC00632 | ENST00000648347 | -4,3273 |
| GATD3 | ENST00000348499 | -4,3295 |
| CNOT8 | ENST00000520671 | -4,3296 |
| ZNF333 | ENST00000540689 | -4,3306 |
| COPS7B | ENST00000373608 | -4,3318 |
| CASC15 | ENST00000666379 | -4,332 |
| LINC00476 | ENST00000691767 | -4,3343 |
| LHX2 | ENST00000446480 | -4,3372 |

| | | |
|---|---|---|
| SERAC1 | ENST00000642903 | 2,3047 |
| COQ10A | ENST00000551566 | 2,2989 |
| KIT | ENST00000688060 | 2,2984 |
| RIPOR1 | ENST00000379312 | 2,2981 |
| TBCB | ENST00000589996 | 2,2948 |
| LARP4B | ENST00000690211 | 2,2938 |
| MPP4 | ENST00000396886 | 2,2885 |
| BYSL | ENST00000475702 | 2,2831 |
| CKMT1B | ENST00000437534 | 2,2756 |
| MASTL | ENST00000342386 | 2,2725 |
| LMNA | ENST00000683032 | 2,2694 |
| TM4SF1 | ENST00000493348 | 2,2635 |
| CCDC125 | ENST00000511257 | 2,2599 |
| GPRC5A | ENST00000014914 | 2,2586 |
| FTSJ1 | ENST00000019019 | 2,2516 |
| ELF3 | ENST00000475698 | 2,2482 |
| RPAIN | ENST00000574003 | 2,2467 |
| SF1 | ENST00000448404 | 2,2438 |
| SCUBE2 | ENST00000450649 | 2,2431 |
| MRPS18C | ENST00000505719 | 2,2414 |
| CEP72 | ENST00000512038 | 2,2368 |
| SLC25A36 | ENST00000514629 | 2,2366 |
| ELAPOR2 | ENST00000394714 | 2,2356 |
| PDE4B | ENST00000480109 | 2,2355 |
| ASAH2 | ENST00000682911 | 2,2333 |
| PSME3IP1 | ENST00000389447 | 2,2331 |
| MPC1 | ENST00000366868 | 2,2214 |
| SCAMP2 | ENST00000567638 | 2,2213 |
| MEOX1 | ENST00000318579 | 2,2211 |
| VLDLR-AS1 | ENST00000648733 | 2,2174 |
| FGFR1 | ENST00000464163 | 2,2163 |
| KRIT1 | ENST00000430102 | 2,2127 |
| CAD | ENST00000428460 | 2,2098 |
| MMP14 | ENST00000548162 | 2,2092 |
| MMP19 | ENST00000548629 | 2,2069 |
| DPF1 | ENST00000456296 | 2,2043 |
| EIF3L | ENST00000406934 | 2,2022 |
| FXYD6 | ENST00000539526 | 2,1993 |
| FLJ12825 | ENST00000515617 | 2,1991 |
| RARS2 | ENST00000688391 | 2,1921 |
| RAB3GAP2 | ENST00000491305 | 2,188 |
| PHF19 | ENST00000462229 | 2,1804 |
| ARSJ | ENST00000503013 | 2,1797 |
| BHLHE22 | ENST00000321870 | 2,1761 |
| PCDHGA6 | ENST00000517434 | 2,1716 |
| CBS | ENST00000398165 | 2,17 |
| PLGLB2 | ENST00000359481 | 2,164 |
| CENPN | ENST00000569461 | 2,1578 |
| OPCML | ENST00000374778 | 2,1552 |
| IFNGR1 | ENST00000458076 | 2,1521 |
| PPP2R3C | ENST00000553273 | 2,1516 |
| SCN2A | ENST00000283256 | 2,1471 |
| ABCA13 | ENST00000435803 | 2,1453 |
| ANXA4 | ENST00000460439 | 2,1385 |
| ARSL | ENST00000683071 | 2,135 |
| FAM66D | ENST00000653408 | 2,1344 |
| ACAD9 | ENST00000681319 | 2,1334 |
| ATRX | ENST00000623316 | 2,1297 |
| WRNIP1 | ENST00000380771 | 2,1279 |
| MXRA5Y | ENST00000420610 | 2,1267 |
| ERBIN | ENST00000503913 | 2,1234 |
| NRG3 | ENST00000545131 | 2,1225 |
| AHI1 | ENST00000533029 | 2,1215 |
| HGS | ENST00000676729 | 2,1211 |
| TCIRG1 | ENST00000265686 | 2,1188 |

| | | |
|---|---|---|
| TAMALIN | ENST00000552049 | -4,3378 |
| ASTN1 | ENST00000424564 | -4,3458 |
| CACNA1C | ENST00000399634 | -4,3484 |
| HSD17B7 | ENST00000485405 | -4,3548 |
| PMCH | ENST00000329406 | -4,355 |
| MARK1 | ENST00000611084 | -4,3692 |
| NEDD4 | ENST00000648451 | -4,37 |
| HAUS4 | ENST00000553420 | -4,3753 |
| BAIAP3 | ENST00000628027 | -4,3785 |
| SNORD3A | ENST00000584923 | -4,387 |
| SIM2 | ENST00000290399 | -4,3929 |
| PLCG1 | ENST00000599785 | -4,4005 |
| NNT | ENST00000660676 | -4,4015 |
| NFE2L1-DT | ENST00000689902 | -4,4051 |
| GOLGA3 | ENST00000692831 | -4,4083 |
| TMEM220 | ENST00000455996 | -4,4086 |
| COMMD4 | ENST00000568301 | -4,4106 |
| GOLGA8T | ENST00000569052 | -4,4139 |
| MIR124-1HG | ENST00000667273 | -4,4175 |
| PARP6 | ENST00000544520 | -4,4205 |
| MAPK9 | ENST00000523583 | -4,4261 |
| COL4A6 | ENST00000621266 | -4,4285 |
| PIK3C3 | ENST00000398870 | -4,4297 |
| PRANCR | ENST00000656495 | -4,433 |
| POLR3E | ENST00000569787 | -4,436 |
| RYR1 | ENST00000689936 | -4,4367 |
| IL16 | ENST00000394660 | -4,4403 |
| RPS6KA2 | ENST00000510118 | -4,4438 |
| CNTRL | ENST00000691066 | -4,4459 |
| GOLGB1 | ENST00000694958 | -4,4489 |
| CBWD1 | ENST00000613988 | -4,456 |
| PPP5D1P | ENST00000675805 | -4,4624 |
| TMEM134 | ENST00000536773 | -4,4638 |
| TFAP2A | ENST00000379608 | -4,4673 |
| MINCR | ENST00000518073 | -4,4768 |
| CUL1 | ENST00000662670 | -4,5008 |
| MAN2B1 | ENST00000465830 | -4,5056 |
| RNFT1 | ENST00000477207 | -4,5079 |
| PPP1R32 | ENST00000432063 | -4,5136 |
| DDR1 | ENST00000454612 | -4,516 |
| EIF4G1 | ENST00000427845 | -4,5197 |
| CCND3 | ENST00000512426 | -4,5225 |
| TNNI2 | ENST00000252898 | -4,524 |
| PGAP2 | ENST00000477358 | -4,5248 |
| PIGT | ENST00000640253 | -4,5325 |
| TJAP1 | ENST00000372444 | -4,5481 |
| ASB16-AS1 | ENST00000588785 | -4,5571 |
| ATP6V0B | ENST00000498664 | -4,5571 |
| ANO6 | ENST00000680498 | -4,5577 |
| FAR2P4 | ENST00000416266 | -4,5632 |
| RPL34 | ENST00000502534 | -4,5632 |
| SDR42E2 | ENST00000687571 | -4,5645 |
| ANAPC11 | ENST00000577425 | -4,5677 |
| CYFIP2 | ENST00000616178 | -4,5699 |
| ELOVL1 | ENST00000621943 | -4,5723 |
| LGALS9 | ENST00000395473 | -4,578 |
| PRR14 | ENST00000287463 | -4,5791 |
| KIF1A | ENST00000650430 | -4,5829 |
| TMC5 | ENST00000541464 | -4,5832 |
| CERS5 | ENST00000547787 | -4,5888 |
| PDE9A | ENST00000398232 | -4,589 |
| STAG3L4 | ENST00000689978 | -4,595 |
| BCAT2 | ENST00000402551 | -4,5953 |
| OSBPL6 | ENST00000315022 | -4,6 |
| AKAP13 | ENST00000557852 | -4,6007 |

| | | |
|---|---|---|
| ELF3-AS1 | ENST00000419190 | 2,1181 |
| PKMYT1 | ENST00000575632 | 2,1091 |
| PDHB | ENST00000461692 | 2,1069 |
| ITGA2B | ENST00000648408 | 2,1065 |
| ATXN2 | ENST00000647305 | 2,1059 |
| CDK11A | ENST00000468397 | 2,1044 |
| ST3GAL2 | ENST00000567586 | 2,1006 |
| BMP1 | ENST00000521385 | 2,1 |
| PRC1 | ENST00000560914 | 2,0966 |
| THAP7 | ENST00000476667 | 2,0963 |
| PDLIM3 | ENST00000620787 | 2,0934 |
| SGIP1 | ENST00000682476 | 2,0928 |
| CDIPT | ENST00000561555 | 2,0915 |
| DMAC2 | ENST00000301183 | 2,091 |
| TTPAL | ENST00000372904 | 2,0885 |
| CRELD1 | ENST00000602411 | 2,0826 |
| ATG4B | ENST00000344376 | 2,0816 |
| METTL17 | ENST00000554588 | 2,081 |
| IQCK | ENST00000564186 | 2,0782 |
| LARS1 | ENST00000674398 | 2,0762 |
| EIF4E2 | ENST00000687222 | 2,0745 |
| SSH3 | ENST00000529224 | 2,073 |
| OPTN | ENST00000378747 | 2,0727 |
| AKAP9 | ENST00000679521 | 2,0725 |
| PSMF1 | ENST00000479715 | 2,0721 |
| PPHLN1 | ENST00000256678 | 2,0642 |
| NPL | ENST00000614468 | 2,0629 |
| SOCS2 | ENST00000547229 | 2,0622 |
| TAF6 | ENST00000688197 | 2,0595 |
| BABAM1 | ENST00000595632 | 2,0409 |
| AUP1 | ENST00000462297 | 2,0381 |
| NDUFAF1 | ENST00000679240 | 2,038 |
| COL1A2 | ENST00000473573 | 2,0299 |
| LRRC43 | ENST00000541498 | 2,0297 |
| CDCA4 | ENST00000392590 | 2,0282 |
| EML2 | ENST00000588889 | 2,0268 |
| KIF5A | ENST00000676081 | 2,0267 |
| C16orf74 | ENST00000602675 | 2,0265 |
| UBE3B | ENST00000538070 | 2,0254 |
| ATL1 | ENST00000441560 | 2,0055 |
| DDR1 | ENST00000513749 | 2,003 |
| PTK2 | ENST00000523805 | 1,9989 |
| PABPC1 | ENST00000522387 | 1,9988 |
| MTHFD1 | ENST00000555709 | 1,9984 |
| PRXL2B | ENST00000493183 | 1,9964 |
| RBBP7 | ENST00000468092 | 1,9958 |
| LRRC34 | ENST00000522596 | 1,9957 |
| NEK8 | ENST00000593261 | 1,9916 |
| AIF1L | ENST00000372301 | 1,9897 |
| MFSD8 | ENST00000641447 | 1,9897 |
| ABCD3 | ENST00000493416 | 1,9884 |
| LSM14B | ENST00000400318 | 1,9877 |
| SGIP1 | ENST00000684369 | 1,987 |
| MSC-AS1 | ENST00000457356 | 1,9856 |
| PCDH7 | ENST00000509759 | 1,9841 |
| C17orf75 | ENST00000582961 | 1,9813 |
| CLCC1 | ENST00000675650 | 1,9808 |
| GRN | ENST00000587958 | 1,9759 |
| MYH9 | ENST00000477189 | 1,974 |
| UBAP2L | ENST00000433615 | 1,9735 |
| CRYZL1 | ENST00000381540 | 1,9728 |
| CHCHD7 | ENST00000523975 | 1,9669 |
| TMEM98 | ENST00000439138 | 1,9658 |
| ZCCHC8 | ENST00000540586 | 1,9629 |
| SOCS2-AS1 | ENST00000500986 | 1,9615 |

| | | |
|---|---|---|
| NUTM2B-AS1 | ENST00000665716 | -4,6007 |
| RAB3GAP2 | ENST00000693454 | -4,6081 |
| SLFN12 | ENST00000394562 | -4,6128 |
| RIMKLB | ENST00000510357 | -4,6258 |
| CDK14 | ENST00000406263 | -4,6273 |
| HARS2 | ENST00000647484 | -4,6289 |
| ZNF687 | ENST00000436614 | -4,6318 |
| RASSF8 | ENST00000615708 | -4,6343 |
| KDM5C | ENST00000349663 | -4,6388 |
| HECTD1 | ENST00000692014 | -4,6483 |
| POFUT2 | ENST00000334538 | -4,6554 |
| DPY19L2P2 | ENST00000411491 | -4,6625 |
| GNAS | ENST00000349036 | -4,6639 |
| ARMC10 | ENST00000425331 | -4,6665 |
| GPR61 | ENST00000527748 | -4,6733 |
| MSL3 | ENST00000398527 | -4,6751 |
| ERH | ENST00000216520 | -4,6754 |
| NEIL1 | ENST00000564784 | -4,6776 |
| PCDHA10 | ENST00000506939 | -4,685 |
| BRPF1 | ENST00000672126 | -4,6876 |
| ARFGEF2 | ENST00000679747 | -4,6952 |
| DECR1 | ENST00000519328 | -4,7026 |
| C21orf58 | ENST00000491666 | -4,7081 |
| MLIP | ENST00000514433 | -4,7124 |
| IFT88 | ENST00000482172 | -4,7184 |
| HCG18 | ENST00000664861 | -4,7289 |
| STAT5A | ENST00000590949 | -4,7318 |
| KIAA1109 | ENST00000264501 | -4,7403 |
| SETDB1 | ENST00000690229 | -4,7413 |
| MSLN | ENST00000563941 | -4,746 |
| LINC01094 | ENST00000671622 | -4,7524 |
| SH3YL1 | ENST00000605370 | -4,7562 |
| FPGS | ENST00000460181 | -4,7572 |
| REXO4 | ENST00000371935 | -4,763 |
| SNHG1 | ENST00000660547 | -4,7632 |
| QARS1 | ENST00000634473 | -4,7735 |
| ZBTB11 | ENST00000312938 | -4,781 |
| SNX11 | ENST00000580875 | -4,7828 |
| BCAP31 | ENST00000672675 | -4,7874 |
| RPS6KL1 | ENST00000555834 | -4,7938 |
| PRMT7 | ENST00000568975 | -4,8071 |
| RYR1 | ENST00000594335 | -4,8195 |
| TPR | ENST00000613151 | -4,8258 |
| IFT172 | ENST00000676119 | -4,8283 |
| LAS1L | ENST00000678074 | -4,8285 |
| APTR | ENST00000398043 | -4,8301 |
| FOS | ENST00000554212 | -4,8344 |
| ETNPPL | ENST00000296486 | -4,8406 |
| TMEM116 | ENST00000549537 | -4,8408 |
| WDR49 | ENST00000479765 | -4,8438 |
| ZSCAN25 | ENST00000394150 | -4,8438 |
| SMARCA4 | ENST00000645648 | -4,8442 |
| LARS1 | ENST00000674277 | -4,8459 |
| ABTB1 | ENST00000474129 | -4,8523 |
| ST18 | ENST00000276480 | -4,8523 |
| ZNF561-AS1 | ENST00000692903 | -4,8534 |
| BRPF3 | ENST00000449261 | -4,8615 |
| ACBD4 | ENST00000592162 | -4,8627 |
| TSNAX | ENST00000413309 | -4,8732 |
| DDX10 | ENST00000689812 | -4,8756 |
| VIPAS39 | ENST00000553888 | -4,8938 |
| IDH1 | ENST00000451391 | -4,9138 |
| CSAG3 | ENST00000638835 | -4,9155 |
| RNA5-8SN2 | ENST00000612463 | -4,9188 |
| CCDC120 | ENST00000603906 | -4,92 |

| | | |
|---|---|---|
| ATG13 | ENST00000527907 | 1,9598 |
| SVIL-AS1 | ENST00000438202 | 1,9589 |
| SHOC2 | ENST00000689997 | 1,9552 |
| DMPK | ENST00000354227 | 1,9508 |
| PRKAG2 | ENST00000652572 | 1,9504 |
| HSPD1 | ENST00000476746 | 1,9481 |
| SNHG12 | ENST00000531126 | 1,9474 |
| NELFCD | ENST00000486263 | 1,9455 |
| STAG3L4 | ENST00000689067 | 1,943 |
| ANAPC11 | ENST00000583839 | 1,9371 |
| ZSCAN2 | ENST00000540936 | 1,9309 |
| PLCH2 | ENST00000278878 | 1,9271 |
| SIGMAR1 | ENST00000477726 | 1,9269 |
| SLC26A6 | ENST00000469693 | 1,9269 |
| GLIDR | ENST00000687919 | 1,9257 |
| NCAPG | ENST00000514176 | 1,9237 |
| ZNF852 | ENST00000436261 | 1,9177 |
| GAS5 | ENST00000689958 | 1,9146 |
| TTC28-AS1 | ENST00000453632 | 1,9126 |
| TLN2 | ENST00000472902 | 1,9099 |
| BST1 | ENST00000514989 | 1,9092 |
| ITGB4 | ENST00000579211 | 1,9081 |
| OSGEP | ENST00000555223 | 1,8974 |
| CTNND1 | ENST00000673661 | 1,8967 |
| LOXL2 | ENST00000520925 | 1,894 |
| RPF1 | ENST00000370656 | 1,8888 |
| ANKRD11 | ENST00000645278 | 1,8862 |
| IL15 | ENST00000477265 | 1,8809 |
| DEPDC5 | ENST00000479261 | 1,8795 |
| TJP2 | ENST00000377245 | 1,8777 |
| PHLDB2 | ENST00000486886 | 1,8772 |
| ITGA3 | ENST00000515147 | 1,8744 |
| ARID1B | ENST00000674298 | 1,8729 |
| ANKS1B | ENST00000549025 | 1,8717 |
| PPP1R12A | ENST00000553081 | 1,8693 |
| SLC35A3 | ENST00000640178 | 1,8691 |
| TSC1 | ENST00000643625 | 1,8629 |
| SARS1 | ENST00000369923 | 1,8612 |
| SUGP1 | ENST00000587716 | 1,8566 |
| ZNF687 | ENST00000324048 | 1,8554 |
| GTF2H1 | ENST00000530496 | 1,855 |
| DHCR7 | ENST00000531364 | 1,854 |
| LRP8 | ENST00000668448 | 1,8528 |
| MKRN1 | ENST00000471104 | 1,8502 |
| DGKB | ENST00000402815 | 1,8482 |
| TXNDC17 | ENST00000574734 | 1,8481 |
| SF3B3 | ENST00000562722 | 1,8479 |
| SLC7A6 | ENST00000566454 | 1,8466 |
| NTNG1 | ENST00000370067 | 1,8456 |
| BAIAP3 | ENST00000324385 | 1,8423 |
| SSBP1 | ENST00000463093 | 1,8411 |
| TOGARAM1 | ENST00000556823 | 1,8403 |
| TPM1 | ENST00000559831 | 1,8403 |
| PKD1P4 | ENST00000433020 | 1,8375 |
| MISP | ENST00000215582 | 1,8319 |
| TAF4B | ENST00000269142 | 1,8314 |
| GPATCH8 | ENST00000587228 | 1,8276 |
| TES | ENST00000496912 | 1,8267 |
| ITGA6 | ENST00000469534 | 1,8259 |
| COL9A2 | ENST00000466267 | 1,8201 |
| MRPL30 | ENST00000409145 | 1,8187 |
| OSBPL1A | ENST00000399441 | 1,8161 |
| CCDC18 | ENST00000401026 | 1,8131 |
| TCP11L1 | ENST00000531632 | 1,813 |
| CBWD5 | ENST00000429800 | 1,8096 |

| | | |
|---|---|---|
| IMP4 | ENST00000428740 | -4,9203 |
| PRDX6-AS1 | ENST00000367716 | -4,9206 |
| SEPTIN7-DT | ENST00000437235 | -4,9237 |
| PIGK | ENST00000359130 | -4,9283 |
| BLMH | ENST00000582749 | -4,9403 |
| TP53 | ENST00000504937 | -4,9457 |
| LMBRD1 | ENST00000649011 | -4,9495 |
| KBTBD11-AS1 | ENST00000650317 | -4,9507 |
| MIR99AHG | ENST00000453910 | -4,9573 |
| RABL2A | ENST00000486403 | -4,963 |
| COASY | ENST00000591779 | -4,9722 |
| VPS35 | ENST00000569950 | -4,9761 |
| MKLN1-AS | ENST00000648242 | -5,0191 |
| MEN1 | ENST00000377313 | -5,0197 |
| MFGE8 | ENST00000558352 | -5,0234 |
| ERO1B | ENST00000688164 | -5,0279 |
| RENBP | ENST00000369997 | -5,0343 |
| TMEM250 | ENST00000561457 | -5,035 |
| AKAP13 | ENST00000559362 | -5,0666 |
| RNF34 | ENST00000554484 | -5,0764 |
| MIA3 | ENST00000470521 | -5,0769 |
| PNCK | ENST00000466638 | -5,0811 |
| TAMM41 | ENST00000486090 | -5,0887 |
| EWSR1 | ENST00000629659 | -5,0897 |
| HM13 | ENST00000469126 | -5,099 |
| TTR | ENST00000237014 | -5,1082 |
| DDX17 | ENST00000479734 | -5,1094 |
| CRYZL2P-SEC16B | ENST00000464428 | -5,1169 |
| SLC37A3 | ENST00000340308 | -5,1185 |
| SMARCA2 | ENST00000324954 | -5,1195 |
| POMGNT1 | ENST00000692202 | -5,1299 |
| DMRTC1 | ENST00000615063 | -5,1334 |
| SLC10A3 | ENST00000651600 | -5,134 |
| PHF21A | ENST00000688570 | -5,1458 |
| BANCR | ENST00000624238 | -5,1485 |
| KMT2C | ENST00000682283 | -5,1683 |
| UBAP2L | ENST00000271877 | -5,1684 |
| DRAM2 | ENST00000539140 | -5,1722 |
| CEP83 | ENST00000551250 | -5,1765 |
| PPFIA1 | ENST00000648755 | -5,1815 |
| LINC01138 | ENST00000614292 | -5,1857 |
| CLCC1 | ENST00000688610 | -5,1953 |
| RBM14 | ENST00000461478 | -5,1982 |
| RPL36 | ENST00000579649 | -5,1995 |
| SMCHD1 | ENST00000584897 | -5,2016 |
| PIK3C2A | ENST00000533645 | -5,2229 |
| MYO1C | ENST00000646049 | -5,2245 |
| CACNA1G | ENST00000506520 | -5,2269 |
| CRY1 | ENST00000546722 | -5,2337 |
| ZKSCAN8 | ENST00000606198 | -5,2352 |
| POLL | ENST00000370162 | -5,2397 |
| ALS2 | ENST00000680939 | -5,2428 |
| EIF2AK3 | ENST00000682892 | -5,2477 |
| SMARCA2 | ENST00000634343 | -5,2559 |
| HERC2P2 | ENST00000613386 | -5,2604 |
| RBM23 | ENST00000557403 | -5,2662 |
| LARP4 | ENST00000518444 | -5,2677 |
| HNRNPU | ENST00000639824 | -5,2738 |
| NUP37 | ENST00000551200 | -5,2812 |
| HECTD2 | ENST00000446394 | -5,2838 |
| SNORD3B-1 | ENST00000577988 | -5,2922 |
| ZDHHC8 | ENST00000405930 | -5,2934 |
| SALL1 | ENST00000690502 | -5,2941 |
| HIPK1 | ENST00000369561 | -5,2946 |
| NPAS3 | ENST00000551008 | -5,305 |

| | | |
|---|---|---|
| CTNND1 | ENST00000534579 | 1,8096 |
| DENND4B | ENST00000485359 | 1,8032 |
| RPL21 | ENST00000272274 | 1,798 |
| PLXNB1 | ENST00000465117 | 1,7978 |
| FANCC | ENST00000289081 | 1,797 |
| HPS1 | ENST00000338546 | 1,7968 |
| MYO5B | ENST00000285039 | 1,7944 |
| FLNA | ENST00000444578 | 1,7929 |
| PDGFC | ENST00000422544 | 1,7902 |
| TYRO3 | ENST00000560992 | 1,7888 |
| PRKRA | ENST00000490501 | 1,783 |
| ADAM9 | ENST00000677165 | 1,7799 |
| CD63 | ENST00000550776 | 1,7745 |
| ASIC2 | ENST00000225823 | 1,7698 |
| GK5 | ENST00000460515 | 1,7694 |
| FKBP15 | ENST00000414250 | 1,7648 |
| SENP3-EIF4A1 | ENST00000614237 | 1,7648 |
| TUBD1 | ENST00000340993 | 1,7567 |
| EPHB4 | ENST00000478459 | 1,7554 |
| RNF216 | ENST00000389900 | 1,752 |
| XAB2 | ENST00000600230 | 1,7501 |
| RABGEF1P1 | ENST00000692179 | 1,7496 |
| TIMM50 | ENST00000598125 | 1,749 |
| VPS13A | ENST00000376646 | 1,7456 |
| YKT6 | ENST00000679310 | 1,7456 |
| MAN1B1 | ENST00000550113 | 1,7424 |
| EMP3 | ENST00000597057 | 1,7399 |
| HAGLR | ENST00000643050 | 1,7352 |
| GARS1 | ENST00000676259 | 1,7314 |
| PTPRZ1 | ENST00000652054 | 1,7275 |
| OLFML2A | ENST00000331715 | 1,7264 |
| SNTG1 | ENST00000642377 | 1,722 |
| EVA1A | ENST00000486489 | 1,7208 |
| ARHGEF35-AS | ENST00000650250 | 1,7194 |
| TMEM222 | ENST00000464720 | 1,7183 |
| RNF31 | ENST00000559882 | 1,7181 |
| PI4KB | ENST00000438243 | 1,7138 |
| WRAP53 | ENST00000534050 | 1,7082 |
| SLC16A1-AS1 | ENST00000420168 | 1,6977 |
| UBE2F | ENST00000480828 | 1,6847 |
| SPC25 | ENST00000479309 | 1,6833 |
| MRPS34 | ENST00000569585 | 1,6828 |
| WDR41 | ENST00000507029 | 1,6817 |
| NCAPH2 | ENST00000418794 | 1,6691 |
| MYRF | ENST00000278836 | 1,6635 |
| PRMT1 | ENST00000534676 | 1,6577 |
| VPS37A | ENST00000521829 | 1,6563 |
| ZMIZ2 | ENST00000265346 | 1,6539 |
| THY1 | ENST00000528295 | 1,6536 |
| GMPPA | ENST00000622191 | 1,6535 |
| RELCH | ENST00000588446 | 1,6471 |
| CBS | ENST00000478709 | 1,6431 |
| RRM2 | ENST00000360566 | 1,6416 |
| HDAC10 | ENST00000470965 | 1,6413 |
| RNH1 | ENST00000533410 | 1,6361 |
| ABCD3 | ENST00000315713 | 1,6327 |
| ADAMTSL5 | ENST00000589839 | 1,6312 |
| PPP2R3C | ENST00000557074 | 1,6271 |
| RRAS2 | ENST00000534746 | 1,6254 |
| NAV1 | ENST00000367296 | 1,6229 |
| CLCN5 | ENST00000376088 | 1,6223 |
| ZNF721 | ENST00000511833 | 1,6175 |
| ZNF880 | ENST00000422689 | 1,615 |
| FRMD5 | ENST00000402883 | 1,6131 |
| SEC24D | ENST00000505134 | 1,611 |

| | | |
|---|---|---|
| NIN | ENST00000673657 | -5,3105 |
| SLIRP | ENST00000555890 | -5,3163 |
| CFTR | ENST00000003084 | -5,3242 |
| AP1B1 | ENST00000317368 | -5,3317 |
| EDRF1 | ENST00000356792 | -5,3495 |
| SMG1P1 | ENST00000446662 | -5,3524 |
| STK11 | ENST00000593219 | -5,3602 |
| USP34-DT | ENST00000603199 | -5,3687 |
| LARGE1 | ENST00000354992 | -5,3737 |
| MMP28 | ENST00000612672 | -5,3759 |
| CTSA | ENST00000678622 | -5,3858 |
| NOTCH2NLR | ENST00000690847 | -5,3909 |
| DNAJC10 | ENST00000681873 | -5,3923 |
| SUGP1 | ENST00000590439 | -5,3954 |
| GALE | ENST00000469556 | -5,4042 |
| TTC3 | ENST00000492275 | -5,4103 |
| PTPN5 | ENST00000358540 | -5,419 |
| STING1 | ENST00000511886 | -5,4193 |
| CLIP4 | ENST00000691852 | -5,4324 |
| APC | ENST00000508624 | -5,4362 |
| PATL2 | ENST00000434130 | -5,4423 |
| RPL22 | ENST00000465387 | -5,4457 |
| GORAB | ENST00000692875 | -5,4513 |
| CTU2 | ENST00000564921 | -5,4562 |
| CLUL1 | ENST00000692774 | -5,4573 |
| TLE4 | ENST00000376544 | -5,4626 |
| HECTD4 | ENST00000550968 | -5,4654 |
| DDHD1 | ENST00000323669 | -5,4714 |
| CELF2 | ENST00000609692 | -5,474 |
| MUC5AC | ENST00000621226 | -5,4759 |
| PDE6B-AS1 | ENST00000598370 | -5,4806 |
| MCRIP2 | ENST00000474840 | -5,5069 |
| ERLEC1 | ENST00000690280 | -5,5127 |
| ACOT7 | ENST00000377860 | -5,5136 |
| TIPIN | ENST00000562124 | -5,5145 |
| TFAP2E | ENST00000682155 | -5,5249 |
| QARS1 | ENST00000636669 | -5,54 |
| ZNF594 | ENST00000399604 | -5,5527 |
| CAPN3 | ENST00000673854 | -5,5533 |
| CNTROB | ENST00000571540 | -5,5534 |
| PANK2 | ENST00000621507 | -5,5574 |
| CHMP1A | ENST00000675309 | -5,5635 |
| SMYD1 | ENST00000419482 | -5,5693 |
| CNTN5 | ENST00000279463 | -5,5757 |
| TRO | ENST00000319167 | -5,5833 |
| TMEM198B | ENST00000636428 | -5,5912 |
| SLC5A7 | ENST00000264047 | -5,5969 |
| COX7A2L | ENST00000482463 | -5,6128 |
| USP8 | ENST00000560379 | -5,6188 |
| NPHP1 | ENST00000417665 | -5,6227 |
| LINC02506 | ENST00000660373 | -5,6259 |
| KRT8 | ENST00000548998 | -5,6338 |
| ITGA4 | ENST00000465522 | -5,6359 |
| STRA6 | ENST00000616000 | -5,6366 |
| BRPF1 | ENST00000497565 | -5,6388 |
| SNHG22 | ENST00000687911 | -5,6455 |
| RBM23 | ENST00000555209 | -5,6733 |
| ETFA | ENST00000559386 | -5,6749 |
| ZFAND2B | ENST00000621130 | -5,6777 |
| ZMYND11 | ENST00000627286 | -5,7112 |
| CD14 | ENST00000302014 | -5,7196 |
| DENND5A | ENST00000681425 | -5,726 |
| PCBP3 | ENST00000681687 | -5,7686 |
| NEB | ENST00000397345 | -5,7698 |
| RBMS1 | ENST00000491781 | -5,7704 |

| | | |
|---|---|---|
| DOCK7 | ENST00000464312 | 1,6095 |
| ABCF3 | ENST00000478288 | 1,6081 |
| HGH1 | ENST00000534255 | 1,607 |
| TRO | ENST00000173898 | 1,5992 |
| EML3 | ENST00000533165 | 1,595 |
| UBE2D3 | ENST00000507845 | 1,5946 |
| C5orf38 | ENST00000515640 | 1,5942 |
| COL4A1 | ENST00000648170 | 1,5919 |
| DYNC1H1 | ENST00000643684 | 1,5874 |
| ADAM9 | ENST00000481873 | 1,5845 |
| ERI1 | ENST00000519292 | 1,584 |
| SWSAP1 | ENST00000312423 | 1,5829 |
| POLR3D | ENST00000518039 | 1,5821 |
| HERPUD1 | ENST00000562914 | 1,5816 |
| PCDHA7 | ENST00000525929 | 1,5795 |
| SPATS2 | ENST00000549412 | 1,5722 |
| SETD6 | ENST00000422445 | 1,5706 |
| SLFNL1-AS1 | ENST00000626479 | 1,5676 |
| COQ8B | ENST00000678404 | 1,5657 |
| PDCD4 | ENST00000483595 | 1,5657 |
| CLPTM1L | ENST00000511268 | 1,5614 |
| CAP1 | ENST00000435719 | 1,5562 |
| GORASP1 | ENST00000695444 | 1,5545 |
| MPDU1 | ENST00000572836 | 1,5544 |
| LINC00461 | ENST00000658935 | 1,5531 |
| DST | ENST00000523967 | 1,5451 |
| CTBP2 | ENST00000494626 | 1,5448 |
| NMRAL1 | ENST00000574733 | 1,5413 |
| SRSF10 | ENST00000473754 | 1,5413 |
| DLAT | ENST00000531306 | 1,5396 |
| TNFRSF19 | ENST00000382258 | 1,5396 |
| SEPTIN2 | ENST00000360051 | 1,5352 |
| ALS2 | ENST00000679701 | 1,5332 |
| PLCE1 | ENST00000685889 | 1,5307 |
| PHGDH | ENST00000369407 | 1,5299 |
| KAZALD1 | ENST00000477267 | 1,5287 |
| TAF9 | ENST00000687205 | 1,5282 |
| RASL10A | ENST00000401450 | 1,5279 |
| SHMT2 | ENST00000556737 | 1,5198 |
| RBM39 | ENST00000374038 | 1,5179 |
| PLXNB1 | ENST00000449094 | 1,5168 |
| RPS9 | ENST00000391751 | 1,5167 |
| TMUB2 | ENST00000587326 | 1,5142 |
| RPL37A | ENST00000624436 | 1,5137 |
| DCAF6 | ENST00000470721 | 1,5136 |
| GON4L | ENST00000361040 | 1,512 |
| NPM1 | ENST00000678774 | 1,5112 |
| MST1 | ENST00000494809 | 1,5087 |
| PABPC1L | ENST00000465761 | 1,5084 |
| ZC3H14 | ENST00000556000 | 1,5073 |
| UBE2D3 | ENST00000321805 | 1,5051 |
| RPSA | ENST00000475346 | 1,5033 |
| MRPL1 | ENST00000506674 | 1,5023 |
| GARS1 | ENST00000674734 | 1,5005 |
| NUP153-AS1 | ENST00000606771 | 1,4988 |
| ANXA5 | ENST00000513523 | 1,4959 |
| PHTF1 | ENST00000369600 | 1,4941 |
| PDCD5 | ENST00000586316 | 1,492 |
| BPTF | ENST00000578307 | 1,4917 |
| RDH11 | ENST00000556692 | 1,4905 |
| DGAT1 | ENST00000531896 | 1,49 |
| FN1 | ENST00000426059 | 1,4876 |
| TSPAN4 | ENST00000530404 | 1,486 |
| ERCC8 | ENST00000675042 | 1,4843 |
| NAA38 | ENST00000570555 | 1,4837 |

| | | |
|---|---|---|
| LMO7 | ENST00000497947 | -5,7776 |
| CES3 | ENST00000303334 | -5,7787 |
| GLB1 | ENST00000307377 | -5,7847 |
| SON | ENST00000421541 | -5,8113 |
| KIF27 | ENST00000413982 | -5,825 |
| LRSAM1 | ENST00000676137 | -5,8263 |
| ZMAT1 | ENST00000458570 | -5,8283 |
| TMEM176A | ENST00000494349 | -5,8309 |
| STRCP1 | ENST00000509801 | -5,8338 |
| DLC1 | ENST00000510250 | -5,8469 |
| GPATCH8 | ENST00000635257 | -5,8472 |
| SH3YL1 | ENST00000403658 | -5,8489 |
| DYNC1I2 | ENST00000479806 | -5,8947 |
| AQP2 | ENST00000199280 | -5,9284 |
| TNNT3 | ENST00000493234 | -5,9286 |
| CD46 | ENST00000360212 | -5,9394 |
| SLC6A1 | ENST00000645974 | -5,9405 |
| BARHL1 | ENST00000263610 | -5,9445 |
| CYB5D2 | ENST00000575251 | -5,9723 |
| KLF3-AS1 | ENST00000665526 | -5,9724 |
| LINC02899 | ENST00000512559 | -6,0172 |
| MRPL28 | ENST00000447696 | -6,0377 |
| ACP3 | ENST00000351273 | -6,0424 |
| OTX2 | ENST00000672264 | -6,0796 |
| LGI4 | ENST00000587780 | -6,1143 |
| AOPEP | ENST00000478473 | -6,1581 |
| CCDC91 | ENST00000545336 | -6,1624 |
| TRDN | ENST00000542443 | -6,205 |
| RNVU1-18 | ENST00000384010 | -6,2136 |
| PARD6G-AS1 | ENST00000692837 | -6,2188 |
| SNHG17 | ENST00000662982 | -6,2509 |
| MAP3K19 | ENST00000486077 | -6,2704 |
| BARHL2 | ENST00000370445 | -6,2729 |
| NUBP1 | ENST00000433392 | -6,293 |
| SLC9A6 | ENST00000370698 | -6,3023 |
| MYO18B | ENST00000543971 | -6,3435 |
| STXBP1 | ENST00000636962 | -6,3483 |
| AP3B2 | ENST00000681452 | -6,3518 |
| ZNF883 | ENST00000639196 | -6,368 |
| RNF43 | ENST00000407977 | -6,3832 |
| EFCAB6 | ENST00000262726 | -6,4463 |
| NFKBIL1 | ENST00000473655 | -6,4498 |
| TMEM30A-DT | ENST00000661161 | -6,4554 |
| SERPINI2 | ENST00000471111 | -6,4723 |
| DYNC1H1 | ENST00000643437 | -6,4968 |
| DES | ENST00000483395 | -6,5305 |
| ADH4 | ENST00000265512 | -6,5372 |
| ADAM15 | ENST00000472434 | -6,5383 |
| LRSAM1 | ENST00000676318 | -6,5959 |
| IDNK | ENST00000533522 | -6,6474 |
| PBRM1 | ENST00000394830 | -6,8741 |
| NF1P8 | ENST00000415351 | -7,0939 |
| FEZF2 | ENST00000486811 | -7,1023 |
| OPRK1 | ENST00000673285 | -7,5696 |
| TTN | ENST00000589042 | -7,8241 |
| MRPS27 | ENST00000695298 | -18,8583 |

| | | |
|---|---|---|
| EPB41L4A-AS1 | ENST00000427306 | 1,4801 |
| YDJC | ENST00000464015 | 1,4798 |
| NBPF15 | ENST00000614785 | 1,4793 |
| SCUBE2 | ENST00000649792 | 1,4716 |
| RETREG1 | ENST00000682808 | 1,471 |
| APP | ENST00000448850 | 1,4695 |
| EIF1AD | ENST00000526451 | 1,4695 |
| NMRAL1 | ENST00000573520 | 1,466 |
| PHACTR4 | ENST00000632202 | 1,4623 |
| LOXL1-AS1 | ENST00000688623 | 1,4607 |
| LPIN1 | ENST00000460096 | 1,4556 |
| LIX1 | ENST00000512378 | 1,45 |
| NEDD4L | ENST00000676301 | 1,4493 |
| NAA50 | ENST00000481432 | 1,4471 |
| VPS35 | ENST00000562420 | 1,447 |
| CDC6 | ENST00000649662 | 1,4444 |
| JKAMP | ENST00000556985 | 1,4413 |
| FAM111A-DT | ENST00000531708 | 1,4408 |
| CD151 | ENST00000530726 | 1,4391 |
| HMSD | ENST00000408945 | 1,4376 |
| LRRC37BP1 | ENST00000412831 | 1,4371 |
| HECTD3 | ENST00000486296 | 1,4351 |
| GMPPB | ENST00000495627 | 1,4345 |
| RAP1GAP | ENST00000542643 | 1,4307 |
| WTIP | ENST00000585928 | 1,43 |
| CEP104 | ENST00000428079 | 1,4267 |
| KDM2B | ENST00000446152 | 1,4233 |
| PPP1R16A | ENST00000533088 | 1,4174 |
| REXO5 | ENST00000261377 | 1,417 |
| EVI5L | ENST00000538904 | 1,4066 |
| VKORC1 | ENST00000300851 | 1,4016 |
| NPRL3 | ENST00000621703 | 1,3989 |
| SEC14L2 | ENST00000416523 | 1,3959 |
| GPBP1L1 | ENST00000487436 | 1,3939 |
| SEC62 | ENST00000487736 | 1,3893 |
| TPM4 | ENST00000657915 | 1,3853 |
| LENG8 | ENST00000616932 | 1,3851 |
| GARS1 | ENST00000485784 | 1,3728 |
| VPS53 | ENST00000681217 | 1,3724 |
| PCGF2 | ENST00000611883 | 1,3718 |
| COCH | ENST00000216361 | 1,3687 |
| KATNBL1 | ENST00000560108 | 1,3654 |
| BORCS8-MEF2 | ENST00000602804 | 1,3641 |
| KCNG1 | ENST00000439216 | 1,3637 |
| UBA5 | ENST00000473651 | 1,3583 |
| PTP4A2 | ENST00000489313 | 1,3506 |
| GYPC | ENST00000259254 | 1,3479 |
| FRY | ENST00000641614 | 1,3469 |
| TALDO1 | ENST00000530666 | 1,3438 |
| CEBPG | ENST00000585933 | 1,3433 |
| LRRC57 | ENST00000323443 | 1,3402 |
| TAF10 | ENST00000532344 | 1,3374 |
| PCOLCE2 | ENST00000295992 | 1,3364 |
| TPR | ENST00000467810 | 1,3321 |
| PCBP2 | ENST00000546463 | 1,332 |
| BTN2A1 | ENST00000541522 | 1,3313 |
| ZFYVE27 | ENST00000370613 | 1,331 |
| RPS15A | ENST00000576436 | 1,3303 |
| KHDRBS3 | ENST00000521461 | 1,3301 |
| ZNF324 | ENST00000593925 | 1,33 |
| NRGN | ENST00000284292 | 1,3224 |
| OGFOD1 | ENST00000565682 | 1,3219 |
| HMCES | ENST00000509042 | 1,3208 |
| PSMD13 | ENST00000532025 | 1,3158 |
| SRD5A3 | ENST00000677177 | 1,3138 |

| | | |
|---|---|---|
| MPI | ENST00000565576 | 1,3078 |
| GTF2H1 | ENST00000526630 | 1,3035 |
| NUP42 | ENST00000497500 | 1,2984 |
| COPS7A | ENST00000539735 | 1,2983 |
| TMEM44 | ENST00000477651 | 1,2936 |
| PARG | ENST00000614063 | 1,2935 |
| CKMT2-AS1 | ENST00000690798 | 1,2892 |
| PPL | ENST00000345988 | 1,2868 |
| LMNA | ENST00000368298 | 1,2867 |
| C7orf50 | ENST00000397098 | 1,2857 |
| ARFGAP1 | ENST00000522959 | 1,2847 |
| RBPJ | ENST00000514380 | 1,2791 |
| CYB5D2 | ENST00000577075 | 1,2736 |
| SBF1 | ENST00000690990 | 1,2706 |
| EIF3C | ENST00000566866 | 1,2701 |
| COL4A2 | ENST00000480609 | 1,2656 |
| BCR | ENST00000475025 | 1,2636 |
| SRM | ENST00000475189 | 1,256 |
| CALD1 | ENST00000466704 | 1,2558 |
| APEX1 | ENST00000556296 | 1,2548 |
| IP6K1 | ENST00000395238 | 1,2542 |
| CSNK2B | ENST00000481269 | 1,2476 |
| IMPDH1 | ENST00000496200 | 1,2467 |
| ITM2C | ENST00000335005 | 1,2448 |
| TEDC1 | ENST00000546492 | 1,2444 |
| IMP4 | ENST00000409649 | 1,242 |
| AGK | ENST00000496784 | 1,2399 |
| SALL1 | ENST00000685868 | 1,2381 |
| PLOD3 | ENST00000424135 | 1,2378 |
| RPS6KB2 | ENST00000528964 | 1,2354 |
| OAT | ENST00000471127 | 1,2338 |
| PIK3CD | ENST00000377346 | 1,2338 |
| ASL | ENST00000395332 | 1,2315 |
| HNRNPM | ENST00000597081 | 1,2277 |
| TCAF2 | ENST00000441159 | 1,2249 |
| CCM2 | ENST00000544363 | 1,2236 |
| FCF1 | ENST00000553615 | 1,2233 |
| PCSK7 | ENST00000524507 | 1,2224 |
| NCDN | ENST00000373253 | 1,2194 |
| PICALM | ENST00000532603 | 1,219 |
| MAST2 | ENST00000467367 | 1,2158 |
| POLD1 | ENST00000600859 | 1,2119 |
| C11orf45 | ENST00000524878 | 1,2069 |
| MYO6 | ENST00000671923 | 1,2053 |
| SMAD7 | ENST00000545051 | 1,2017 |
| RPL3 | ENST00000420536 | 1,2016 |
| GUSBP3 | ENST00000513408 | 1,1996 |
| SGTA | ENST00000678109 | 1,1996 |
| ZNF559 | ENST00000317221 | 1,1972 |
| ERRFI1 | ENST00000487559 | 1,1958 |
| FGD3 | ENST00000416701 | 1,1951 |
| ARPC4 | ENST00000417500 | 1,1911 |
| SETDB1 | ENST00000481219 | 1,1873 |
| DERA | ENST00000526521 | 1,1857 |
| GUSBP13 | ENST00000506490 | 1,1747 |
| AP3D1 | ENST00000586370 | 1,1736 |
| PABPC4 | ENST00000676863 | 1,1714 |
| CUTA | ENST00000482684 | 1,1671 |
| MITD1 | ENST00000409107 | 1,1648 |
| SERPINH1 | ENST00000525876 | 1,1556 |
| HYOU1 | ENST00000694930 | 1,1438 |
| HOXD8 | ENST00000450510 | 1,1425 |
| COPZ1 | ENST00000549043 | 1,1392 |
| SSR3 | ENST00000463503 | 1,1374 |
| SLCO4C1 | ENST00000310954 | 1,1355 |

| | | |
|---|---|---|
| CDKN3 | ENST00000335183 | 1,1348 |
| HEATR5A | ENST00000538864 | 1,1339 |
| TCOF1 | ENST00000394269 | 1,1337 |
| COL4A1 | ENST00000647632 | 1,1256 |
| ATF7 | ENST00000548118 | 1,125 |
| TBC1D17 | ENST00000600354 | 1,1226 |
| PTCD3 | ENST00000484203 | 1,121 |
| NPAS2 | ENST00000474550 | 1,1153 |
| IKBKG | ENST00000689906 | 1,0959 |
| BAX | ENST00000513217 | 1,0881 |
| LZTS2 | ENST00000370223 | 1,0871 |
| SPON2 | ENST00000509697 | 1,0816 |
| NBPF15 | ENST00000577412 | 1,0811 |
| SSU72 | ENST00000378726 | 1,0791 |
| KLC1 | ENST00000445352 | 1,0767 |
| EIF2B4 | ENST00000418146 | 1,0756 |
| LIG4 | ENST00000692222 | 1,0751 |
| AFG3L1P | ENST00000421164 | 1,0671 |
| GALE | ENST00000459934 | 1,0631 |
| FAM126A | ENST00000432176 | 1,0608 |
| ADRM1 | ENST00000465805 | 1,0575 |
| URI1 | ENST00000360605 | 1,0574 |
| GPSM2 | ENST00000435987 | 1,0572 |
| SLC52A2 | ENST00000530047 | 1,0527 |
| PHKG2 | ENST00000563913 | 1,0519 |
| TAGLN2 | ENST00000478033 | 1,0492 |
| UNC5B | ENST00000373192 | 1,0477 |
| EIF3J | ENST00000535391 | 1,042 |
| LMNA | ENST00000675431 | 1,0409 |
| GSC | ENST00000238558 | 1,0401 |
| ZIK1 | ENST00000597850 | 1,0334 |
| NOP56 | ENST00000492135 | 1,033 |
| MRPL42 | ENST00000552326 | 1,0295 |
| B4GALT2 | ENST00000481924 | 1,0289 |
| BMP6 | ENST00000283147 | 1,0221 |
| DLK2 | ENST00000372488 | 1,0219 |
| EHBP1 | ENST00000471179 | 1,0216 |
| ZBTB44 | ENST00000527478 | 1,021 |
| KCTD20 | ENST00000536244 | 1,0122 |
| TCEA2 | ENST00000415602 | 1,0088 |
| ZDHHC6 | ENST00000684173 | 1,0074 |
| WDR1 | ENST00000502702 | 1,0052 |
| S100A6 | ENST00000496817 | 1,0034 |
| AARS1 | ENST00000675297 | 1,0031 |

**Supplementary Table 4 B. GO Analysis**

| ID | Description | GeneRatio | BgRatio | pvalue | p.adjust | qvalue | geneID |
|---|---|---|---|---|---|---|---|
| GO:0090150 | establishment of protein localization to membrane | 53/1286 | 379/21081 | 1,39E-08 | 6,38E-05 | 5,98E-05 | SSR2/RPS8/RPL5/RAB3GAP2/MACF1/RPS7/RPL37A/ATG3/SSR3/TP63/RPSA/SEC62/GORASP1/COX18/KIF13A/FYN/AFDN/BAG6/SEC61G/AGK/TCAF2/VPS37A/LYPLA1/SCRIB/RPS20/HSPA5/RPLP2/PAK1/BBS1/ZDHHC6/OPTN/TAMALIN/CACNB3/RPL21/RPS29/RPS15A/VPS35/RPS2/RAB34/NMT1/ERBB2/MTCL1/ARFRP1/RAB8A/RPL18A/RAB11B/RPS11/SGTA/RPS9/BAX/PDCD5/RPL3/GET1 |
| GO:0006605 | protein targeting | 62/1286 | 483/21081 | 2,30E-08 | 6,38E-05 | 5,98E-05 | SSR2/TOMM40L/RPS8/RPL5/RPS7/MPV17/GCC2/RPL37A/ATG3/RAB7A/SSR3/RPSA/SEC62/GRPEL1/UBE2D3/FYN/ECI2/SEC61G/AGK/AP4M1/PMPCB/TCAF2/VPS37A/CLU/RPS20/VPS13A/DNLZ/HSPA5/CRAT/CAT/RPLP2/PAK1/ATG13/SPCS2/ZDHHC6/CACNB3/RPL21/RPS29/RNF31/ZFAND6/RPS15A/NUDT7/DECR2/RPS2/HGS/NPEPPS/ERBB2/GABARAP/SREBF1/USP36/MTCL1/RPL18A/TIMM50/AP3D1/RPS11/SGTA/RPS9/PDCD5/RPL3/HPS4/UBE2L3/GET1 |
| GO:0034329 | cell junction assembly | 56/1286 | 471/21081 | 1,34E-06 | 0,00248896 | 0,00233524 | PATJ/COL16A1/MACF1/CDC42/FBLIM1/OBSL1/FN1/ITGA6/PDCD6IP/CTNNB1/PHLDB2/RYK/PLXNB1/CLASP2/SETD5/DLG1/WDR1/CDH10/PCDHB10/AFDN/DST/PPP1R9A/ACHE/SLITRK4/FGF13/SLITRK2/FLNA/CLDN2/GPRASP2/NLGN3/PTK2/TSC1/BDNF/CD151/CTNND1/THY1/PTPRO/RAP1B/CD9/ARHGEF7/TJP1/TLN2/VPS35/CLDN6/ITGB4/ASIC2/ACTG1/RAMP2/TBCD/FBF1/LAMA3/EPB41L3/SMAD7/CAMSAP3/BCR/APP |
| GO:0006413 | translational initiation | 32/1286 | 220/21081 | 4,31E-06 | 0,0059865 | 0,00561677 | TPR/RPS8/RPL5/EIF4G3/RPS7/EIF2B4/EIF4E2/RPL37A/RPSA/NPM1/EIF4H/PABPC1/RPS20/GLE1/RPLP2/EIF1AD/PPP1CA/RPS6KB2/RPL21/RPS29/EIF3J/RPS15A/EIF3C/RPS2/EIF4A1/EIF6/RPL18A/RPS11/RPS9/RPL3/EIF3L/ATF4 |
| GO:0016197 | endosomal transport | 36/1286 | 265/21081 | 5,55E-06 | 0,00616894 | 0,00578795 | ARL4C/CCDC93/SNX17/ALS2/RAB6D/GCC2/RAB7A/TBC1D14/REPS1/YKT6/VPS37A/SCRIB/DENND1A/PICALM/EHD1/VPS51/WASHC2C/GBF1/TAMALIN/PHETA1/RAB35/VPS36/HEATR5A/SNX1/EHD4/VPS35/BAIAP3/SNF8/HGS/VPS53/MYO5B/ARFRP1/RAB8A/AP3D1/RAB11B/TBC1D17 |
| GO:0048193 | Golgi vesicle transport | 48/1286 | 407/21081 | 9,36E-06 | 0,00716557 | 0,00672303 | MACF1/COPA/TRAPPC12/CCDC93/DYNC1I2/RAB6D/GCC2/SEC13/ANKRD28/GORASP1/TBC1D14/SEC24D/NKD2/LMAN2/KIF13A/AP4M1/YKT6/LYPLA1/ANK1/NRBP2/VPS13A/BBS1/HYOU1/VPS51/GBF1/OPTN/MON2/KIF5A/COPZ1/GOLGA3/SCFD1/DYNC1H1/MIA2/KLC1/SNAP23/SCAMP2/SNX1/STX4/COPZ2/RAB34/SPIRE1/ARFGAP1/ARFRP1/RAB8A/PPP6R1/AP3D1/DNM2/NAPA |
| GO:0007163 | establishment or maintenance of cell polarity | 32/1286 | 230/21081 | 1,10E-05 | 0,00716557 | 0,00672303 | DOCK7/GPSM2/CAP1/LMNA/CDC42/RND3/MAP2/PARD3B/PDCD6IP/CLASP2/DLG1/WDR1/PTK7/DST/KRIT1/FGF13/ANK1/SCRIB/PTK2/TCIRG1/PAK1/PDLIM1/GBF1/RAP1B/NCKAP1L/RNF41/FBF1/CYTH1/MTCL1/MISP/BRSK1/CAMSAP3 |
| GO:0007229 | integrin-mediated signaling pathway | 21/1286 | 120/21081 | 1,11E-05 | 0,00716557 | 0,00672303 | PHACTR4/ADAM15/COL16A1/CDC42/FN1/ITGA6/BST1/DST/CCM2/FLNA/ADAM9/PTK2/ADAMTS13/THY1/CD63/BCAR1/ITGA3/ITGB4/NME2/LAMA3/LAMA5 |
| GO:0070831 | basement membrane assembly | 7/1286 | 15/21081 | 1,29E-05 | 0,00716557 | 0,00672303 | NTNG1/PHLDB2/CLASP2/PLOD3/LAMB1/PHLDB1/RAMP2 |
| GO:0051648 | vesicle localization | 33/1286 | 244/21081 | 1,47E-05 | 0,00745224 | 0,00699199 | BBS5/MAP2/CTNNB1/SEC13/RAB7A/ANKRD28/CLASP2/GORASP1/SEC24D/KIF13A/DTNBP1/MYO6/YKT6/NLGN3/IKBKG/SCRIB/TCIRG1/PICALM/GBF1/KIF5A/SCFD1/DYNC1H1/SNAP23/STARD3/SYT4/MYO5B/PPP6R1/BRSK1/AP3D1/RAB11B/TPGS1/DNM2/NAPA |
| GO:1903332 | regulation of protein folding | 6/1286 | 12/21081 | 3,42E-05 | 0,01293743 | 0,01213841 | DNAJB2/HSPA5/GRN/SGTA/PDCD5/ST13 |
| GO:0030198 | extracellular matrix organization | 49/1286 | 442/21081 | 3,91E-05 | 0,01354306 | 0,01270664 | COL9A2/CTSK/NTNG1/ADAM15/COL16A1/FN1/MPV17/ITGA6/COL8A1/PHLDB2/CLASP2/VCAN/SPARC/DDR1/SERAC1/PLOD3/LAMB1/COL1A2/COL4A5/LOXL2/BMP1/PTK2/ADAMTS13/SERPINH1/PHLDB1/CAPN1/NFKB2/KAZALD1/COL13A1/MMP19/POSTN/COL4A2/COL4A1/ANXA2/ACAN/ITGA3/ITGB4/RAMP2/NF1/TTR/LAMA3/LAMA5/COL9A3/ADAMTS10/BCL3/ADAMTSL5/MMP11/JAM2/APP |
| GO:0043062 | extracellular structure organization | 49/1286 | 443/21081 | 4,14E-05 | 0,01354306 | 0,01270664 | COL9A2/CTSK/NTNG1/ADAM15/COL16A1/FN1/MPV17/ITGA6/COL8A1/PHLDB2/CLASP2/VCAN/SPARC/DDR1/SERAC1/PLOD3/LAMB1/COL1A2/COL4A5/LOXL2/BMP1/PTK2/ADAMTS13/SERPINH1/PHLDB1/CAPN1/NFKB2/KAZALD1/COL13A1/MMP19/POSTN/COL4A2/COL4A1/ANXA2/ACAN/ITGA3/ITGB4/RAMP2/NF1/TTR/LAMA3/LAMA5/COL9A3/ADAMTS10/BCL3/ADAMTSL5/MMP11/JAM2/APP |

| | | | | | | | |
|---|---|---|---|---|---|---|---|
| GO:0051650 | establishment of vesicle localization | 30/1286 | 225/21081 | 4,65E-05 | 0,01434482 | 0,01345888 | BBS5/MAP2/CTNNB1/SEC13/RAB7A/ANKRD28/CLASP2/GORASP1/SEC24D/KIF13A/DTNBP1/MYO6/YKT6/IKBKG/SCRIB/TCIRG1/PICALM/GBF1/KIF5A/SCFD1/DYNC1H1/SNAP23/STARD3/SYT4/MYO5B/PPP6R1/AP3D1/RAB11B/DNM2/NAPA |
| GO:0018095 | protein polyglutamylation | 6/1286 | 13/21081 | 6,02E-05 | 0,01707977 | 0,01602492 | TTLL7/TTLL4/CEP41/TTLL6/TPGS2/TPGS1 |
| GO:0010822 | positive regulation of mitochondrion organization | 21/1286 | 134/21081 | 6,15E-05 | 0,01707977 | 0,01602492 | TP63/UBE2D3/HDAC6/DDHD2/ATG13/ZDHHC6/OPTN/CAMKK2/ADCK1/RNF31/HIF1A/VPS35/NMT1/NPEPPS/USP36/SPIRE1/BAX/PDCD5/PLA2G6/HPS4/UBE2L3 |
| GO:0031589 | cell-substrate adhesion | 44/1286 | 393/21081 | 7,56E-05 | 0,01911035 | 0,01793009 | AJAP1/TACSTD2/NTNG1/SRGAP2/ADAM15/COL16A1/MACF1/CDC42/SNED1/FN1/ITGA6/CTNNB1/COL8A1/PHLDB2/CLASP2/BST1/DDR1/SGCE/LAMB1/FLNA/ADAM9/PTK2/TMEM8B/ADAMTS13/TSC1/MEN1/THY1/COL13A1/WASHC2C/PTPRO/CD63/POSTN/ARHGEF7/TYRO3/ITGA3/ITGB4/ACTG1/P4HB/TBCD/NF1/LAMA5/CAMSAP3/DNM2/BCR |
| GO:0034332 | adherens junction organization | 14/1286 | 71/21081 | 8,36E-05 | 0,02020689 | 0,01895891 | NECTIN4/CDC42/CTNNB1/CDH10/BMP6/AFDN/CTNND1/NUMB/TJP1/KIFC3/RAMP2/TBCD/SMAD7/CAMSAP3 |
| GO:0044839 | cell cycle G2/M phase transition | 37/1286 | 317/21081 | 0,0001179 | 0,02731138 | 0,02562463 | PPP1R12B/DYNC1I2/MTA3/CEP70/PSMD6/CEP63/NPM1/CDC25C/CEP41/AKAP9/FHL1/HAUS6/CNTRL/USP47/PSMD13/MASTL/OPTN/PPP1R12A/FBXL3/KCNH5/PSME2/DYNC1H1/PSMA6/TICRR/PKMYT1/CDC6/CTC1/PSME3/RAD51C/PSMF1/BABAM1/BRSK1/AKAP8/DNM2/CSNK1E/CHEK2/APP |
| GO:0006457 | protein folding | 31/1286 | 255/21081 | 0,0001977 | 0,03708491 | 0,03479454 | NFYC/PPIL3/DNAJB2/NGLY1/GRPEL1/HSPA9/CANX/TTC1/CSNK2B/CLU/DNLZ/HSPA5/ERP44/RIC3/CHORDC1/HSP90B1/UNC45A/MESD/PPIB/TRAP1/CCT6B/RGS9/MPDU1/P4HB/TBCD/GRN/DNAJC7/SGTA/PDCD5/ALG12/ST13 |
| GO:0010810 | regulation of cell-substrate adhesion | 29/1286 | 233/21081 | 0,0002094 | 0,03713951 | 0,03484576 | AJAP1/TACSTD2/ADAM15/COL16A1/MACF1/CDC42/FN1/ITGA6/COL8A1/PHLDB2/CLASP2/BST1/DDR1/FLNA/PTK2/TSC1/MEN1/THY1/WASHC2C/PTPRO/POSTN/ARHGEF7/ITGA3/ACTG1/P4HB/TBCD/NF1/CAMSAP3/DNM2 |
| GO:0032388 | positive regulation of intracellular transport | 29/1286 | 234/21081 | 0,0002254 | 0,03728422 | 0,03498154 | EFCAB7/TPR/CDC42/MAP2/TP63/UBE2D3/RIOK2/FYN/TCAF2/FLNA/EHD1/PAK1/ATG13/RAPGEF3/CACNB3/RNF31/DYNC1H1/ANXA2/RIPOR1/NUTF2/NMT1/NPEPPS/ERBB2/USP36/MTCL1/OAZ1/PDCD5/HPS4/UBE2L3 |
| GO:0006479 | protein methylation | 26/1286 | 203/21081 | 0,0002773 | 0,03849744 | 0,03611983 | ARID4B/LMNA/KDM1A/SETDB1/FAM98A/CTNNB1/SATB1/SETD5/TET2/H1-4/NELFE/ATRX/PCMTD1/NTMT1/KDM4C/PHF19/PRMT3/TRMT112/MEN1/EEF1AKMT2/ETFBKMT/PRMT5/SETD6/SUPT6H/PRMT1/SETD4 |
| GO:0090316 | positive regulation of intracellular protein transport | 25/1286 | 192/21081 | 0,0002792 | 0,03849744 | 0,03611983 | EFCAB7/TPR/CDC42/TP63/UBE2D3/RIOK2/FYN/TCAF2/FLNA/PAK1/ATG13/RAPGEF3/CACNB3/RNF31/RIPOR1/NUTF2/NMT1/NPEPPS/ERBB2/USP36/MTCL1/OAZ1/PDCD5/HPS4/UBE2L3 |
| GO:0010769 | regulation of cell morphogenesis involved in differentiation | 37/1286 | 331/21081 | 0,0002811 | 0,03849744 | 0,03611983 | TACSTD2/LRP8/BARHL2/MACF1/CDC42/ABI2/OBSL1/FN1/MAP2/RYK/PLXNB1/GORASP1/CDKL3/CAMK2B/PPP1R9A/FGF13/FLNA/GPRASP2/NLGN3/PTK2/NSMF/BDNF/PAK1/THY1/ZFYVE27/WASHC2C/PTPRO/FBXW8/POSTN/ARHGEF7/IST1/ARHGDIA/P4HB/BAIAP2/NEDD4L/BRSK1/DNM2 |
| GO:0150115 | cell-substrate junction organization | 17/1286 | 108/21081 | 0,0002831 | 0,03849744 | 0,03611983 | COL16A1/MACF1/FN1/ITGA6/PHLDB2/CLASP2/DST/PTK2/TSC1/CD151/THY1/ARHGEF7/ITGB4/ACTG1/LAMA3/CAMSAP3/BCR |
| GO:0033539 | fatty acid beta-oxidation using acyl-CoA dehydrogenase | 5/1286 | 11/21081 | 0,0002839 | 0,03849744 | 0,03611983 | ACAD11/ETFBKMT/IVD/ETFA/ACADVL |
| GO:0009161 | ribonucleoside monophosphate metabolic process | 11/1286 | 53/21081 | 0,0002962 | 0,03859501 | 0,03621137 | UCK2/CAD/IMPDH2/DLG1/IMPDH1/PRPS1/TJP2/NT5C2/GMPR2/PFAS/ADSL |
| GO:0001881 | receptor recycling | 9/1286 | 37/21081 | 0,0002985 | 0,03859501 | 0,03621137 | ALS2/CAMLG/ACHE/SCRIB/ARAP1/OPTN/PHETA1/ANXA2/RAB11B |
| GO:0000086 | G2/M transition of mitotic cell cycle | 34/1286 | 298/21081 | 0,0003346 | 0,04132824 | 0,0387758 | PPP1R12B/DYNC1I2/MTA3/CEP70/PSMD6/CEP63/CDC25C/CEP41/AKAP9/FHL1/HAUS6/CNTRL/USP47/PSMD13/MASTL/OPTN/PPP1R12A/FBXL3/KCNH5/PSME2/DYNC1H1/PSMA6/TICRR/PKMYT1/CDC6/CTC1/PSME3/RAD51C/PSMF1/BRSK1/DNM2/CSNK1E/CHEK2/APP |
| GO:0000460 | maturation of 5.8S rRNA | 9/1286 | 38/21081 | 0,0003702 | 0,04321716 | 0,04054806 | RPF1/WDR12/EXOSC7/NSA2/ERI1/UTP20/FCF1/ERI2/EIF6 |

| | | | | | | | |
|---|---|---|---|---|---|---|---|
| GO:0045197 | establishment or maintenance of epithelial cell apical/basal polarity | 10/1286 | 46/21081 | 0,0003732 | 0,04321716 | 0,04054806 | CDC42/PARD3B/PDCD6IP/DLG1/WDR1/PTK7/ANK1/SCRIB/MTCL1/CAMSAP3 |
| GO:0030859 | polarized epithelial cell differentiation | 7/1286 | 24/21081 | 0,0004266 | 0,04839483 | 0,04540596 | AJAP1/CDC42/TP63/PTK7/AHI1/SCRIB/CAMSAP3 |
