## Supplementary Tables 5, 6, 7 for "Functional validation of allele-specific LMNB1 silencing in patient-derived astrocytes as a therapeutic option for Autosomal Dominant Leukodystrophy"

**Supplementary Table 5. Media Composition**

| Medium | Composition | Concentration | Source | Stock number |
|---|---|---|---|---|
| Neural induction Medium (NIM) | DMEM-F12 | 1:100 | ThermoFisher | 31330038 |
| | GlutaMAX | 1:100 | ThermoFisher | 35050038 |
| | MEM NEEA | 1:1000 | ThermoFisher | 11140050 |
| | 2-Mercaptoethanol | 25mg/mL | ThermoFisher | 21985023 |
| | Human Insulin | 10mM | Sigma Merck | I3536 |
| | SB431542 | 250nM | Tocris | 1614/10 |
| | LDN193189 | 100nM | Tocris | 6053/10 |
| | Retinoic Acid | 1:100 | Sigma Merck | R2625 |
| | PenStrep | 1:100 | Sigma Merck | P4333 |
| N2 Medium | DMEM F-12 | 1:100 | ThermoFisher | 31330038 |
| | GlutaMAX | 1:100 | ThermoFisher | 35050038 |
| | MEM NEEA | 1:1000 | ThermoFisher | 11140050 |
| | 2-Mercaptoethanol | 1:100 | ThermoFisher | 21985023 |
| | N2 supplement | 1mM | ThermoFisher | 17502001 |
| | Smoothened agonist | 100nM | Tocris | 6390/1 |
| | Retinoic Acid | 1:100 | Sigma Merck | R2625 |
| | PenStrep | 1:100 | Sigma Merck | P4333 |
| N2B27 Medium | DMEM F-12 | 1:100 | ThermoFisher | 31330038 |
| | GlutaMAX | 1:100 | ThermoFisher | 35050038 |
| | MEM NEEA | 1:1000 | ThermoFisher | 11140050 |
| | 2-Mercaptoethanol | 1:100 | ThermoFisher | 21985023 |
| | Human Insulin | 1:100 | Sigma Merck | I3536 |
| | N2 supplement | 1:50 | ThermoFisher | 17502001 |
| | B27 supplement | 1mM | ThermoFisher | 17504044 |
| | Smoothened agonist | 100nM | Tocris | 6390/1 |
| | Retinoic Acid | 1:100 | Sigma Merck | R2625 |
| | PenStrep | 1:100 | Sigma Merck | P4333 |
| PDGF Medium | DMEM F-12 | 1:100 | ThermoFisher | 31330038 |
| | GlutaMAX | 1:100 | ThermoFisher | 35050038 |
| | MEM NEEA | 1:1000 | ThermoFisher | 11140050 |
| | 2-Mercaptoethanol | 1:100 | ThermoFisher | 21985023 |
| | Human Insulin | 1:100 | Sigma Merck | I3536 |
| | N2 supplement | 1:50 | ThermoFisher | 17502001 |
| | B27 supplement | 10ng/mL | ThermoFisher | 17504044 |
| | PDGFaa | 10ng/mL | ThermoFisher | PHG0035 |
| | IGF-1 | 5ng/mL | ThermoFisher | PHG0078 |
| | HGF | 10ng/mL | ThermoFisher | PHG0254 |
| | NT3 | 60ng/mL | ThermoFisher | PHC7036 |
| | T3 | 100ng/mL | Sigma Merck | T2877 |
| | Biotin | 1mM | Sigma Merck | B4639 |
| | cAMP | 1:100 | Sigma Merck | A9501 |
| | PenStrep | 1:100 | Sigma Merck | P4333 |

**Supplementary Table 6. Antibodies**

| Antigen name | Host | Diluition | Source | Stock Number | RRID |
|---|---|---|---|---|---|
| **Primary Antibodies** | | | | | |
| **CD49f** | Rat | 1:100 | Biolegend | 313602 | AB_345296 |
| **GFAP** | Mouse | 1:400 | Sigma | MAB360 | AB_11212597 |
| **AQP4** | Rabbit | 1:1000 | Sigma | HPA014784 | AB_1844967 |
| **LMNB1** | Rabbit | 1:1000 | Abcam | AB16048 | AB_443298 |
| **MBP** | Mouse | 1:1000 | Biolegend | 808402 | AB_2564742 |
| **CALB** | Rabbit | 1:1000 | Swant | CB-38a | AB_3107026 |
| **GFP** | Chicken | 1:1000 | IBA Lifesciences | GFP-1020 | AB_10000240 |
| **Secondary Antibodies** | | | | | |
| **AlexaFluor488 Anti-Rt** | Donkey | 1:400 | Jackson Immunoresearch | 712-545-153 | AB_2340684 |
| **Cy3 Anti-Ms** | Donkey | 1:800 | Jackson Immunoresearch | 715-165-151 | AB_2315777 |
| **AlexaFluor647 Anti-Ms** | Donkey | 1:400 | Jackson Immunoresearch | 715-605-151 | AB_2340863 |
| **AlexaFluor647 Anti-Rb** | Donkey | 1:800 | Jackson Immunoresearch | 711-605-152 | AB_2492288 |
| **Cy3 Anti-Rb** | Donkey | 1:400 | Jackson Immunoresearch | 711-165-152 | AB_2307443 |
| **AlexaFluor488 Anti-Chk** | Donkey | 1:400 | Jackson Immunoresearch | 712-545-155 | AB_2340375 |

**Supplementary Table 7. RTqPCR assays and primers**

| Gene | Assay |
|---|---|
| *LMNB1* | Hs01059210_m1 |
| *SOX2* | Hs04234836_s1 |
| *OCT4/POU5F1* | Hs04260367_gH |
| *PAX6* | Hs00240871_m1 |
| *OLIG2* | Hs00380164_s1 |
| *NKX2-2* | Hs00159616_m1 |
| *HMBS* | Hs00609297_m1 |
| *GAPDH* | Hs99999905_m1 |
| *18S* | 4332641 |

| Gene | Fw primer | Rv primer |
|---|---|---|
| *SHMT2* | GAGACCGAAGTGCCATCACA | TCCTCACGGAACTGTCGAGA |
| *PTPRT* | CTGGAGGCTGGTGTTCGATT | CTACCTGGGCAGTGTCCATC |
| *GRM5* | GGAAGAAGGCTTGGTACGCT | ACTATGGGCTTCTTGGAGCG |
| *ACTB* | TCAACACCCCAGCCATGTAC | ATCACGATGCCAGTGGTACG |
