## Supplementary Table 8 for "Functional validation of allele-specific LMNB1 silencing in patient-derived astrocytes as a therapeutic option for Autosomal Dominant Leukodystrophy"

**Supplementary table 8 A. scRNA-seq**

| gene | cluster | p_val | avg_log2FC |
|---|---|---|---|
| SUB1 | scramble | 7,50E-06 | 1,357990156 |
| GPM6B | scramble | 7,94E-06 | 1,793873959 |
| MAPKAPK5 | scramble | 6,64E-05 | 2,736965594 |
| ARPP19 | scramble | 0,000408476 | 1,087007732 |
| SERINC1 | scramble | 0,000490876 | 1,831768445 |
| SPAG9 | scramble | 0,000520394 | 2,456648283 |
| LCMT2 | scramble | 0,000887297 | 6,139551352 |
| TOP2B | scramble | 0,000942043 | 1,420049298 |
| YWHAE | scramble | 0,001029142 | 0,615001327 |
| BMPR1B | scramble | 0,001131647 | 1,446134796 |
| TSPAN3 | scramble | 0,001691842 | 1,024874669 |
| PPP1CB | scramble | 0,002989306 | 1,338009452 |
| SORL1 | scramble | 0,003430753 | 1,866362789 |
| AUTS2 | scramble | 0,005298757 | 2,078501974 |
| CXXC4 | scramble | 0,005325536 | 0,253756592 |
| EAF2 | scramble | 0,005478296 | 2,245306813 |
| CASC9 | scramble | 0,005709991 | 1,634431177 |
| PFN2 | scramble | 0,00626258 | 1,265458007 |
| EIF3J-DT | scramble | 0,007542146 | 3,556948125 |
| AL512646.1 | scramble | 0,00764816 | 1,706762188 |
| SPARC | scramble | 0,008315234 | 0,990962292 |
| LMNB1 | scramble | 0,008679954 | 1,769075534 |
| ZBED5 | scramble | 0,008839981 | 2,166672024 |
| ZFP36L1 | scramble | 0,008869779 | 1,076498137 |
| FXYD6 | scramble | 0,009231099 | 1,057844075 |
| DYNLRB1 | shT4 | 3,73E-10 | 2,262032186 |
| ANXA1 | shT4 | 6,29E-07 | 1,452643183 |
| CRYAB | shT4 | 8,89E-06 | 3,336854639 |
| AKR1C3 | shT4 | 2,55E-05 | 2,839327312 |
| ANXA2 | shT4 | 3,19E-05 | 1,533897541 |
| IFITM3 | shT4 | 4,99E-05 | 1,040363201 |
| NSUN2 | shT4 | 7,20E-05 | 4,309855263 |
| HAX1 | shT4 | 7,40E-05 | 1,953661285 |
| DRC1 | shT4 | 0,000102479 | 3,350497247 |
| PCID2 | shT4 | 0,000111381 | 1,596430024 |
| CMTM3 | shT4 | 0,000115288 | 2,044394119 |
| PSMA3 | shT4 | 0,000119051 | 0,758929221 |
| LSM10 | shT4 | 0,000141928 | 2,133605561 |
| PTK2 | shT4 | 0,000177261 | 1,380975663 |
| CALD1 | shT4 | 0,000204146 | 1,164714145 |
| S100A10 | shT4 | 0,000206632 | 0,825980241 |
| ANXA2P2 | shT4 | 0,000227439 | 2,157541277 |
| TMED2 | shT4 | 0,000263806 | 1,274047399 |
| SELENOM | shT4 | 0,000277443 | 1,053077052 |
| POLR2I | shT4 | 0,000287532 | 1,191790148 |
| SLC47A2 | shT4 | 0,000289444 | 1,613159393 |

| | | | |
|---|---|---|---|
| YWHAG | shT4 | 0,000363203 | 1,272544625 |
| TWSG1 | shT4 | 0,000363605 | 1,870606478 |
| RPLP1 | shT4 | 0,00037387 | 1,04696066 |
| EIF3K | shT4 | 0,00037512 | 1,075749421 |
| C9orf116 | shT4 | 0,000393323 | 4,754887502 |
| SQLE | shT4 | 0,000397631 | 1,465663572 |
| RNF181 | shT4 | 0,000416076 | 1,576335323 |
| FAM114A1 | shT4 | 0,000432022 | 2,244307198 |
| FGF2 | shT4 | 0,000441988 | 3,256339753 |
| COPZ1 | shT4 | 0,00044948 | 1,470814956 |
| GTF2H2 | shT4 | 0,000454755 | 3,131244533 |
| VPS45 | shT4 | 0,000467324 | 1,78181983 |
| NUPR1 | shT4 | 0,000469889 | 1,396047046 |
| SUMO3 | shT4 | 0,000491205 | 1,487445995 |
| ATP5MC2 | shT4 | 0,000495444 | 1,041411495 |
| CALCOCO2 | shT4 | 0,000522938 | 1,964463843 |
| CCN1 | shT4 | 0,000541433 | 1,52702266 |
| MIR4435-2HG | shT4 | 0,000557076 | 1,857980995 |
| TVP23B | shT4 | 0,000593292 | 1,398549376 |
| COX7A2L | shT4 | 0,000594469 | 0,840999961 |
| RIC1 | shT4 | 0,000620386 | 3,896164189 |
| RPL24P4 | shT4 | 0,000650605 | 1,671377253 |
| LAMC1 | shT4 | 0,000678804 | 1,508611615 |
| CNN3 | shT4 | 0,00068329 | 0,986814577 |
| CSTB | shT4 | 0,000724581 | 1,312857104 |
| PEAK1 | shT4 | 0,000747979 | 2,406816992 |
| BOLA3 | shT4 | 0,000770042 | 0,94705112 |
| ADAR | shT4 | 0,000785811 | 2,436639754 |
| SLFN5 | shT4 | 0,000790579 | 4,201633861 |
| BRD3OS | shT4 | 0,000900742 | 1,942284502 |
| PSMB8 | shT4 | 0,000910176 | 3,633461018 |
| A2M | shT4 | 0,000925196 | 2,596573791 |
| PRDX5 | shT4 | 0,000949162 | 0,976815213 |
| H2AFJ | shT4 | 0,00095687 | 3,047305715 |
| SYNE1 | shT4 | 0,000997397 | 2,565199246 |
| EIF4G3 | shT4 | 0,001013085 | 0,58405657 |
| ENY2 | shT4 | 0,001018968 | 1,135376775 |
| PGK1 | shT4 | 0,001045649 | 1,044969447 |
| VIM | shT4 | 0,00105089 | 0,748443845 |
| HSD17B10 | shT4 | 0,001055006 | 2,187627003 |
| DHCR24 | shT4 | 0,001101958 | 2,290358016 |
| SMCO4 | shT4 | 0,001149188 | 2,402964667 |
| ERRFI1 | shT4 | 0,001193616 | 6,624490865 |
| DLC1 | shT4 | 0,001341786 | 4,847996907 |
| MYLK | shT4 | 0,001351989 | 2,728060041 |
| NF2 | shT4 | 0,001378891 | 3,328781766 |
| PARK7 | shT4 | 0,001397818 | 0,576615858 |
| SLC25A3 | shT4 | 0,001398552 | 0,814150484 |

| | | | |
|---|---|---|---|
| PYGO1 | shT4 | 0,001455763 | 3,115477217 |
| RPSAP58 | shT4 | 0,001462442 | 1,060754031 |
| MYL12A | shT4 | 0,001468143 | 1,330916878 |
| VTI1B | shT4 | 0,001600603 | 1,263034406 |
| ATP5PO | shT4 | 0,001684902 | 0,807952188 |
| LATS2 | shT4 | 0,001721248 | 4,922832139 |
| IFITM2 | shT4 | 0,001752981 | 0,191586375 |
| CYSTM1 | shT4 | 0,001802125 | 1,131148765 |
| KATNAL1 | shT4 | 0,001807063 | 1,084316286 |
| CTSA | shT4 | 0,0018827 | 1,633394407 |
| SLC16A2 | shT4 | 0,001882708 | 4,222392421 |
| FTH1P23 | shT4 | 0,002030112 | 1,736965594 |
| WDR37 | shT4 | 0,002096899 | 4,196397213 |
| AREL1 | shT4 | 0,002240731 | 3,516575526 |
| TRAPPC2L | shT4 | 0,002349193 | 0,258241395 |
| ARNT2 | shT4 | 0,002396564 | 2,460680165 |
| HLA-B | shT4 | 0,002458404 | 0,888264762 |
| C1QBP | shT4 | 0,002520594 | 0,896865085 |
| SON | shT4 | 0,002614343 | 0,340183505 |
| TUBB4B | shT4 | 0,002766993 | 1,682530897 |
| GNG10 | shT4 | 0,002786645 | 1,280107919 |
| DRG1 | shT4 | 0,002858883 | 0,987927168 |
| SLIRP | shT4 | 0,002979715 | 1,024767905 |
| CRAT | shT4 | 0,003129814 | 1,493040011 |
| DGUOK | shT4 | 0,003133732 | 1,163418473 |
| TEX9 | shT4 | 0,003157499 | 1,018615678 |
| SLC7A1 | shT4 | 0,00328546 | 3,19401062 |
| RPLP0P6 | shT4 | 0,003310339 | 0,931052646 |
| CLIC4 | shT4 | 0,003312223 | 1,179056013 |
| NRBF2 | shT4 | 0,003342722 | 1,251538767 |
| BUD31 | shT4 | 0,003447129 | 0,646501826 |
| CCDC113 | shT4 | 0,003474083 | 4,258016331 |
| CYTOR | shT4 | 0,003545807 | 2,460841189 |
| LITAF | shT4 | 0,003779629 | 1,269839584 |
| RARS2 | shT4 | 0,003795864 | 0,312189092 |
| IQCE | shT4 | 0,003964474 | 3,273018494 |
| SRSF10 | shT4 | 0,004001333 | 1,891912032 |
| ECM2 | shT4 | 0,004110381 | 3,568842835 |
| TXNRD1 | shT4 | 0,004116547 | 0,386468347 |
| MRPL13 | shT4 | 0,004168261 | 0,869716214 |
| OPTN | shT4 | 0,004176158 | 1,273483355 |
| MLH3 | shT4 | 0,004181973 | 4,297680549 |
| HDDC3 | shT4 | 0,004243446 | 3,906890596 |
| MAP3K19 | shT4 | 0,004247084 | 4,64385619 |
| AC016739.1 | shT4 | 0,004262429 | 2,058893689 |
| AP2S1 | shT4 | 0,004417413 | 0,716207034 |
| CNIH4 | shT4 | 0,004508473 | 1,4665678 |
| OSBPL1A | shT4 | 0,004548997 | 1,273018494 |

| | | | |
|---|---|---|---|
| PEX3 | shT4 | 0,00455074 | 1,5334322 |
| KDELR3 | shT4 | 0,004592926 | 3,191141487 |
| MRGBP | shT4 | 0,00464175 | 4,115477217 |
| SLC25A24 | shT4 | 0,004646536 | 5,058893689 |
| PSENEN | shT4 | 0,004656023 | 0,976636007 |
| MSRB3 | shT4 | 0,00468015 | 3,584962501 |
| ZSWIM6 | shT4 | 0,004684013 | 4,169925001 |
| DTD1 | shT4 | 0,004696601 | 1,116441908 |
| WARS | shT4 | 0,004700043 | 2,105048554 |
| PIGH | shT4 | 0,004808196 | 0,716501492 |
| MYOF | shT4 | 0,004815263 | 2,046012398 |
| AKR1B1 | shT4 | 0,004854218 | 1,334056926 |
| CERK | shT4 | 0,005052629 | 2,679378103 |
| ARL16 | shT4 | 0,005056128 | 0,852101305 |
| PHF11 | shT4 | 0,005137227 | 3,795859283 |
| BAD | shT4 | 0,005165843 | 0,925652659 |
| LRIG3 | shT4 | 0,005250904 | 3,688055994 |
| PCCB | shT4 | 0,005282278 | 0,454565863 |
| DAG1 | shT4 | 0,005303727 | 1,579944426 |
| FDPS | shT4 | 0,005338954 | 1,520776753 |
| FAM32A | shT4 | 0,005440662 | 1,361532169 |
| ARID5B | shT4 | 0,005463192 | 1,819315652 |
| TOGARAM1 | shT4 | 0,005477193 | 2,342686655 |
| CCDC47 | shT4 | 0,005568657 | 1 |
| ARSA | shT4 | 0,005575018 | 3,133855747 |
| SERGEF | shT4 | 0,005924644 | 2,251538767 |
| GMPR2 | shT4 | 0,006119541 | 1,498453507 |
| CWC22 | shT4 | 0,006126855 | 1,119794513 |
| SF3A3 | shT4 | 0,006224932 | 1,340340132 |
| FIS1 | shT4 | 0,006289775 | 1,097492962 |
| NDRG4 | shT4 | 0,006308236 | 1,514573173 |
| H2AFV | shT4 | 0,006319888 | 1,21919354 |
| KIF1B | shT4 | 0,006372595 | 1,232173442 |
| NDUFA11 | shT4 | 0,006392765 | 0,631355406 |
| TRAPPC4 | shT4 | 0,006416618 | 1,179579338 |
| BLVRA | shT4 | 0,006581231 | 1,169212383 |
| SPACA9 | shT4 | 0,006609143 | 0,899237023 |
| PLEKHA7 | shT4 | 0,006654955 | 4 |
| UQCRC1 | shT4 | 0,006655318 | 1,302365829 |
| TDG | shT4 | 0,006664314 | 3,078002512 |
| PPP1R14C | shT4 | 0,006712874 | 1,462343214 |
| DHX57 | shT4 | 0,0067597 | 4,437405312 |
| RCAN1 | shT4 | 0,006790065 | 2,383292858 |
| CARD19 | shT4 | 0,006846337 | 1,131911676 |
| IGF2BP3 | shT4 | 0,006860299 | 3,36923381 |
| DNAJA2 | shT4 | 0,006893964 | 1,152003093 |
| GOT2 | shT4 | 0,006899821 | 1,227410496 |
| IFT43 | shT4 | 0,006959472 | 1,336563441 |

| | | | |
|---|---|---|---|
| PTGR1 | shT4 | 0,007009092 | 3,516575526 |
| TLCD3A | shT4 | 0,00704342 | 2,584962501 |
| DDX50 | shT4 | 0,00706015 | 0,899367637 |
| MTCH2 | shT4 | 0,007068201 | 1,210217707 |
| ALDH4A1 | shT4 | 0,007090838 | 3,600904045 |
| MDH1 | shT4 | 0,00718092 | 0,647204917 |
| ATL3 | shT4 | 0,007224622 | 1,767419391 |
| INPPL1 | shT4 | 0,007225808 | 3,736965594 |
| PARP8 | shT4 | 0,007228969 | 4,502500341 |
| OPA3 | shT4 | 0,00723424 | 3,938599455 |
| ABI3BP | shT4 | 0,007235822 | 5,754887502 |
| SYNC | shT4 | 0,007235822 | 4,502500341 |
| ZNF710-AS1 | shT4 | 0,007235822 | 4,502500341 |
| ZNF225 | shT4 | 0,007236876 | 4,415037499 |
| CAD | shT4 | 0,007237404 | 5,029747343 |
| Z95115.1 | shT4 | 0,007238458 | 6,073248982 |
| AC090114.1 | shT4 | 0,007241623 | 2,874469118 |
| COL11A1 | shT4 | 0,007262253 | 1,497662556 |
| OGDH | shT4 | 0,00727857 | 2,212303604 |
| CTTN | shT4 | 0,007306764 | 0,950531324 |
| ACLY | shT4 | 0,007342382 | 1,46994584 |
| SBF2 | shT4 | 0,007400286 | 1,219009782 |
| LIN7B | shT4 | 0,007641263 | 3,678071905 |
| PIGX | shT4 | 0,007642032 | 1,032950422 |
| GAPDH | shT4 | 0,007733853 | 0,629564471 |
| CCN2 | shT4 | 0,007801619 | 2,499353785 |
| CEBPZOS | shT4 | 0,007820628 | 0,969626351 |
| SAT1 | shT4 | 0,007944292 | 0,609986725 |
| FSIP1 | shT4 | 0,007958274 | 3,91753784 |
| DNPH1 | shT4 | 0,008089676 | 0,923501863 |
| ACOT8 | shT4 | 0,00823323 | 3,089637212 |
| TIMM50 | shT4 | 0,008388845 | 1,287866259 |
| WDR77 | shT4 | 0,008406294 | 3,152003093 |
| RPS27AP12 | shT4 | 0,008416236 | 0,366943211 |
| MRPL34 | shT4 | 0,008495511 | 1,419538892 |
| RPL23AP42 | shT4 | 0,008503445 | 0,579424317 |
| YAP1 | shT4 | 0,008548473 | 1,668794092 |
| OBSL1 | shT4 | 0,008572664 | 2,470319935 |
| THUMPD3 | shT4 | 0,008599012 | 1,370837695 |
| EXOC5 | shT4 | 0,008671925 | 0,361991265 |
| TIMM10 | shT4 | 0,008785623 | 0,376563351 |
| NSDHL | shT4 | 0,008787765 | 2,313540309 |
| CSNK2B | shT4 | 0,00883358 | 0,921671639 |
| B3GALNT2 | shT4 | 0,009070336 | 3,378511623 |
| P4HA2 | shT4 | 0,009171723 | 1,263034406 |
| PHF20L1 | shT4 | 0,009238463 | 1,502500341 |
| AP2B1 | shT4 | 0,009291501 | 0,694497453 |
| GPX8 | shT4 | 0,009309266 | 1,612835678 |

| | | | |
|---|---|---|---|
| MRPS7 | shT4 | 0,009364175 | 0,57022887 |
| KPNA6 | shT4 | 0,009381635 | 1,767553914 |
| ANKRD12 | shT4 | 0,009484597 | 1,218729207 |
| C8orf34 | shT4 | 0,0094941 | 1,238159737 |
| SMARCE1 | shT4 | 0,009579543 | 0,693258309 |
| ZDHHC17 | shT4 | 0,009716093 | 1,227535725 |
| RBPMS | shT4 | 0,009716317 | 3,42786154 |

**Supplementary Table 8 B. GO Analysis - Biological Process**

| ID | Description | GeneRatio | BgRatio | pvalue | p.adjust | qvalue | geneID |
|---|---|---|---|---|---|---|---|
| GO:0052548 | regulation of endopeptidase activity | 17/191 | 428/18903 | 1,73E-06 | 0,005070467 | 0,00466059 | CRYAB/ANXA2/PCID2/PSMA3/CCN1/CSTB/PSMB8/A2M/PRDX5/DHCR24/DLC1/PARK7/PSENEN/BAD/FIS1/GAPDH/CCN2 |
| GO:0052547 | regulation of peptidase activity | 17/191 | 459/18903 | 4,43E-06 | 0,006476952 | 0,005953381 | CRYAB/ANXA2/PCID2/PSMA3/CCN1/CSTB/PSMB8/A2M/PRDX5/DHCR24/DLC1/PARK7/PSENEN/BAD/FIS1/GAPDH/CCN2 |
| GO:0006694 | steroid biosynthetic process | 10/191 | 178/18903 | 1,37E-05 | 0,013403416 | 0,012319937 | AKR1C3/SQLE/HSD17B10/DHCR24/OSBPL1A/AKR1B1/FDPS/ACLY/ACOT8/NSDHL |
| GO:1901617 | organic hydroxy compound biosynthetic process | 11/191 | 245/18903 | 4,12E-05 | 0,027916038 | 0,025659415 | AKR1C3/SQLE/FGF2/HSD17B10/DHCR24/PARK7/OSBPL1A/FDPS/ACLY/ACOT8/NSDHL |
| GO:0009199 | ribonucleoside triphosphate metabolic process | 11/191 | 249/18903 | 4,77E-05 | 0,027916038 | 0,025659415 | NUPR1/ATP5MC2/PGK1/ATP5PO/DGUOK/BAD/FIS1/NDUFA11/CAD/OGDH/GAPDH |
| GO:0046034 | ATP metabolic process | 10/191 | 217/18903 | 7,44E-05 | 0,032888416 | 0,030229846 | NUPR1/ATP5MC2/PGK1/ATP5PO/DGUOK/BAD/FIS1/NDUFA11/OGDH/GAPDH |
| GO:0009141 | nucleoside triphosphate metabolic process | 11/191 | 267/18903 | 8,92E-05 | 0,032888416 | 0,030229846 | NUPR1/ATP5MC2/PGK1/ATP5PO/DGUOK/BAD/FIS1/NDUFA11/CAD/OGDH/GAPDH |
| GO:0010810 | regulation of cell-substrate adhesion | 10/191 | 222/18903 | 8,99E-05 | 0,032888416 | 0,030229846 | PTK2/S100A10/CCN1/PEAK1/DLC1/NF2/C1QBP/ECM2/DAG1/ABI3BP |
| GO:0031589 | cell-substrate adhesion | 13/191 | 369/18903 | 1,01E-04 | 0,032902677 | 0,030242955 | PTK2/S100A10/CCN1/LAMC1/PEAK1/DLC1/NF2/C1QBP/ECM2/DAG1/ABI3BP/CTTN/CCN2 |
| GO:2000116 | regulation of cysteine-type endopeptidase activity | 10/191 | 235/18903 | 1,44E-04 | 0,039305539 | 0,036128234 | CRYAB/PCID2/CCN1/PRDX5/DHCR24/DLC1/PARK7/BAD/FIS1/CCN2 |
| GO:0031639 | plasminogen activation | 4/191 | 27/18903 | 1,48E-04 | 0,039305539 | 0,036128234 | ANXA2/S100A10/PGK1/DHCR24 |
| GO:0009060 | aerobic respiration | 9/191 | 194/18903 | 1,62E-04 | 0,039591265 | 0,036390863 | NUPR1/COX7A2L/PARK7/ATP5PO/DGUOK/NDUFA11/UQCRC1/MDH1/OGDH |
| GO:0009205 | purine ribonucleoside triphosphate metabolic process | 10/191 | 242/18903 | 1,82E-04 | 0,041016017 | 0,037700443 | NUPR1/ATP5MC2/PGK1/ATP5PO/DGUOK/BAD/FIS1/NDUFA11/OGDH/GAPDH |
| GO:0009144 | purine nucleoside triphosphate metabolic process | 10/191 | 247/18903 | 2,15E-04 | 0,044920088 | 0,041288925 | NUPR1/ATP5MC2/PGK1/ATP5PO/DGUOK/BAD/FIS1/NDUFA11/OGDH/GAPDH |
| GO:0043281 | regulation of cysteine-type endopeptidase activity involved in apoptotic process | 9/191 | 205/18903 | 2,45E-04 | 0,047812871 | 0,043947867 | CRYAB/CCN1/PRDX5/DHCR24/DLC1/PARK7/BAD/FIS1/CCN2 |
