## Supplementary Table 9 for "Functional validation of allele-specific LMNB1 silencing in patient-derived astrocytes as a therapeutic option for Autosomal Dominant Leukodystrophy"

### Supplementary Table 9. Statistics

| Figure | Applied Test | n | P value | Statistics | Post hoc analyses | Post hoc results |
|---|---|---|---|---|---|---|
| **Fig. 1D** | Ordinary one-way ANOVA | CTRL=3<br>IT3.1=2<br>IT3.2=4<br>IT3.3=2 | p<0.01 | F=16.87 | Tukey's multiple comparisons test | CTRL vs IT3.1 = p<0.01<br><br>CTRL vs IT3.2 = p<0.01<br><br>CTRL vs IT3.3 = p<0.05 |
| **Fig. 1E** | Ordinary one-way ANOVA | CTRL=8<br>IT3.1=2<br>IT3.2=3<br>IT3.3=3 | p<0.0001 | F=20.88 | Tukey's multiple comparisons test | CTRL vs IT3.1 = p<0.001<br><br>CTRL vs IT3.2 = p<0.05<br><br>CTRL vs IT3.3 = p<0.001 |
| **Fig. 1F** | Chi-square test | CTRL=3 cell lines (2-4 independent experiments/cell line)<br>ADLD=3 cell lines (2-4 independent experiments/cell line)<br>CTRL=259 cells<br>ADLD=179 cells | p<0.0001 | $\chi^2$=70.38 | | |
| **Fig. 1G** | Fisher's exact test | CTRL=3 cell lines (2-4 independent experiments/cell line)<br>ADLD=3 cell lines (2-4 independent experiments/cell line)<br>CTRL=259 cells<br>ADLD=179 cells | p<0.0001 | | | |
| **Fig. 3C** | Two-way RM ANOVA | CTRL ACM=2 cell lines (2 independent | <u>TimeXGenotype</u><br>p<0.0001 | <u>TimeXGenotype</u><br>F=119.2<br><u>Time</u> F=1302 | Sidak's multiple | <u>6h</u><br>CTRL vs IT3.1 = ns<br>CTRL vs IT3.2 = ns<br>CTRL vs IT3.3 = ns |

| | | | | | | |
|---|---|---|---|---|---|---|
| | | experiments/cell line)<br>ADLD ACM=3 cell lines (2 independent experiments/cell line) | Time p<0.0001<br>Genotype p<0.0001<br>Experiment ns | Genotype F=596.3<br>Experiment F=0.19 | comparisons test | 16h<br>CTRL vs IT3.1 = p<0.05<br>CTRL vs IT3.2 = ns<br>CTRL vs IT3.3 = p<0.05 |
| **Fig. 3D** | Two-way RM ANOVA | CTRL ACM=2 cell lines (2 independent experiments/cell line)<br>ADLD ACM=3 cell lines (2 independent experiments/cell line) | TimeXGenotype p<0.05<br>Time p<0.0001<br>Genotype p<0.05<br>Experiment ns | TimeXGenotype F=3.23<br>Time F=246.1<br>Genotype F=8.95<br>Experiment F=1.32 | Sidak's multiple comparisons test | 4h<br>CTRL vs IT3.1 = ns<br>CTRL vs IT3.2 = ns<br>CTRL vs IT3.3 = ns<br><br>8h<br>CTRL vs IT3.1 = ns<br>CTRL vs IT3.2 = ns<br>CTRL vs IT3.3 = ns<br><br>12h<br>CTRL vs IT3.1 = p<0.05<br>CTRL vs IT3.2 = ns<br>CTRL vs IT3.3 = p<0.01 |
| **Fig. 3E** | Ordinary one-way ANOVA | CTRL SCR ACM=6 independent experiments, 2 cell lines<br>IT3.1 SCR ACM=2 independent experiments<br>IT3.2 SCR ACM=2 independent experiments<br>IT3.3 SCR ACM=2 independent experiments<br>IT3.1 shRNA ACM=2 | p<0.0001 | F=12.74 | Tukey's multiple comparisons test | CTRL SCR ACM vs IT3.1 SCR ACM = p<0.001<br><br>CTRL SCR ACM vs IT3.2 SCR ACM = p<0.0001<br><br>CTRL SCR ACM vs IT3.3 SCR ACM = p<0.0001<br><br>CTRL SCR ACM vs IT3.1 shRNA ACM = ns<br><br>CTRL SCR ACM vs IT3.2 |

| | | | | | | |
|---|---|---|---|---|---|---|
| | | independent experiments<br>IT3.2 shRNA ACM=2 independent experiments<br>IT3.3 shRNA ACM=2 independent experiments | | | | shRNA ACM = ns<br><br>CTRL SCR ACM vs IT3.3 shRNA ACM = ns<br><br>IT3.1 SCR ACM vs IT3.1 shRNA ACM = p<0.01<br><br>IT3.2 SCR ACM vs IT3.2 shRNA ACM = p<0.01<br><br>IT3.3 SCR ACM vs IT3.3 shRNA ACM = p<0.001 |
| **Fig. 3F** | Ordinary one-way ANOVA | CTRL SCR ACM=6 independent experiments, 2 cell lines<br>IT3.1 SCR ACM=2 independent experiments<br>IT3.2 SCR ACM=2 independent experiments<br>IT3.3 SCR ACM=2 independent experiments<br>IT3.1 shRNA ACM=4 independent experiments<br>IT3.2 shRNA ACM=4 independent experiments<br>IT3.3 shRNA ACM=4 independent experiments | p<0.0001 | F=1.929 | Tukey's multiple comparisons test | CTRL SCR ACM vs IT3.1 SCR ACM = p<0.01<br><br>CTRL SCR ACM vs IT3.2 SCR ACM = ns<br><br>CTRL SCR ACM vs IT3.3 SCR ACM = p<0.0001<br><br>CTRL SCR ACM vs IT3.1 shRNA ACM = ns<br><br>CTRL SCR ACM vs IT3.2 shRNA ACM = ns<br><br>CTRL SCR ACM vs IT3.3 shRNA ACM = ns<br><br>IT3.1 SCR ACM vs IT3.1 shRNA ACM = p<0.05 |

| | | | | | | |
|---|---|---|---|---|---|---|
| | | | | | | IT3.2 SCR ACM vs IT3.2 shRNA ACM = ns<br><br>IT3.3 SCR ACM vs IT3.3 shRNA ACM = p<0.0001 |
| **Fig. 3G** | Ordinary one-way ANOVA | IT3.1 shRNA ACM=4 independent experiments<br>IT3.2 shRNA ACM=4 independent experiments<br>IT3.3 shRNA ACM=4 independent experiments | p<0.001 | F=1.433 | Tukey's multiple comparisons test | IT3.1 shRNA ACM vs IT3.2 shRNA ACM = ns<br><br>IT3.1 shRNA ACM vs IT3.3 shRNA ACM = p<0.01<br><br>IT3.2 shRNA ACM vs IT3.3 shRNA ACM = p<0.001 |
| **Fig. 4B** | RM one-way ANOVA | SR=3 independent experiments<br>CTRL SCR ACM=3 independent experiments<br>IT3.3 SCR ACM=3 independent experiments<br>IT3.3 shRNA ACM=3 independent experiments | p<0.05 | F=8.93 | Tukey's multiple comparisons test | SR vs CTRL SCR ACM = ns<br><br>SR vs IT3.3 SCR ACM = p<0.01<br><br>SR vs IT3.3 shRNA ACM = ns<br><br>CTRL SCR ACM vs IT3.3 SCR ACM = p<0.01<br><br>CTRL SCR ACM vs IT3.3 shRNA ACM = ns<br><br>ADLD SCR ACM vs IT3.3 shRNA ACM = p<0.05 |

| | | | | | | |
|---|---|---|---|---|---|---|
| **Fig. 5B** | RM one-way ANOVA | SR=3 independent experiments<br>IT3.3 SCR=3 independent experiments<br>IT3.3 shRNA=3 independent experiments | p<0.01 | F=47.78 | Tukey's multiple comparisons test | SR vs IT3.3 SCR = p<0.05<br><br>SR vs IT3.3 shRNA = ns<br><br>IT3.3 SCR vs IT3.3 shRNA = p<0.05 |
| **Suppl. Fig. 1E** | Two-way RM ANOVA | CTRL=3 cell lines (3-4 independent experiments/cell line)<br>ADLD=3 cell lines (3-4 independent experiments/cell line) | *PAX6*<br>TimeXGenotype ns<br>Time p<0.05<br>Genotype ns<br>Experiment ns<br><br>*OLIG2*<br>TimeXGenotype ns<br>Time p<0.05<br>Genotype ns<br>Experiment ns<br><br>*NKX2-2*<br>TimeXGenotype ns<br>Time p<0.05<br>Genotype ns<br>Experiment ns | *PAX6*<br>TimeXGenotype F=2.56<br>Time F=4.71<br>Genotype F=3.84<br>Experiment F=1.33<br><br>*OLIG2*<br>TimeXGenotype F=0.52<br>Time F=6.79<br>Genotype F=0.037<br>Experiment F=1.56<br><br>*NKX2-2*<br>TimeXGenotype F=1.90<br>Time F=7.62<br>Genotype F=1.99<br>Experiment F=1.05 | Sidak's multiple comparisons test | *PAX6*<br>CTRL vs ADLD DIV0 = ns<br>CTRL vs ADLD DIV8 = ns<br>CTRL vs ADLD DIV12 = ns<br><br>*OLIG2*<br>CTRL vs ADLD DIV0 = ns<br>CTRL vs ADLD DIV8 = ns<br>CTRL vs ADLD DIV12 = ns<br><br>*NKX2-2*<br>CTRL vs ADLD DIV0 = ns<br>CTRL vs ADLD DIV8 = ns<br>CTRL vs ADLD DIV12 = ns |
| **Suppl. Fig. 1F** | Unpaired T-test (2-tailed) | CTRL=3 cell lines (2-3 experiments/cell line)<br>ADLD=3 cell lines (2-3 experiments/cell line) | ns | | | |
| **Suppl. Fig. 2C** | Unpaired T-test (2-tailed) | CTRL=3 cell lines (2-3 experiments/cell line)<br>ADLD=3 cell lines (2-3 | ns | | | |

| | | | | | | |
|---|---|---|---|---|---|---|
| | | experiments/cell line) | | | | |
| **Suppl. Fig. 2D** | Unpaired T-test (2-tailed) | CTRL=3 cell lines (2-3 experiments/cell line) ADLD=3 cell lines (2-3 experiments/cell line) | ns | | | |
| **Suppl. Fig. 2E** | Unpaired T-test (2-tailed) | CTRL=3 cell lines (2-3 experiments/cell line) ADLD=3 cell lines (2-3 experiments/cell line) | ns | | | |
| **Suppl. Fig. 2F** | Unpaired T-test (2-tailed) | CTRL=3 cell lines (2-3 experiments/cell line) ADLD=3 cell lines (2-3 experiments/cell line) | ns | | | |
| **Suppl. Fig. 2G** | Unpaired T-test (2-tailed) | CTRL=3 cell lines (2 experiments/cell line) ADLD=3 cell lines (2 experiments/cell line) | ns | | | |
| **Suppl. Fig. 3A** | Unpaired T-test (2-tailed) | CTRL=3 cell lines (2 experiments/cell line) ADLD=3 cell lines (2 experiments/cell line) | p<0.05 | | | |
| **Suppl. Fig. 3B** | Unpaired T-test (2-tailed) | CTRL=3 cell lines (2 experiments/cell line) ADLD=3 cell lines (2-3 experiments/cell line) | p<0.01 | | | |
| **Suppl. Fig. 3C** | Unpaired T-test (2-tailed) | CTRL=3 cell lines (2 experiments/cell line) ADLD=3 cell lines (2-3 experiments/cell line) | p<0.05 | | | |
| **Suppl. Fig. 4D** | Ordinary one-way ANOVA | | | | | CTRL vs ADLD UNTREATED = p<0.001 |

| | | | | | | |
|---|---|---|---|---|---|---|
| | | CTRL=2 cell lines (1-2 experiments/cell lines)<br>ADLD UNTREATED=3 cell lines<br>ADLD SCR=3 cell lines<br>ADLD shRNA=3 cell lines | $p<0.001$ | $F=21.43$ | Tukey's multiple comparisons test | CTRL vs ADLD SCR = $p<0.01$<br><br>CTRL vs ADLD SHRNA = ns<br><br>ADLD UNTREATED vs ADLD SCR = ns<br><br>ADLD UNTREATED vs ADLD shRNA = $p<0.01$<br><br>ADLD SCR vs ADLD shRNA = $p<0.05$ |
| **Suppl. Fig. 5A** | RM one-way ANOVA | n=2 independent experiments/time point | ns | $F=13.78$ | Tukey's multiple comparisons test | 16h post LPC vs 72h post LPC = $p<0.05$ |
